# LonGP: an additive Gaussian process regression model for longitudinal study designs

**DOI:** 10.1101/259564

**Authors:** Lu Cheng, Siddharth Ramchandran, Tommi Vatanen, Niina Lietzen, Riitta Lahesmaa, Aki Vehtari, Harri Lähdesmäki

**Affiliations:** Department of Computer Science, Aalto University School of Science, FI-00076 Aalto, Finland.; Broad Institute of MIT and Harvard, Cambridge, MA 02142, USA.; Turku Centre for Biotechnology, University of Turku and Åbo Akademi University, Turku 20520, Finland.

## Abstract

**Motivation:** Biomedical research typically involves longitudinal study designs where samples from individuals are measured repeatedly over time and the goal is to identify risk factors (covariates) that are associated with an outcome value. General linear mixed effect models have become the standard workhorse for statistical analysis of data from longitudinal study designs. However, analysis of longitudinal data can be complicated for both practical and theoretical reasons, including difficulties in modelling, correlated outcome values, functional (time-varying) covariates, nonlinear effects, and model inference.

**Results:** We present LonGP, an additive Gaussian process regression model for analysis of experimental data from longitudinal study designs. LonGP implements a flexible, non-parametric modelling framework that solves commonly faced challenges in longitudinal data analysis. In addition to inheriting all standard features of Gaussian processes, LonGP can model time-varying random effects and non-stationary signals, incorporate multiple kernel learning, and provide interpretable results for the effects of individual covariates and their interactions. We develop an accurate Bayesian inference and model selection method, and implement an efficient model search algorithm for our additive Gaussian process model. We demonstrate LonGP’s performance and accuracy by analysing various simulated and real longitudinal -omics datasets. Our work is accompanied by a versatile software implementation.

**Availability:** LonGP software tool is available at http://research.cs.aalto.fi/csb/software/longp/.

## 1 Introduction

A majority of biomedical research involves longitudinal studies where individuals are followed over a period of time and measurements are repeatedly collected from the subjects of the study. Longitudinal studies are effective in identifying various risk factors that are associated with an outcome, such as disease initiation, disease onset or any disease associated molecular biomarker. Characterisation of such risk factors is essential in understanding disease pathogenesis as well as in assessing individuals’ disease risk, patient stratification, treatment choice evaluation and, in future personalised medicine paradigm, planning disease prevention strategies.

There are several classes of longitudinal study designs, including prospective vs. retrospective studies and observational vs. experimental studies, and each of these can be implemented with a particular application-specific experimental design. As the risk factors (or covariates) can also be either static or time-varying, statistical analysis tools need to be versatile enough so that they can be appropriately tailored to every application. General linear mixed effect models and generalized estimating equations have become popular statistical techniques for longitudinal data analysis (Gibbons *et al*, 2010). Although numerous advanced extensions of these two statistical techniques have been proposed, longitudinal data analysis is still complicated for several reasons, such as difficulties in choosing covariance structures to model correlated outcomes, handling irregular sampling times and missing values, accounting for time-varying covariates, choosing appropriate nonlinear effects, modelling non-stationary signals, and accurate model inference.

Modern statistical methods for longitudinal data analysis make less or better assumptions about the underlying data generating mechanisms. These methods use predominantly non-parametric models, such as splines (Wu and Zhang, 2006), and more recently latent stochastic processes, such as Gaussian processes (GP). Several Bayesian non-parametric methods have been proposed for longitudinal and other data analysis. Most pertinent to this work are recent work on Bayesian semi-parametric models (Quintana *et al*, 2016) and additive GP regression (Qamar and Tokdar, 2014) for longitudinal data analysis. Interestingly, very similar models have been developed in machine learning community. Additive GPs together with type-II maximum likelihood based multiple kernel learning were introduced in (Duvenaud *et al*, 2011). Similar GP multiple kernel learning has also been formulated in terms of hypothesis testing (Liu and Coull, 2017).

We present a non-parametric model, LonGP, for longitudinal data analysis that is formulated as an additive GP which handles commonly faced challenges in longitudinal data analysis. Being a GP model, LonGP inherits the best features of GPs. Additionally, it can model time-varying random effects and non-stationary signals as well as provide interpretable results for the effects of individual covariates and their interactions. We develop a fully Bayesian predictive inference for LonGP and use that to carry out model selection, i.e., to identify covariates that are associated with a given study outcome value. We demonstrate LonGP’s performance and accuracy by analysing various simulated and real longitudinal -omics data sets.

## 2 Methods

### 2.1 Notation

We model target variables (gene/protein/bacteria/etc) one at a time. Let us assume that there are *P* individuals and there are *n*_*i*_ time series measurements from the *i*th individual. The total number of data points is thus 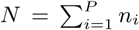. We denote the target variable by a column vector ***y*** *=* (*y*_1_,*y*_2_, … *y*_*N*_)*^T^* and the covariates by *X* = (***x***_1_, ***x***_2_, …, ***x***_N_), where ***x****_i_* = (*x*_*i*1_,*x*_*i*2_, …,*x*_*id*_)*^T^* is a *d*-dimensional column vector and *d* is the number of covariates. We denote the domain of the *j*th variable by 𝒳_*j*_ and the joint domain of all covariates is 𝒳 *=* 𝒳_1_ × 𝒳_2_ × … × 𝒳*_d_.* In general, we use a bold font letter to denote a vector, an uppercase letter to denote a matrix and a lowercase letter to denote a scale value.

### 2.2 Gaussian process

Gaussian process (GP) can be seen as a distribution of nonlinear functions (Rasmussen and Williams, 2006). For inputs ***x, x***′ ∈ 𝒳, GP is defined as

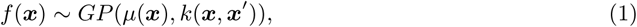

where *µ*(***x***) is the mean and *k*(***x, x***′) is a positive-semidefinite kernel function that defines the co-variance between any two realizations of *f*(***x***) and *f*(***x****’*) by

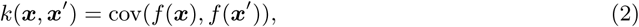

which is called “kernel” for short. The mean is often assumed to be zero, i.e., *µ*(***x***) ≐ 0, and the kernel has parameters ***θ***, i.e., *k*(***x, x****’|****θ***). For any finite collection of inputs *X* = (***x***_1_, ***x***_2_, …, ***x***_*N*_), the function values ***f***(*X*) *=* (*f*(***x***_1_), *f*(***x***_2_), …, *f*(***x***_*N*_))*^T^* have joint multivariate Gaussian distribution

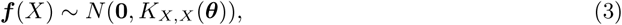

where elements of the *N*-by-*N* covariance matrix are defined by the kernel [*K*_*X,X*_(***θ***)]_*i,j*_ = *k*(***x***_*i*_, ***x****_j_|****θ***).

We use the following hierarchical Gaussian process model

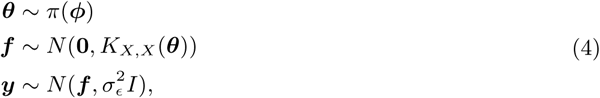

where *π*(*ϕ*) defines a prior for the kernel parameters (including 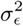), 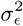 is the noise variance and *I* is the *N*-by-*N* identity matrix. For a Gaussian noise model we can marginalise ***f*** analytically (Rasmussen and Williams, 2006)

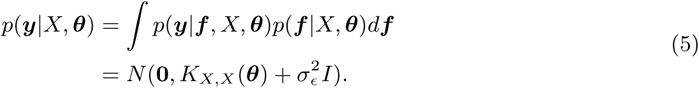

### 2.3 Additive Gaussian process

To define a flexible and interpretable model, we use the following additive GP model with *D* kernels

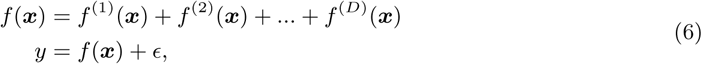

where each *f*^(^*^j^*^)^ (***x***) ~ *GP*(0, *k*^(^*^j^*^)^ (***x, x***′|***θ***^(^*^j^*^)^)) is a separate GP with kernel specific parameters ***θ***^(^*^j^*^)^ and *ϵ* is the additive Gaussian noise. By definition, for any finite collection of inputs *X* = (*x*_1_, *x*_2_, …, *x*_*N*_), each GP ***f***^(^*^j^*^)^(*X*) follows a multivariate Gaussian distribution. Since a sum of multivariate Gaussian random variables is still Gaussian, the latent function ***f*** also follows a multivariate Gaussian distribution. Denote 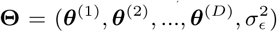, then the marginal likelihood for the target variable ***y*** is

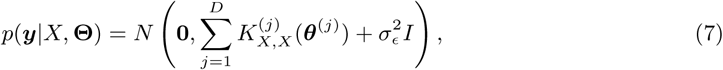

where the latent function ***f*** has been marginalised out as in Eq. (5). To simplify notation, we define

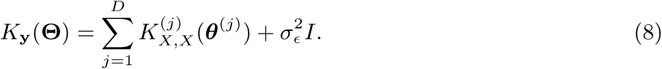

For the purposes of identifying covariate subsets that are associated with a target variable, we assume that each GP depends only on a small subset of covariates *f^(j^*^)^(*x*): 𝒳*^(j)^* → 𝒴, where 𝒳*^(j)^* = Π 𝒳_*i*_,*i* ϵ *I*_*j*_ ⊆ {*1*, …,*d*} and 𝒴 is the domain for target variable. *I*_*j*_ are indices of the covariates associated with the *j*th kernel.

### 2.4 Kernel functions for covariates

Longitudinal biomedical studies typically include a variety of continuous, categorical and binary covariates. Typical continuous covariates include *age, time from a disease event* (sampling time point minus disease event time point), and *season* (time from beginning of a year). Typical categorical or binary covariates include *group* (case or control), *gender* and *id* (id of an individual). In practice, a key question in setting up the additive GP model is how to choose appropriate kernels for different covariates and their subsets (or interactions).

#### 2.4.1 Stationary kernels

In LonGP, we use the following specific kernels which only involve one or two covariates.

- Squared exponential (SE) kernel for continuous covariates

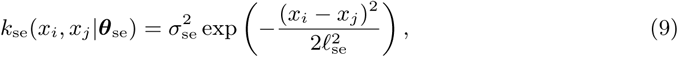

where *ℓ*_se_ is the length-scale parameter, 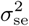 is the magnitude parameter and 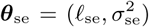. Length-scale *ℓ*_se_ controls the smoothness and magnitude parameter 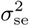 controls the magnitude of the kernel.
- Periodic kernel for continuous covariates

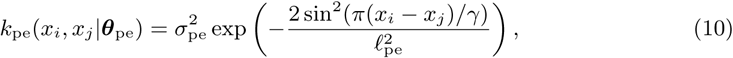

where *ℓ*_se_ is the length-scale parameter, 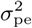 is the magnitude parameter, *γ* is the period parameter and 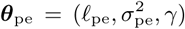. Length-scale *ℓ*_se_ controls the smoothness, 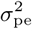 controls the magnitude and *γ* is the period of the kernel. In our model, *γ* corresponds to a year.
- Constant kernel

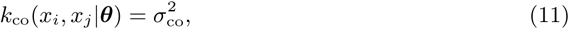

where 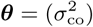 is the magnitude parameter of the constant signal.
- Categorical kernel for discrete-valued covariates

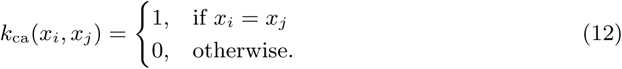
- Binary (mask) kernel for binary covariates

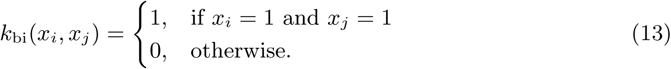
- Product kernel between any two valid kernels, such as *k*_bi_(·) and *k*_se_(·) (similarly for any other pair of kernels)

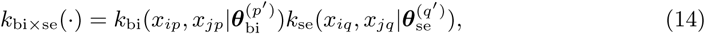

where 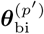 and 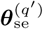 are kernel parameters for the *p*th and *q*th covariates, respectively.

#### 2.4.2 Non-stationary kernel

It may be realistic to assume that the target variable (e.g., a protein) changes rapidly only near a special event, such as disease initiation or onset. This poses a challenge for GP modelling with squared exponential kernel since the kernel is stationary: changes are homogeneous across the whole time window. Non-stationary GPs can be implemented by using special non-stationary kernels, such as the neural network kernel, by defining the kernel parameters to depend on input covariates (Heinonen *et al.*, 2016; Tolvanen *et al.*, 2014; Saul *et al.*, 2016) or via input or output warpings (Snelson *et al*, 2004). We propose to use the input warping approach and define a bijective mapping *ω*: (−∞, +∞) → (−*c, c*) for a continuous time/age covariate *t* as

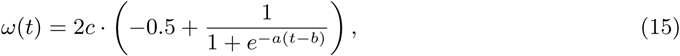

where *a, b* and *c* are predefined parameters: *a* controls the size of the effective time window, *b* controls its location, and *c* controls the maximum range. The non-stationary kernel is then defined as

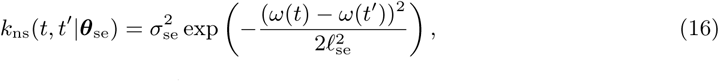

where ***θ***_se_ are the parameters of the SE kernel.

Suppl. Fig. 1 shows an example transformation with *a* = 0.5, *b* = 0 and *c* = 40, where we limit the disease related change to be within one year of the disease event. Effectively, all changes in the transformed space corresponds approximately to ±12 month time window in the original space. Suppl. Fig. 2 shows randomly sampled functions using stationary and non-stationary SE kernels with the same kernel parameters. The non-stationary SE kernel naturally models signals that are spikelike or exhibit a level difference between before and after the disease event, which can be interpreted as a permanent disease effect.

The same parameters as Suppl. Fig. 1 are used for non-stationary kernels in all experiments of Sec. 3.

#### 2.4.3 Kernel specification in practice

The datasets analysed in this work include 11 covariates and covariate pairs which we model using the following kernels (see Sec. 2.5 for prior specifications).

- *age:* The shared age effect is modelled with a slowly changing stationary SE kernel.
- *time from a disease event* or *diseaseAge:* We use the product of the binary kernel and the non-stationary SE kernel (assuming cases are coded as 1 and controls as 0).
- *season:* We assume that the target variable exhibits an annual period and is modelled with the periodic kernel.
- *group*: We model a baseline difference between the cases and controls, which corresponds to average difference between the two groups, using the product of the binary kernel and the constant kernel.
- *gender*: We use the same kernel as for *group* covariate.
- *loc*: Binary covariate indicating if an individual comes from a certain location. We use the same kernel as for group covariate.
- *id*: We assume baseline differences between different individuals and model that by the product of the categorical kernel and the constant kernel.
- *group* × *age*: We assume that the differences between cases and controls varies across age. That difference is modelled by the product of the binary kernel and the stationary SE kernel.
- *gender* × *age*: The same kernel as for *group* × *age* is used for this interaction term. It implements a different age trend for males and females.
- *id* × *age*: We assume different individuals exhibit different age trends. This random effect is modelled by the product of the categorical kernel and the SE kernel. kernel is especially helpful for modelling individuals with outlying data points.
- *group* × *gender*: This interaction term assumes that male (or female) cases have a baseline difference compared to others. The product of two binary kernels and the constant kernel is used.

Although discrete covariates are modelled as a product of the constant kernel and the binary or categorical kernel, the constant kernel is not explicitly included in our notation.

Fig. 1 shows an example with data simulated from an additive GP model, 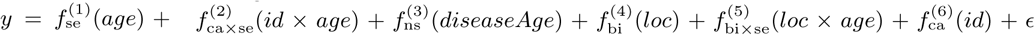. This example provides an intuitive illustration of the effects of different kernels described above. In case a study contains other covariates or interaction terms, the additive Gaussian process regression provides a very flexible modelling framework that can be adjusted to a number of different applications.

**Figure 1:**
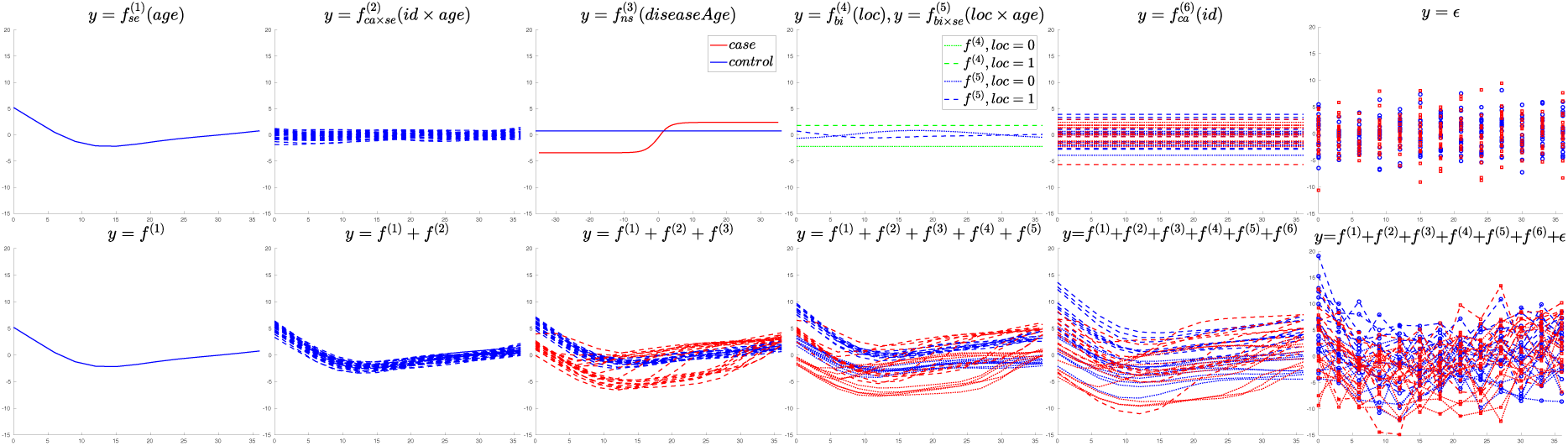
Additive Gaussian process. The top panel shows random functions drawn from different components, i.e., GPs of the specific kernels. The lower panel shows the cumulative effects of the different components. The bottom right panel shows the simulated data.

In practice, we often observe missing values in the covariates. Missing values can be due to technical problems in measurements or because some covariates may not be applicable for certain samples, e.g., *diseaseAge* is not applicable to controls since they do not have a disease. In LonGP, we construct a binary flag vector for each covariate. The missing values are flagged as 0 and non-missing values are flagged as 1. Then, we construct a binary kernel for this flag vector and multiply it with any kernel that involves the covariate. Consequently, any kernel involving a missing value is evaluated to 0, which means that their contribution to the target variable is 0. All missing values are handled in this way by default and we do not use any extra notations for it. Interaction terms always refer to product kernels with non-missing values, assuming missing values are already handled.

### 2.5 Prior specifications

Before the actual GP regression, we standardise the target variable and all continuous covariates such that the mean is zero and the standard deviation is one. This helps in defining generally applicable priors for the kernel parameters. After the GP regression, the predictions are transformed back to the original scale. We visualise the results in the original scale after centering the data by subtracting the mean.

We define a prior 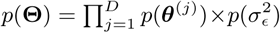 for the kernel parameters as follows. For continuous covariates without interactions, we use the log normal prior (*µ* = 0 and *σ^2^* = (log(1) − log(0.1))^2^/4) for the length-scales (*ℓ*_se_ and *ℓ*_pe_) and the square root student-*t* prior (*µ* = 0, *σ^2^* = 1 and *ν* = 20) for the magnitude parameters (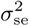 and 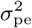). This length-scale prior penalises small length-scales such that smoothness less than 0.1 has very small probability and the mode is approximately at 0.3. For continuous covariates with interactions, the prior for the magnitude parameters is the same as for without interactions and the half truncated student-*t* prior (*µ* = 0, *σ^2^* = 1, *ν* = 4) is used for the length-scale, which allows smaller length-scales.

Scaled inverse chi-squared prior (*σ^2^* = 0.01 and *ν* = 1) is used for the noise variance parameter 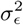. The period parameter *γ* of the periodic kernel is predefined by the user. Square root student-*t* prior (*µ* = 0, *σ^2^* = 1 and *ν* = 4) is used for the magnitude parameter 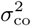 of all constant kernels. Suppl. Fig. 3 visualises all the above-described priors with their default hyperparameter values.

### 2.6 Model inference and prediction

Given the additive GP model specified in Sections 2.2-2.5, we are next interested in the posterior inference of the model conditioned on data (***y, X***). Assume, for now, that for each additive component *f^(j)^* the kernel *k^(j^*^)^(·), its inputs 𝒳*^(j)^* and prior are specified. We use two different inference methods, Markov chain Monte Carlo (MCMC) and a deterministic evaluation of the posterior with the central composite design (CCD).

For MCMC we use the slice sampler as implemented in the GPStuff package (Neal, 2003; Vanhatalo et *al.*, 2013) to sample the parameter posterior

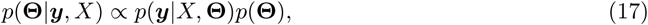

where the likelihood is defined in Eq. (7). After convergence checking from 4 independent Markov chains (details in Suppl. Sec. 2), we obtain S posterior samples 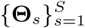, where 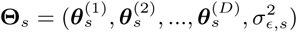. We use the posterior samples to approximate the predictive density for test data 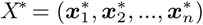

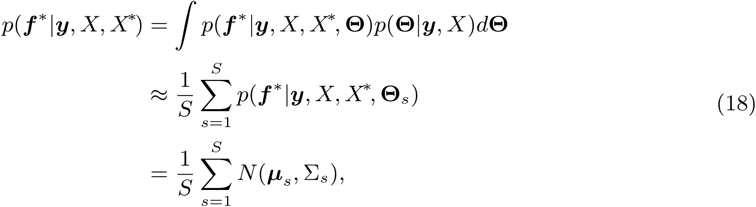

where

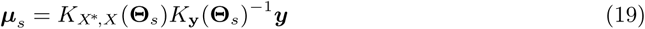

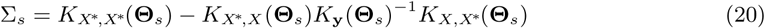

are the standard GP prediction equations adapted to additive GPs with 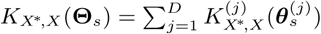 encoding the sum of cross-covariances between the inputs *X* and test data points *X\** (*k*_*x*\* *x*\*_ is defined similarly) and *K*_y_(Θ*_s_*) is defined in Eq. (8).

As an alternative approach to slice sampling for higher dimensional models, we also use a deterministic finite sum using the central composite design (CCD) to approximate the predictive densities for GPs as proposed in (Rue et *al.*, 2009; Vanhatalo et *al.*, 2010). CCD assumes a split-Gaussian posterior *q*(·) for (log-transformed) parameters *γ* = log(Θ) and defines a set of *R* points 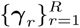 (fractional factorial design, the mode and so-called star points along whitened axes) to estimate the predictive density with a finite sum

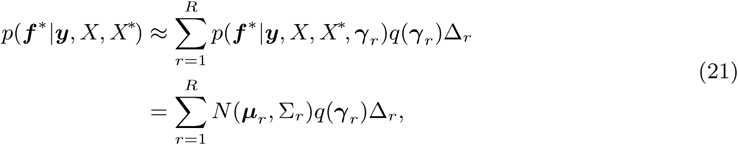

where *N*(*µ*_*r*_,Σ_*r*_) is computed as in Eqs. (19-20), *q*(*γ*_*r*_) is the split-Gaussian posterior and Δ_*r*_ are the area weights for the finite sum (see (Vanhatalo et *al.*, 2010) for details).

Predictions and visualisations for an individual kernel *k*^(^*^j^*^)^ (1 ≤ *j* ≤ *D*) are obtained by replacing ***µ***_*s*_ and Σ_*s*_ in Eqs. (18) and (21) with

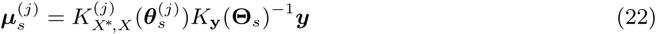

and

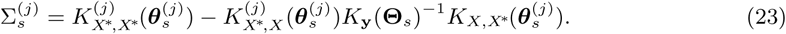

Similarly, predictions for a subset of kernels are obtained by replacing 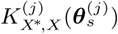 with the relevant sums.

### 2.7 Model comparison

We have described how to build and infer an additive GP model for a given target variable using a set of kernels and a set of covariates for each kernel. A model *M* can be specified by a 3-tuple 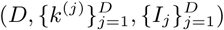, where *D* ≥ 1. However, all covariates may not be relevant for the prediction task and often the scientific question is to identify a subset of the covariates that are associated with the target variable. For model selection, we use two cross-validation variants and Bayesian bootstrap as described below.

#### 2.7.1 Leave-one-out cross-validation

We use leave-one-out cross-validation (LOOCV) to compare the models when a continuous covariate such as *age, diseaseAge* or *season* is added to a model. In this case, a single time point of an individual is left out as test data and the rest are kept as training data. We use MCMC to infer the parameters of a given model and calculate the following leave-one-out predictive density:

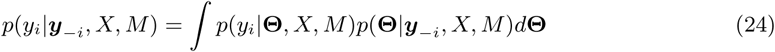

where ***y***_−*i*_ = ***y*** \*y*_*i*_ and Θ are the parameters of the GP model *M*. This can be calculated by setting ***f***\* ← *y_i_, X\** ← ***x***_*i*_, ***y*** ← ***y***_−*i*_ and *X* ← *X\x*_*i*_ in Eq. (18). The standard LOOCV would require us to run the inference *N* times, which is time consuming when *N* is large. In practice, we use importance sampling to sample *p*(Θ|***y***_-_*_i_, X, M*) where the posterior *p*(Θ|***y***, *X, M*) of the full data ***y*** is used as the proposal distribution. We thus approximate Eq. (24) as

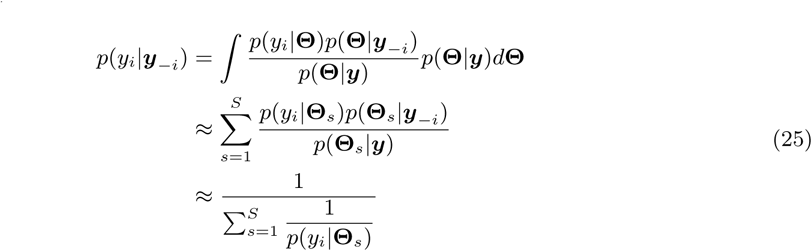

where we have omitted *X* and *M* in the notation for simplicity and Θ_*s*_ is a MCMC sample from the full posterior *p*(Θ|***y***). However, directly applying Eq. (25) usually results in high variance and is not recommended. We use a recently developed Pareto smoothed importance sampling to control the variance by smoothing the importance ratios *p*(Θ_*s*_|***y***_-*i*_)/*p*(Θ_*s*_|***y***) (for details, see (Vehtari *et al.*, 2017, 2016)).

The importance sampling phase is fast and it is shown to be accurate (Vehtari *et al.*, 2017). Therefore, we only need to run MCMC inference once for the full training data. Once the leave-one-out predictive probabilities in Eq. (24) are obtained for all the data points, the GP models are compared using Bayesian bootstrap described in Sec. 2.7.3.

#### 2.7.2 Stratified cross-validation

In stratified cross-validation (SCV), we leave out all time points of an individual as test data and use the rest as training data. SCV is used when a categorical/binary covariate, such as group or gender, is added to the model. Let ***y***_*i*_ denote all measured time points corresponding to an individual *i* (*X*_*i*_ is defined similarly) and ***y***_−*i*_ = ***y*** \ ***y***_*i*_. Similar to LOOCV, we want to compute the predictive density of the test data points ***y***_*i*_

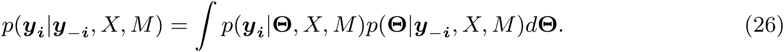

This can be calculated by setting ***f***\* ← *y_i_, X\** ← ***X***_*i*_, ***y*** ← ***y***_−*i*_ and *X* ← ***X***_−*i*_ in Eq. (21). Since importance sampling does not work well in this case, we apply the CCD inference *P* times (once for each individual). Also, we use CCD with SCV as it is much faster than MCMC.

#### 2.7.3 Model comparison using Bayesian bootstrap

After obtaining the leave-one-out predictive densities (Eq. (24) or (26)) for a collection of models, we use Bayesian bootstrap to compare the involved models. Let us start with a simple case where two models *M*_1_ and *M*_2_ are compared. In the LOOCV setting, we compare the models by computing the average difference of their log-predictive densities

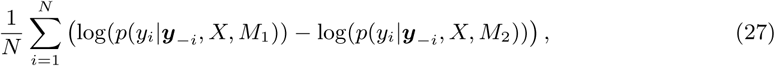

which measures the difference of the average prediction accuracy of the two models. If Eq. (27) is greater than 0, then model *M*_1_ is better than *M*_2_, otherwise model *M*_2_ is better than *M*_1_.

Comparison in Eq. (27) does not provide a probabilistic quantification of how much better one model is compared to the other. We thus approximate the relative probability of a model being better than another model using Bayesian bootstrap (Rubin, 1981), which assumes *y*_*i*_ only takes values from the observations ***y*** *=* (*y*_1_, *y*_2_, … *y*_*N*_)*^T^* and has zero probability at all other values. In Bayesian bootstrap, the probabilities of the observation values follow the N-dimensional Dirichlet distribution Dir(1, 1, …, 1). More specifically, we bootstrap the samples *N*_*B*_ times (*b* = 1,…, *N_B_*) and each time we get the same *N* observations ***y***, with each observation taking weight *w*_*bi*_ (*i* = 1, …, *N*) from the *N*-dimensional Dirichlet distribution. The *N*_*B*_ bootstrap samples are then summarised to obtain the probability of *M*_1_ being better than *M*_2_

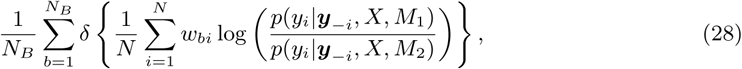

where *δ*{·} is the Heaviside step function and *w*_*bi*_ is the bootstrap weight for the *i*th data point in the *b*th bootstrap iteration (see Vehtari *et al.* (2017) for more details). We call the result of Eq. (28) LOOCV factor (LOOCVF).

The above strategy also works when comparing multiple models. Instead of calculating the heaviside step function in the *b*th bootstrap iteration, we simply choose the model with the highest rank by sorting the models using

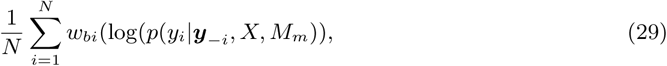

where *m* indices the model. In the end, we count the occurrences *N*_*m*_ of each model being the best across all *N*_*B*_ bootstrap samples and we compute the posterior probability of model *M*_*m*_ as *N*_*m*_/*N*_*B*_, which we term as the posterior rank probability.

For SCV, we replace *y*_*i*_ with ***y***_*i*_ and ***y***_*-i*_ with ***y***_*-i*_ in Eqs. (27-28) and follow the same procedure as above to compare the models. Eq. (28) is then termed as the SCV factor (SCVF). In practice, we set the threshold of the LOOCVF to be 0.8 and SCVF to be 0.95, i.e., the LOOCVF (resp. SCVF) of the extended model versus the original model needs to be larger than 0.8 (resp. 0.95) for a continuous covariate (resp. binary covariate) to be added.

Although Eq. (29) can be used to compare any subset of models, complex models will dominate the posterior rank probability when compared together with simpler models. Hence, LonGP only uses it to compare candidate models of similar complexity (see next Section and Suppl. Sec. 3).

### 2.8 Step-wise additive GP regression algorithm

The space of all models is large and thus an exhaustive search for the best model over the whole model space would be too slow in practice. Two commonly used model (or feature) selection methods include forward and backward search techniques. Starting with the most complex model, as in the backward search approach, is not practical in our case, so we propose to use a greedy forward search approach similar to step-wise linear regression model building. That is, we start from the base model that only includes the id covariate. Then we add continuous covariates to the model sequentially until the model cannot be further improved. During each iteration, we first identify the covariate that improves the model the most (Eq. (29)) and test if the LOOCVF of a new proposed model versus the current model exceeds the threshold of 0.8 (Eq. (28)). While including a continuous covariate, we also include relevant interaction terms (allowed interaction terms defined by the user). After adding continuous covariates, we add discrete (categorical or binary) covariates sequentially to the model until it cannot be further improved. As with continuous covariates, during each iteration, we first identify the discrete covariate that improves the model the most and test if the SCVF of a new proposed model versus the current model exceeds the threshold of 0.95. While including a discrete covariate, we also include relevant interaction terms (allowed interactions specified by user). Details of our forward search algorithm are given in Suppl. Sec. 3 together with a pseudo-algorithm description. We note that although step-wise model selection strategies are commonly used with essentially all modelling frameworks, they have the danger of overfitting a given data. To avoid overfitting, we implement our search algorithm such that an additional component is added to the current model only if the more complex model improves the model fit significantly, as measured by the LOOCVF and SCVF.

Once all the covariates have been added, the kernel parameters of the final model are sampled using MCMC and kernel-specific predictions on the training data *X* are computed using Eq. (18). Additionally, a user can choose to exclude kernels that have a small effect size as measured by the fraction of total variance explained. we require component specific variances to be at least 1%. The software is implemented using features from the GPStuff package (Vanhatalo *et al.*, 2013) and implementation is discussed in Suppl. Sec. 4.

## 3 Results

We tested LonGP on simulated datasets and two real datasets including longitudinal metagenomics (Vatanen *et al.*, 2016) and proteomics datasets (Liu *et al.*, 2018).

### 3.1 Simulated datasets

We first carried out a large simulation study to test and demonstrate LonGP’s ability to correctly infer associations between covariates and target variables from longitudinal data. Here we are primarily interested in answering two questions: is LonGP able to select the correct model as well as the correct covariates that were used to generate the data, and can we detect disease associated signals. We simulated -omics datasets from five different generating additive GP models (AGPM):

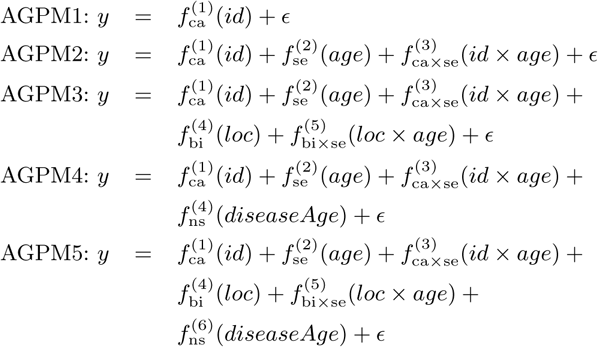

To set up our simulation scenario, we first use 20 cases and 20 controls (i.e., *P* = 40) specified by the *group* covariate, each with *n*_*i*_ = 13 data points ranging from 0 month to 36 months with an increment of three months, thus specifying the *age* covariate. Other covariates are randomly simulated using the following rules. The disease occurrence time is sampled uniformly from 0 to 36 months for each case subject and *diseaseAge* is computed accordingly. We make the effect of *diseaseAge* non-stationary by transforming it with the sigmoid function from Eq. (15), such that majority of changes occur in the range of -12 to +12 months. The *location* and *gender* are i.i.d. sampled from a Bernoulli distribution with *p* = 0.5 for each individual, where *gender* acts as an irrelevant covariate. The continuous covariates are subjected to standardisation after being generated, such that the mean of each covariate is 0 and standard deviation is 1. We then use the kernels described in Sec. 2.4, where the length-scales for continuous (standardised) covariates are set to 1 for the shared components and 0.8 for the interaction components. We set the variances of each shared component to 4 and noise to 3, i.e., 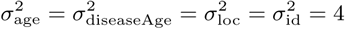 and 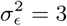. With these specifications, we generate 100 datasets for each AGPM. A randomly generated longitudinal data set from AGPM5 is visualised in Fig. 1 (Note the order of latent functions is changed for better visualisation.).

In the inference, all covariates including gender are used, which means that there are 2^4^ = 16 candidate models to be selected. Interaction terms are allowed for all covariates except for *diseaseAge*. Table 1 shows the distribution of selected models for each generating additive GP model, with the numbers in bold font indicating correctly identified models. Table 1 shows that LonGP can achieve between 88 and 98% accuracy in inferring the correct model with these parameter settings. Results in Table 1 also shows that it becomes more challenging to identify the correct model as the generating model becomes more complex, which is expected. LonGP can accurately detect the disease related signal as well since the diseaseAge covariate is included in the final model in 97% of the simulation runs for both AGPM4 and AGPM5 models (see Table 1). Moreover, LonGP is notably specific in detecting the *diseaseAge* covariate as the percentage of false positives is only 0%, 1%, and 0% for AGPM1, AGPM2, and AGPM3, respectively (see Table 1).

**Table 1.**
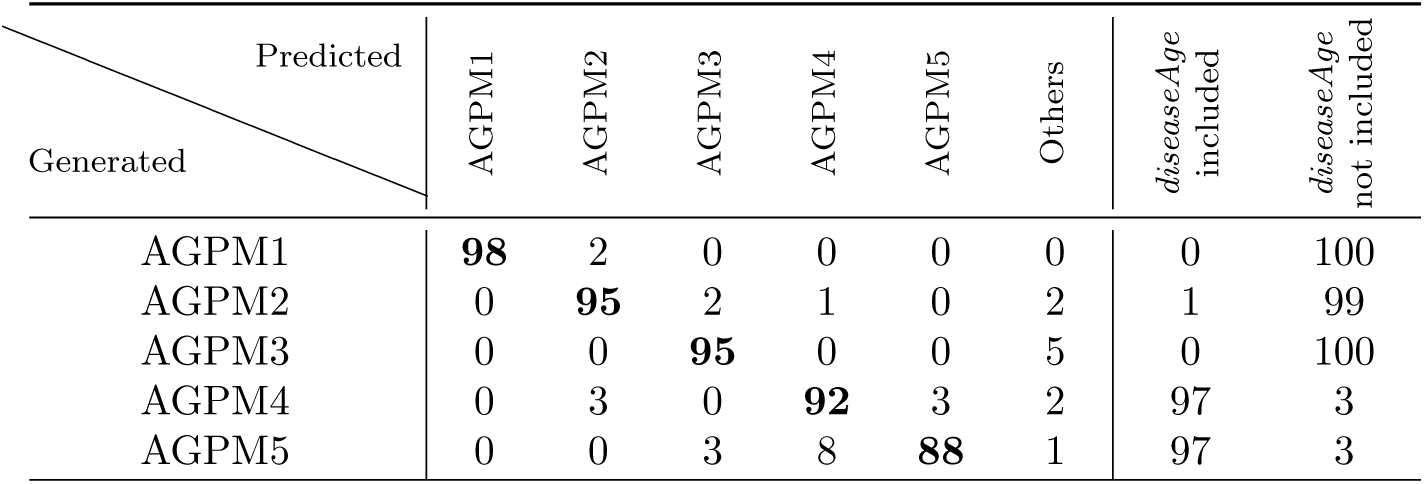
Model inference results for simulated data with 20 cases and 20 controls, noise variance 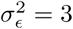 and samples taken every 3 months. Rows show the number of times each model is inferred as the best model out of 100 Monte Carlo simulations for each generating model. ‘Others’ corresponds to all the other 11 possible APGM models. The last two columns show the number of times the diseaseAge covariate has or has not been included in the final model

| Generated \ Predicted |  |  |  |  |  |  | *diseaseAge*<br>included | *diseaseAge*<br>not included |
| --- | --- | --- | --- | --- | --- | --- | --- | --- |
|  | AGPM1 | AGPM2 | AGPM3 | AGPM4 | AGPM5 | Others |  |  |
| AGPM1 | **98** | 2 | 0 | 0 | 0 | 0 | 0 | 100 |
| AGPM2 | 0 | **95** | 2 | 1 | 0 | 2 | 1 | 99 |
| AGPM3 | 0 | 0 | **95** | 0 | 0 | 5 | 0 | 100 |
| AGPM4 | 0 | 3 | 0 | **92** | 3 | 2 | 97 | 3 |
| AGPM5 | 0 | 0 | 3 | 8 | **88** | 1 | 97 | 3 |

To better characterise LonGP’s performance in different scenarios, we tested how the amount of additive noise affects the results. We varied the noise variance as 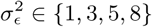 and kept all other settings unchanged, effectively changing the signal to noise ratio, or the effect size relative to the noise level. Fig. 2a) shows that the model selection accuracy increases consistently as the noise variance decreases. We next tested how the number of study subjects (i.e., the sample size *P*) affects the inference results. We set the number of case-control pairs to {(10, 10),(20,20), (30,30), (40,40)} and keep all other settings unchanged. As expected, Fig. 2b) shows how LonGP’s model selection accuracy increases as the sample size increases. Similarly, LonGP maintains its high sensitivity and specificity in detecting diseaseAge covariate across the additive noise variances and samples sizes considered here (see Suppl. Tables 5 and 6).

**Figure 2:**
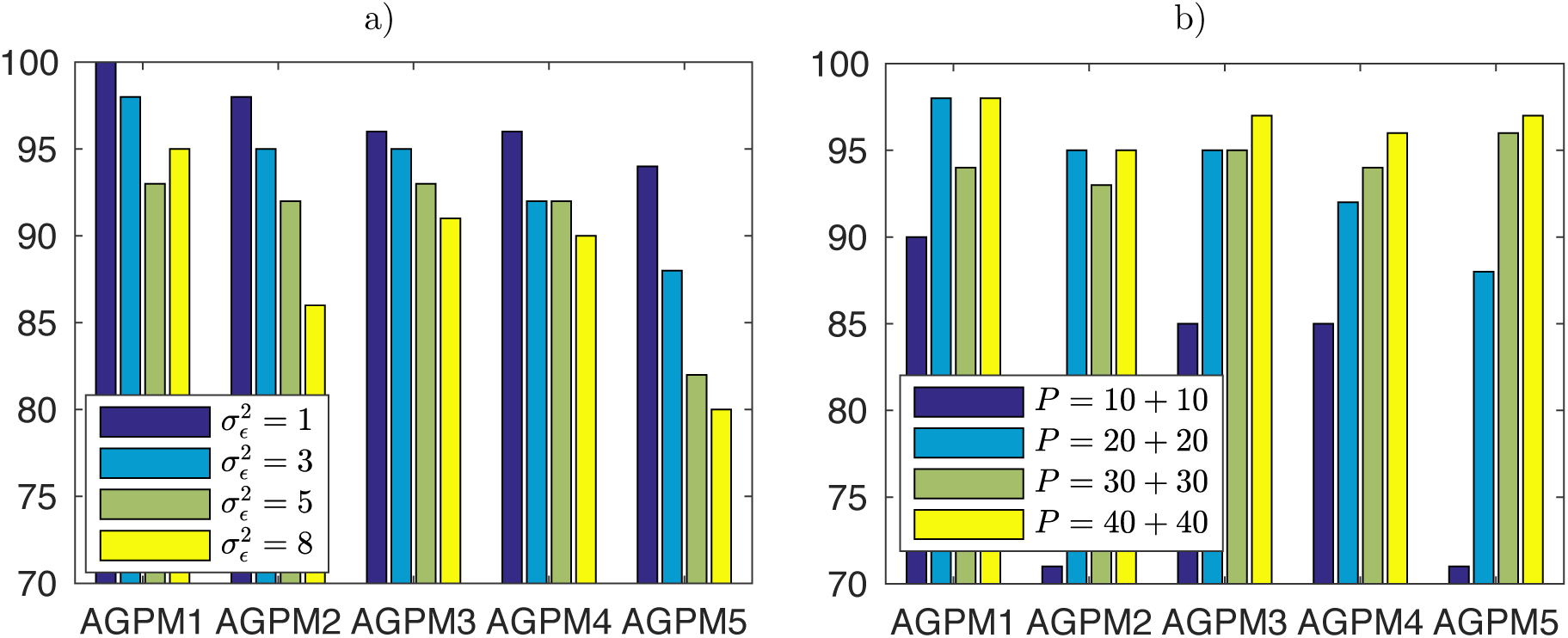
a) Model selection accuracy as a function of noise variance. b) Model selection accuracy as a function of sample size. *y*-axis shows the number of times the correct model is inferred as the best model out of 100 Monte Carlo simulations.

Finally, we also quantify how the sampling interval (i.e., the number of time points per individual) affects the inference results. We varied the sampling intervals as {2,3, 4,6} (months) corresponding to *n*_*i*_ ∈ {19, 13,10, 7} time points for each individual and kept all other simulation settings unchanged. Suppl. Table 3 shows that again the model selection accuracy changes consistently with the number of measurement time points. Suppl. Table 7 shows that changing the sampling interval has a small but systematic effect on the sensitivity and specificity of detecting the *diseaseAge* covariate.

Overall, our results suggest that we can accurately infer the correct model structure and also detect a relatively weak disease related signal with as few as 10 case-control pairs and notable noise variance. Moreover, the model selection accuracy increases as the number of individuals (biological replicates), the number of time points and signal to noise ratio increases.

### 3.2 Longitudinal metagenomics dataset

We used LonGP to analyse a longitudinal metagenomics dataset (Vatanen *et al.*, 2016). In this dataset, 222 children from Estonia, Finland and Russia were followed from birth until the age of three with collection of monthly stool samples which were subsequently analysed by metagenomic sequencing. The aim of this study was to characterise the developing gut microbiome in infants from countries with different socioeconomic status and to determine the key factors affecting the early gut microbiome development. Here we model the microbial pathway profiles quantifying the functional potential of the metagenomic communities. There are in total *N* = 785 metagenomic samples. We require a pathway to be detected in at least 500 samples to be included in our LonGP analysis, which results in 394 valid microbial pathways. Let *c*_*ij*_ denote the number of reads mapping to genes in the *j*th (*j* = 1, …, 394) pathway in sample *i* (*i* = 1, …, 785) and *C*_*i*_ is the total number of sequencing reads for sample *i*. The target variable is defined by *c*_*ij*_/*C*_*i*_ · median(*C*_1_, *C*_2_, …, *C*_N_).

We selected the following 7 covariates for our additive GP regression based on their known interaction with the gut microbiome: age, bfo, caesarean, est, fin, rus and *id. bfo* indicates whether an infant was breastfed at the time of sample *collection*; caesarean indicates if an infant was born by Caesarean section; *est, fin* and *rus* are binary covariates indicating the home country of the study subjects (Estonia, Finland and Russia, respectively). We use SE kernel for *age* and *bfo*, categorical kernel for *id*, and binary kernel for *caesarean, est, fin*, and *rus*. Interactions are allowed for all covariates except for *bfo*.

We applied LonGP to analyse each microbial pathway as a target variable separately and inferred the covariates for each target variable as described above. The selected models and explained variances of the components for all 394 pathways are available in Suppl. File 1. A key discovery in Vatanen *et al.* (Vatanen *et al.*, 2016) was that “Lipid A biosynthesis” pathway was significantly enriched in the gut microbiomes of Finnish children compared to Russian children. Our analysis confirmed the linear model based analysis in (Vatanen *et al.*, 2016) by selecting the following model for “Lipid A biosynthesis” pathway: 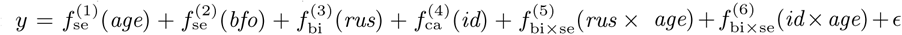, which shows the difference between the Russian and Finnish study groups. Explained variance of *bfo* was 0.2% and *bfo* was thus excluded from the final model. Fig. 3 shows the normalized “Lipid A biosynthesis” data together with the additive GP predictions using kernels 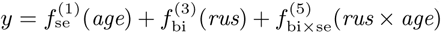. The obtained model fit is similar to that reported in (Vatanen *et al.*, 2016) with an exception that the apparent non-linearity is captured by the additive GP model but otherwise the new model conveys the same information. Our analysis also identified many novel pathways with differences between Finnish, Estonian and Russian microbiomes, reported in Suppl. File 1.

**Figure 3:**
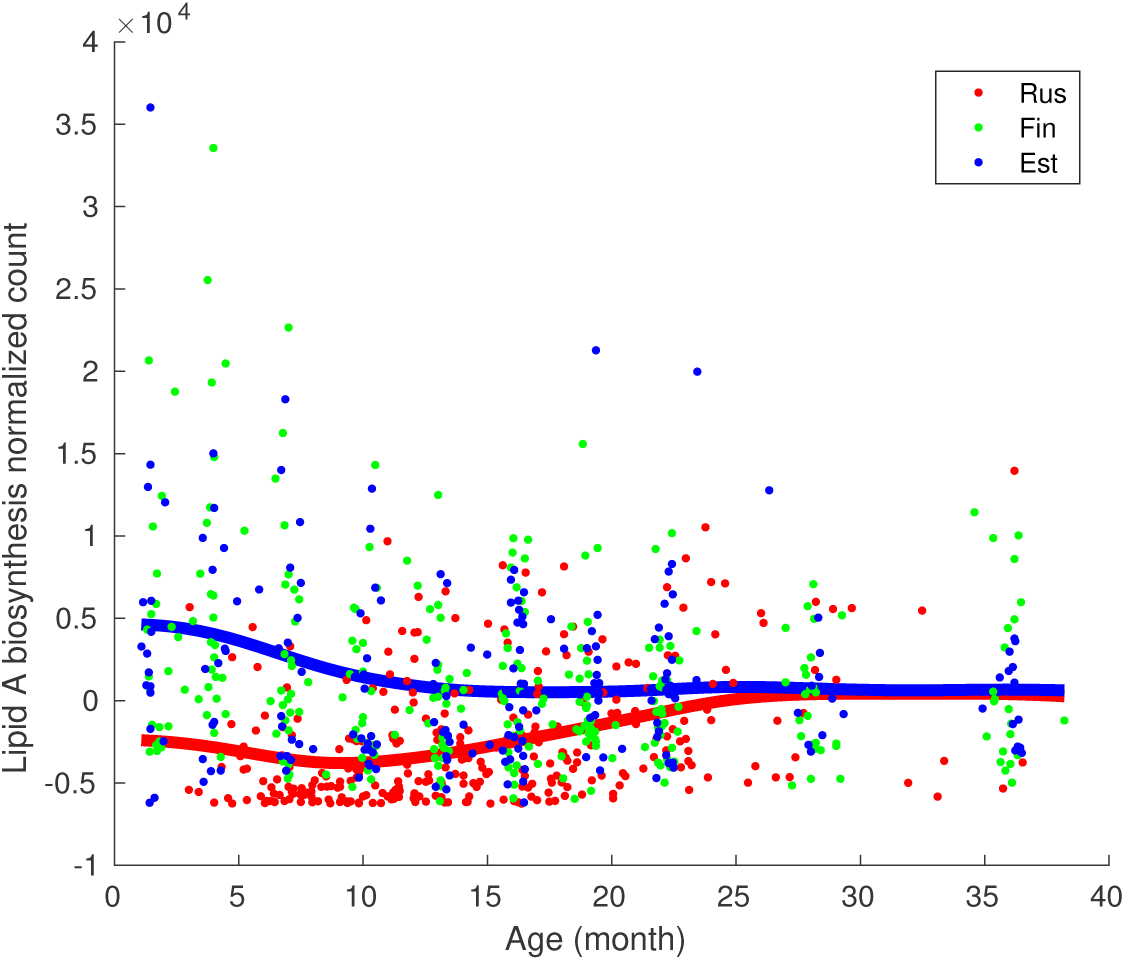
LonGP regression results for “Lipid A biosynthesis” pathway. Normalized read counts of Russian, Finnish and Estonian infant samples are colored by red, green and blue dots, respectively. The blue line shows the nonlinear age trend for Finnish and Estonian infants. The red line shows the age trend of Russian infants. The red and blue lines are generated as the sum of components 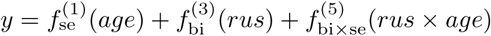.

### 3.3 Longitudinal proteomics dataset

We next analysed a longitudinal proteomics dataset from a type 1 diabetes (T1D) study (Liu *et al.*, 2018). Liu et al. measured the intensities of more than 2000 proteins from plasma samples of 11 T1D patients and 10 healthy controls which were collected at 9 time points, resulting in a total of 189 samples. Detection of T1D associated auto-antibodies in the blood is currently held as the best early marker that predict the future development of T1D, and most of the individuals turning positive for multiple T1D auto-antibodies will later on develop the clinical disease. The disease event of interest is called seroconversion, which is the first time point when T1D-specific antibodies are detected in blood. Identifying early markers for T1D that would be detected even before the auto-antibodies is a grand challenge. It would allow early disease prediction and possibly even intervention.

Liu et al. used a linear mixed model with quadratic terms to detect proteins that behave differently between cases and controls. However, they did not model changes near the seroconversion in their model and only regressed on age. We use LonGP to re-analyse this longitudinal proteomics dataset (Liu *et al.*, 2018) and try to find additional proteins with differing plasma expression profiles between cases and controls in general as well as focusing on samples collected close to seroconversion. The modelling is done with the following covariates: *age, sero* (measurement time minus seroconversion time), *group* (case or control), *gender*, and *id*. 1538 proteins with less than 50% missing values are kept for further analysis. We follow the same preprocessing steps as described in (Liu *et al.*, 2018) to get the normalised protein intensities. We use SE kernel for *age*, input warped non-stationary SE kernel for *sero*, binary kernel for *group* as well as for *gender*, and categorical kernel for *id*. Interactions are allowed for all covariates except for *sero*. The selected models and explained variances of each component for all 1538 proteins are reported in Suppl. File 2.

We detected 38 proteins that are associated with the group covariate. Protein with Uniprot ID Q7LGC8 shows a group difference (the protein level of cases are higher than controls) and the selected model is 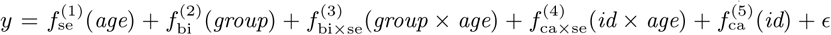. Fig. 4 shows the contribution of each component and the cumulative effects. Fig. 5 shows the cumulative effect 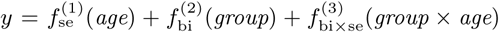 against the real protein intensity to better visualise the predicted group difference.

**Figure 4:**
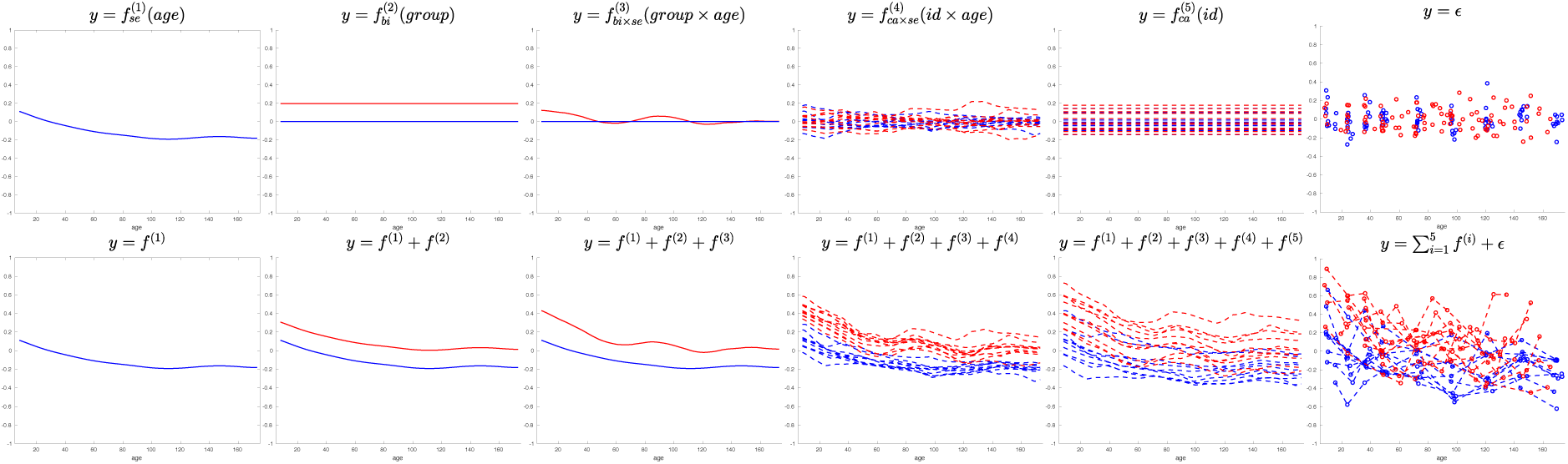
Predicted components and cumulative effect for protein Q7LGC8. Top panel shows contributions of individual components and lower panel shows cumulative effects. Red lines are cases and blue lines are controls. Bottom right panel shows the (centered) data.

**Figure 5:**
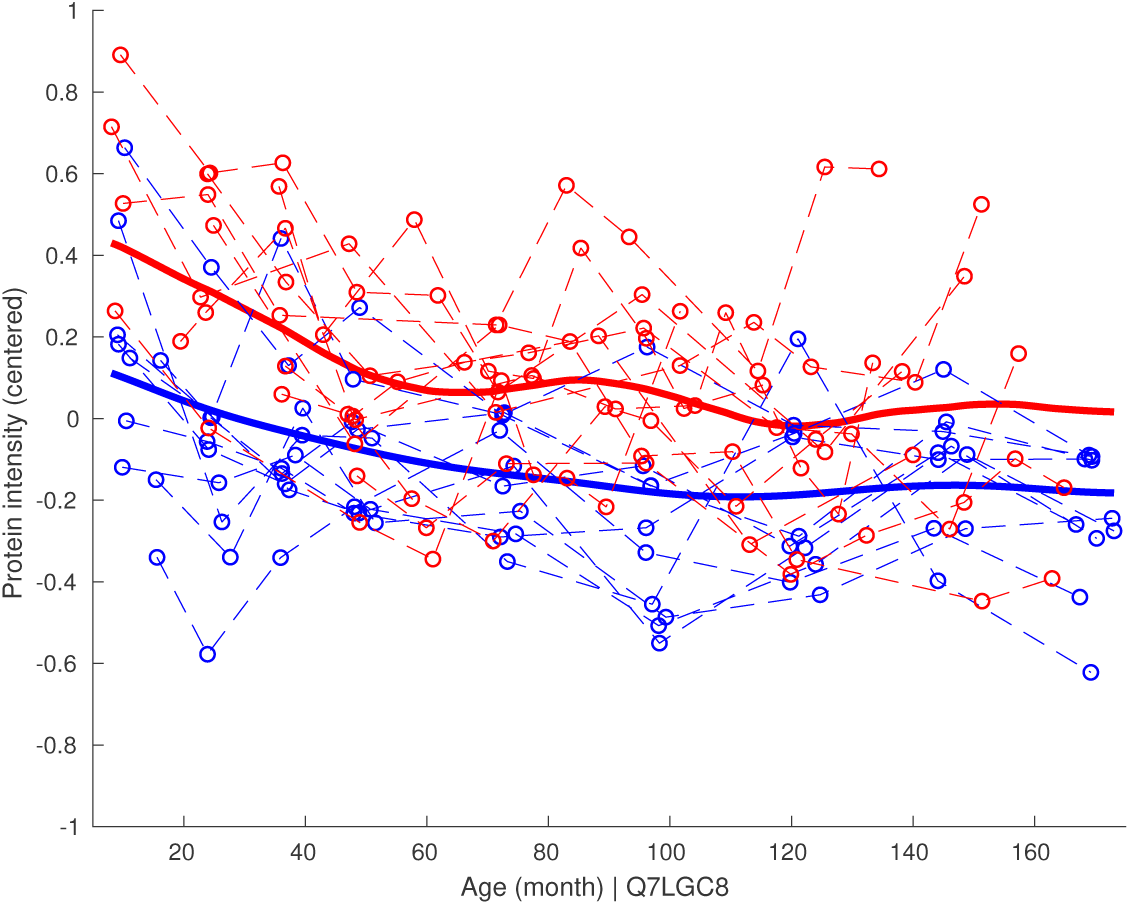
Cumulative effect 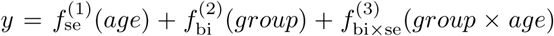 against real (centered) intensity of protein Q7LGC8. Red lines are cases and blue lines are controls.

We detected 30 proteins that are associated with the sero covariate. We visualise two of those proteins (Uniprot IDs: P07602, Q14982) that show a signal near seroconversion time point. For both proteins LonGP detects model 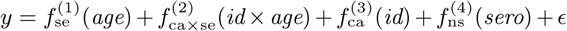. Fig. 6 shows the contribution of the *sero* component together with the real (centered) protein intensities as a function of seroconversion age for protein P07602. The *sero* component increases and then stablises at a higher baseline after seroconversion in the cases. This is shown by the lower baseline of cases before seroconversion and higher baselines after seroconversion. Suppl. Fig. 5 shows the predicted mean of each component as well as the cumulative effects for protein P07602. Suppl. Fig. 6 shows a different type of *sero* effect for protein Q14982 where a temporary increase of the protein intensity near the seroconversion event is observed in many T1D patients, in contrast to the slowly decreasing age trend. Suppl. Fig. 7 shows the predicted individual components and the cumulative effects for protein Q14982.

**Figure 6:**
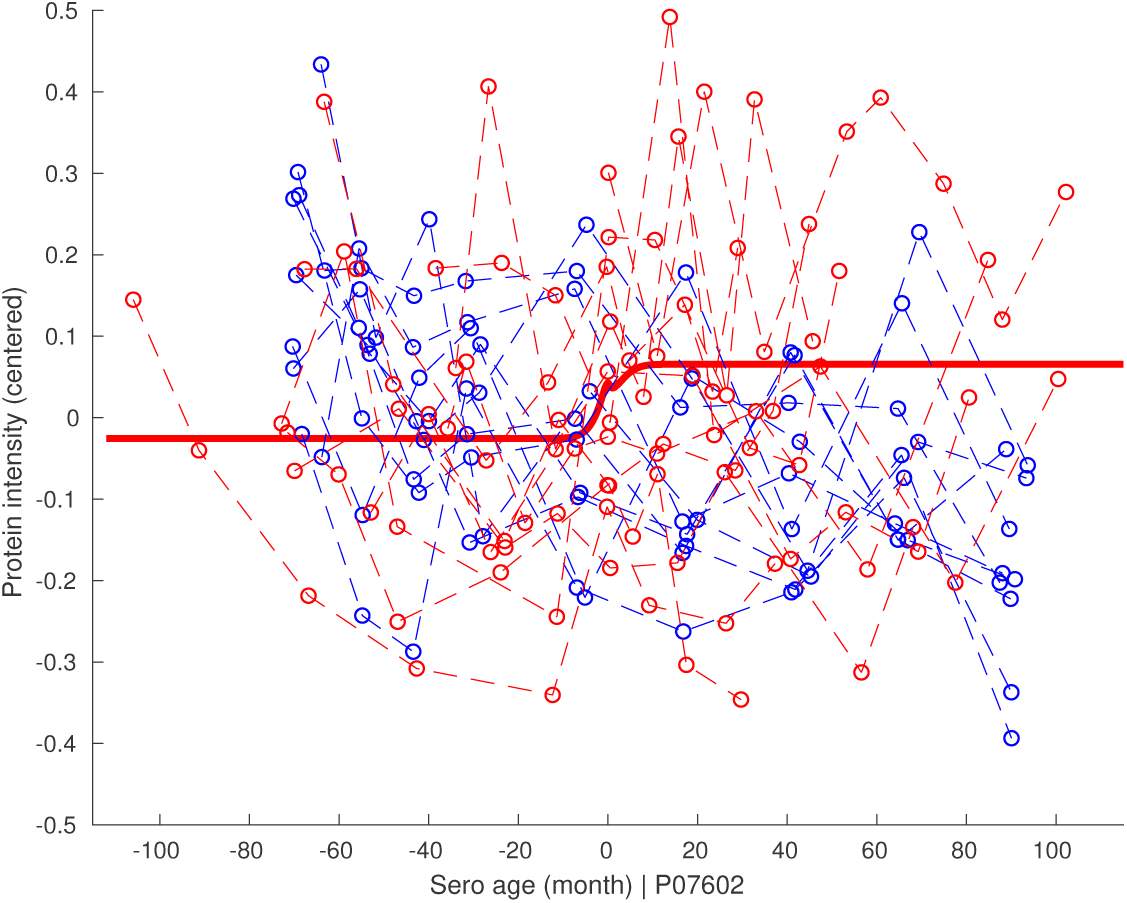
Predicted mean of the sero component for protein Q14982. The dashed red lines show the measurements of cases and the dashed blue lines are measurements of controls. *x*-axis is seroconversion age and *y*-axis is centered protein intensity. Mean seroconversion age of all cases (79.42 month) is used as the seroconversion age for controls. The solid red line corresponds to the mean of the seroconversion component 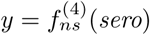.

## 4 Discussion and Conclusions

General linear mixed effect model is a simple yet powerful modelling framework that has been widely accepted in biomedical literature. Still, applications of linear models can be challenging, especially when the underlying data generating mechanisms contain unknown nonlinear effects and correlation structures or non-stationary signals.

Here we have described LonGP, a non-parametric additive Gaussian process model for longitudinal data analysis, which we demonstrate to solve many of the commonly faced modelling challenges. As LonGP builds on GP regression, it can automatically handle irregular sampling time points and time-varying covariates. Missing values are also easily accounted for via binary mask kernels without any extra effort. More generally, LonGP provides a flexible framework to choose appropriate covariance structures for the correlated outcomes via the GP kernel functions, and the chosen kernels are properly adjusted to given data by carrying out Bayesian inference for the kernel parameters. Gaussian processes are known to be capable of approximating any continuous function. Thus, LonGP is applicable to any longitudinal data set. Furthermore, incorporating non-stationary kernels into the kernel mixture easily adapts LonGP for non-stationary signals. Finally, LonGP is equipped with an advanced Bayesian predictive inference method that utilises several recent, state-of-the-art techniques which make model inference accurate and improves running time especially for larger data sizes and more complex models.

Compared with traditional linear regression methods, LonGP is helpful in finding relatively weak signals that have an arbitrary shape. For protein P07602 in the longitudinal proteomics dataset (Liu *et al.*, 2018), the dominant factor is age (explained variance 25%) and the disease related effect sero (explained variance 5.5%) is a minor factor, as shown in Suppl. Fig. 5. Revealing such disease related effects is essential in understanding mechanisms of disease progression and uncovering biomarkers for diagnostic purposes. The seroconversion associated proteins revealed by our study provide a list of candidate proteins for further analysis with a more extensive sample size using, for example, targeted proteomics approaches. Similarly, in the longitudinal metagenomics dataset (Vatanen *et al.*, 2016), we also observe non-linear effects for many of the covariates, some of which warrant further experimental studies.

Overall, supported by our results, we believe LonGP can be a valuable tool in longitudinal data analysis.

## Acknowledgements

We acknowledge the computational resources provided by the Aalto Science-IT and CSC-IT Center for Science, Finland.

## LonGP: an additive Gaussian process regression model for longitudinal study designs (supplementary)

Lu Cheng, Siddharth Ramchandran, Tommi Vatanen, Niina Lietzen, Riitta Lahesmaa, Aki Vehtari and Harri Lähdesmäki

January 29, 2018

### 1 Supplementary figures

**Figure 1:**
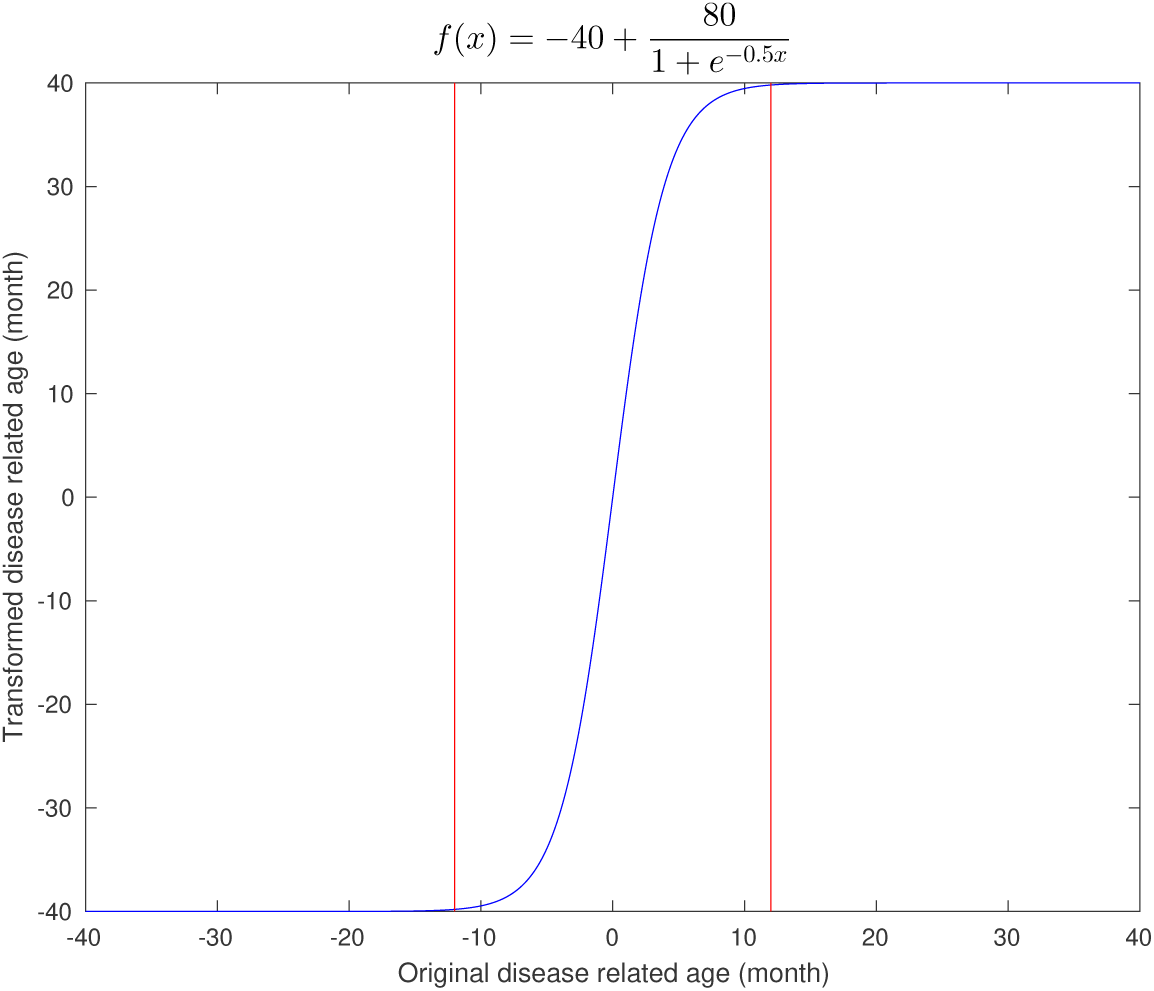
Non-stationary transformation. The x-axis is the original disease related age and the y-axis is the transformed disease related age. Sigmoid function 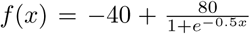 is used for the transformation. The red bars indicate the positions of ±12 month.

**Figure 2:**
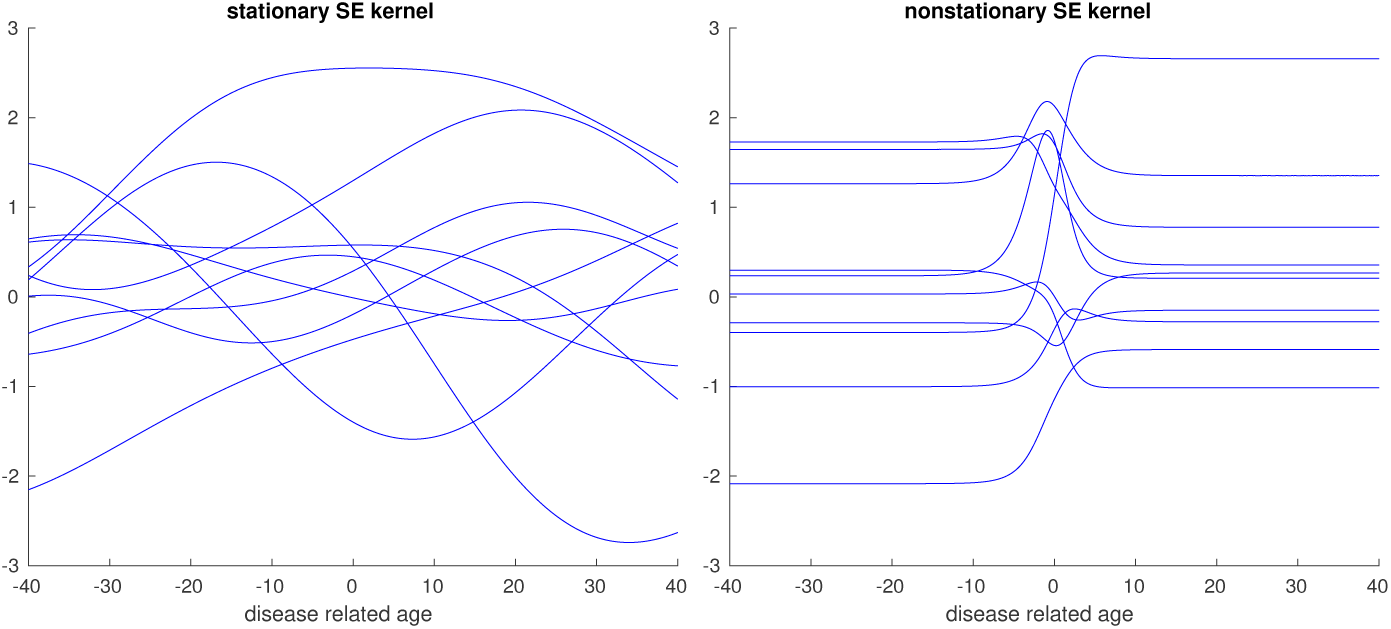
Functions drawn from stationary and non-stationary SE kernel. The left panel shows functions drawn form a stationary SE kernel with length-scale *l*_se_ = 1 and magnitude 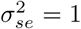. The right panel shows functions drawn form a non-stationary SE kernel by first applying the transformation shown in Figure 1 and then generated using the same SE kernel with scale *l*_se_ = 1 and magnitude 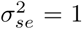. Random functions are drawn using the standardised inputs and then transformed back to original range.

**Figure 3:**
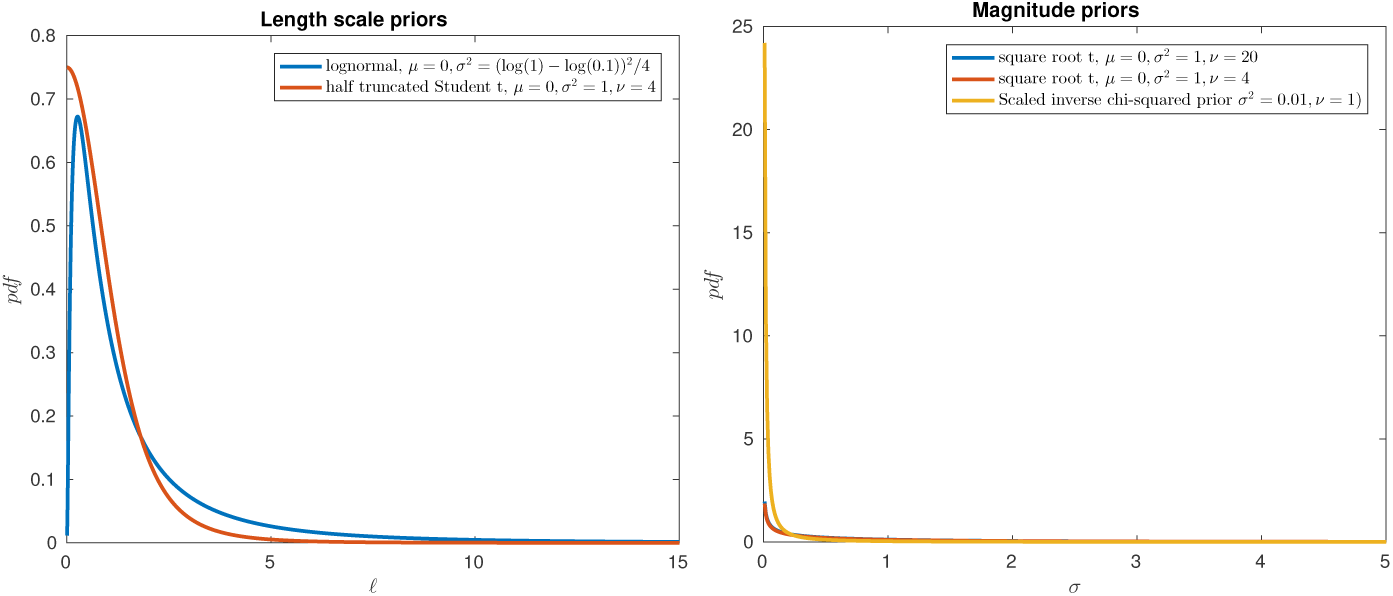
Priors for kernel parameter. The left panel shows priors for length-scales and the right panel shows priors for magnitude and noise variance. Note that the target variable and continuous covariates are all standardised to mean 0 and standard deviation 1.

**Figure 4:**
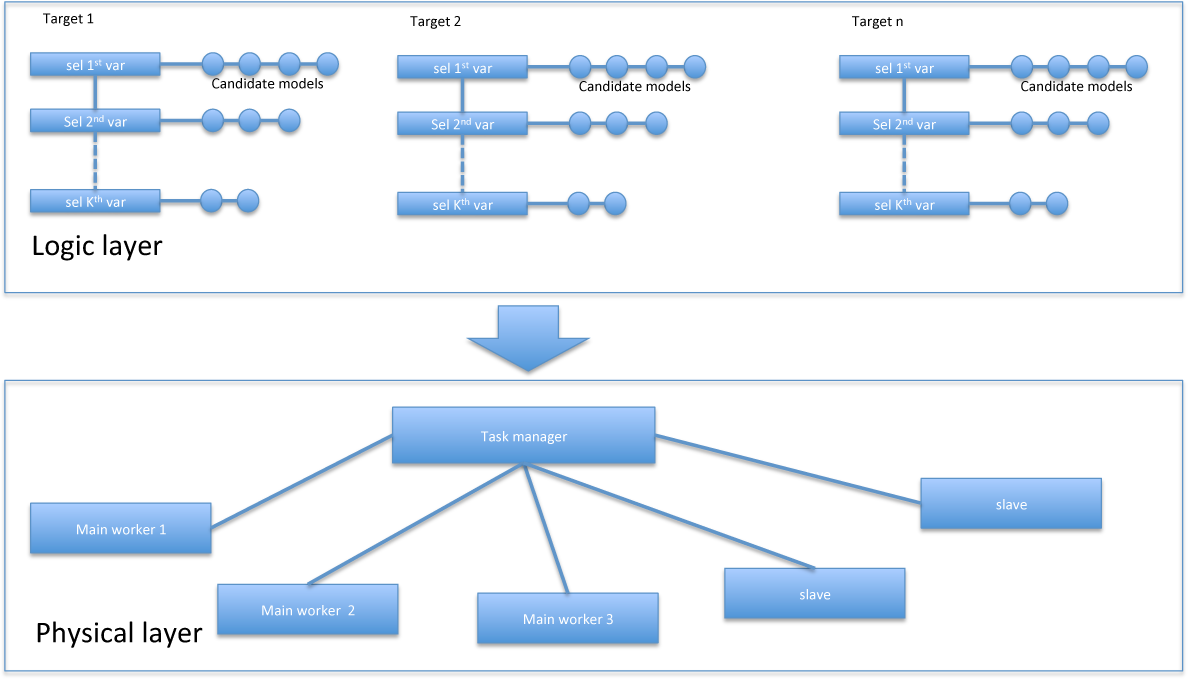
Software architecture. The task manager monitors the whole process and schedules the tasks. The main worker ensures the tasks for a given target is executed in the right order. The slaves run parallel jobs assigned by the task managers.

**Figure 5:**
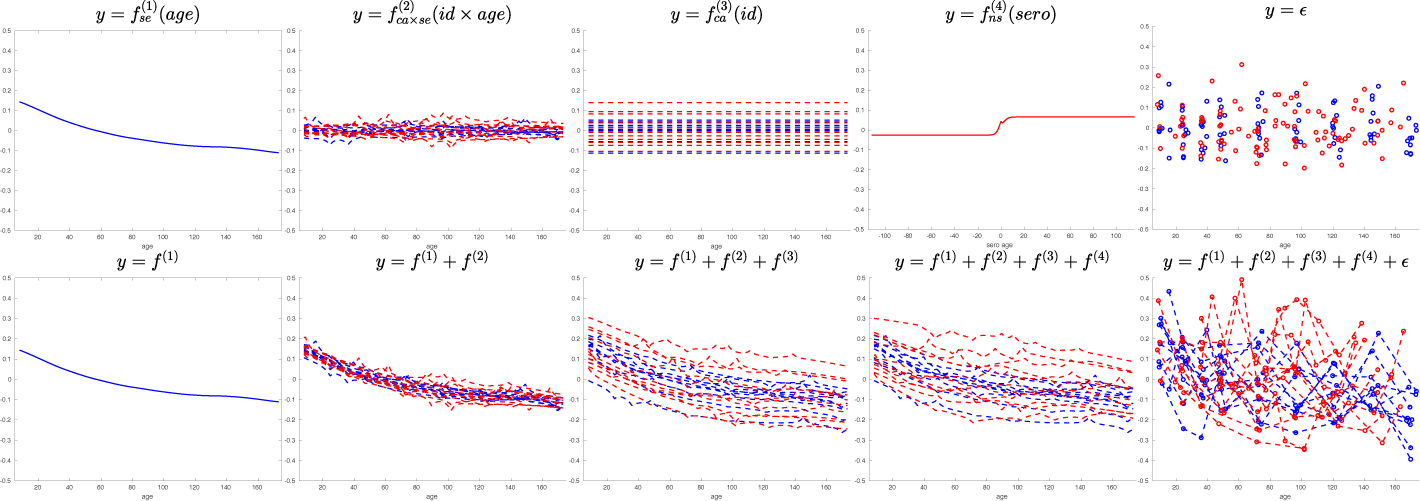
Predicted components and cumulative effects for protein P07602. Top panel shows contributions of individual components and lower panel shows cumulative effects. Red lines correspond to cases and blue lines correspond to controls. Bottom right panel shows the (centered) data. Note the *x*-axis of *f*^(4)^ is seroconversion age.

**Figure 6:**
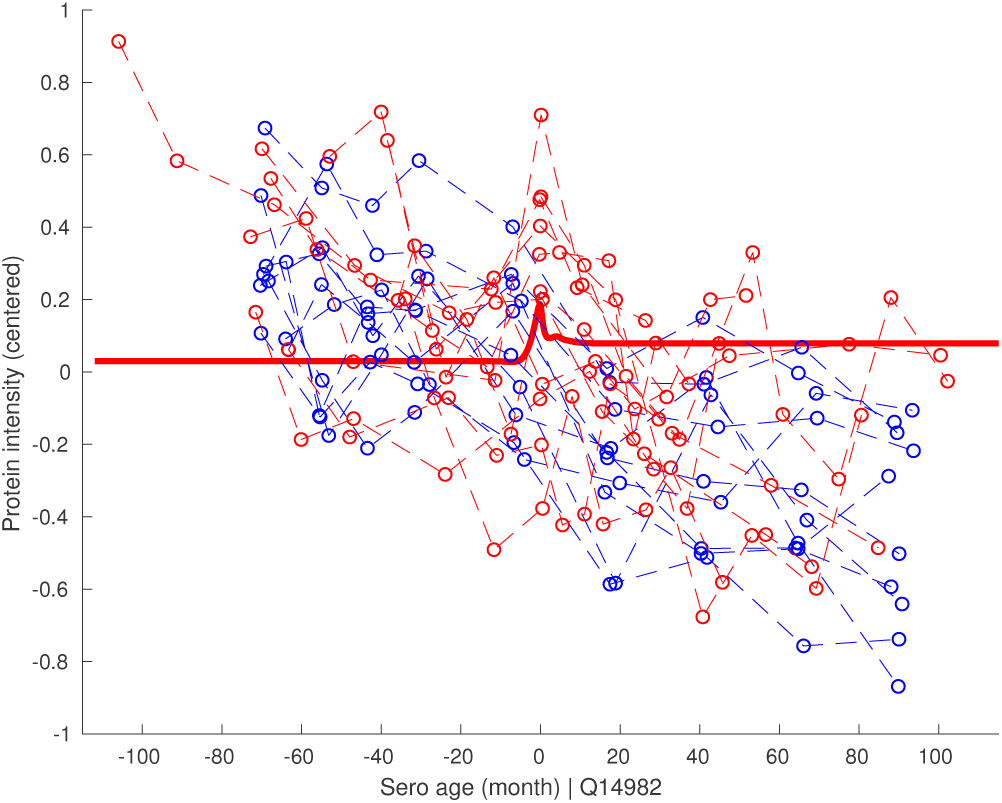
Predicted mean of the sero component for protein P07602. The dashed red lines show the measurements of cases and the dashed blue lines are controls. *x*-axis is seroconversion age and *y*-axis is centered protein intensity. Mean seroconversion age of all cases (79.42 month) is used as the seroconversion age for controls. The solid red line corresponds to the mean of the seroconversion component 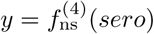.

**Figure 7:**
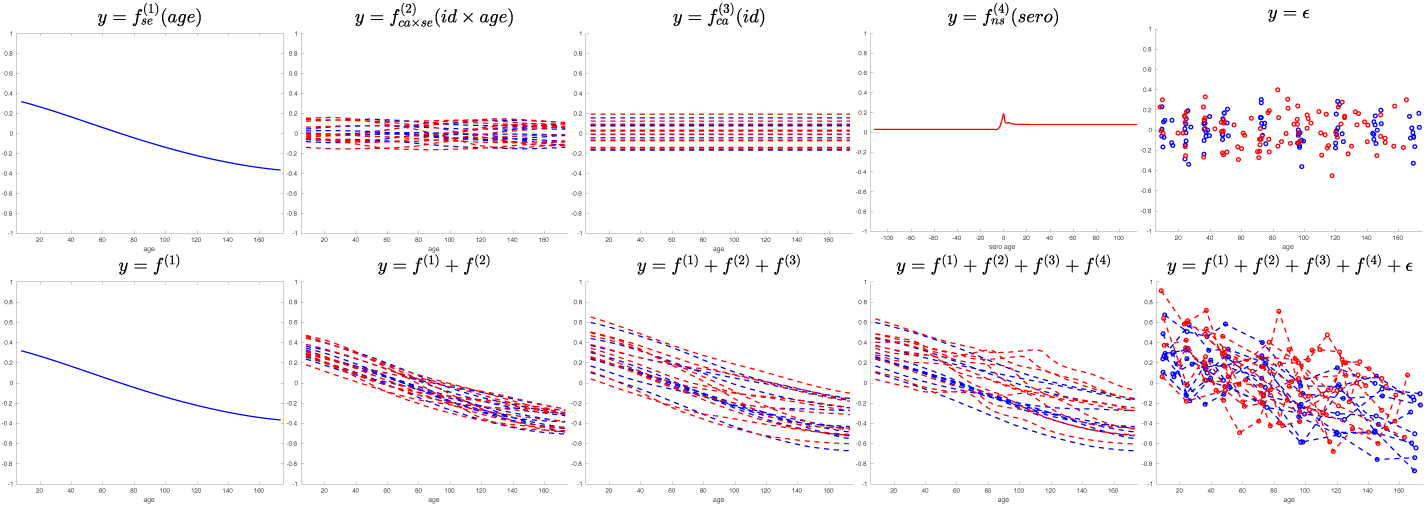
Predicted components and cumulative effects for protein Q14982. Top panel shows contributions of individual components and lower panel shows cumulative effects. Red lines correspond to cases and blue lines correspond to controls. Bottom right panel shows the (centered) data. Note the *x*-axis of *f*^(4)^ is seroconversion age.

### 2 MCMC details

We start 4 independent Markov chains from different, randomly initialised initial parameter values. Then, we combine the 4 chains and check the convergence by throwing away 500 burn-in samples and thinning the remaining 2000 samples by 5. If converged, then quit; otherwise we thin the combined chain further by 2. If not converged, we repeat the process and check the convergence from the resulting combined markov chains, for at most 4 times. The potential reduction scaling factor (PRSF) [1] *R* is used to check the convergence by the following rules: if *R* <= 1.1, converged; if 1.1 < *R* <= 1.2, does not converge well; if *R* > 1.2, does not converge.

### 3 LonGP algorithm

This section describes in detail how the covariate selection process works. Let us denote a given set of continuous covariates by **C** = (*V*_1_,*V*_2_, …., *V*_*c*_) and the binary covariates by **B** = (*V*_*c*+1_,*V*_*c*+2_, …., *V*_*c+b*_), where *c* and *b* are the number of continuous and binary/categorical variables. The categorical covariate id must be included in set **B**. In LonGP, the user needs to provide the kernel types (Sec. 2.4) for all the given covariates, as well as indicate whether interactions for each covariate is allowed. The data are automatically standardised and the parameter priors for kernels are predefined (see Sec. 2.5). For any given subset of covariates (must include *id*), the additive GP model is constructed by the following rules:

1. Construct a kernel for each covariate according to the given kernel type and add it to the model.
2. For each continuous covariate that allows interaction, construct product kernels with all categorical/binary covariates that also allow interactions (and that are also covariates of a given model) and add them to the model.
3. For each pair of categorical/binary covariates (excluding *id*) that allows interactions, construct a product kernel and add it to the model.
4. Add the noise to finalise the model.

For any covariate subset **V**, we can construct a GP model GPM(**V**) according to these four steps. The covariates are then selected by the following algorithm:

**Algorithm 1:**
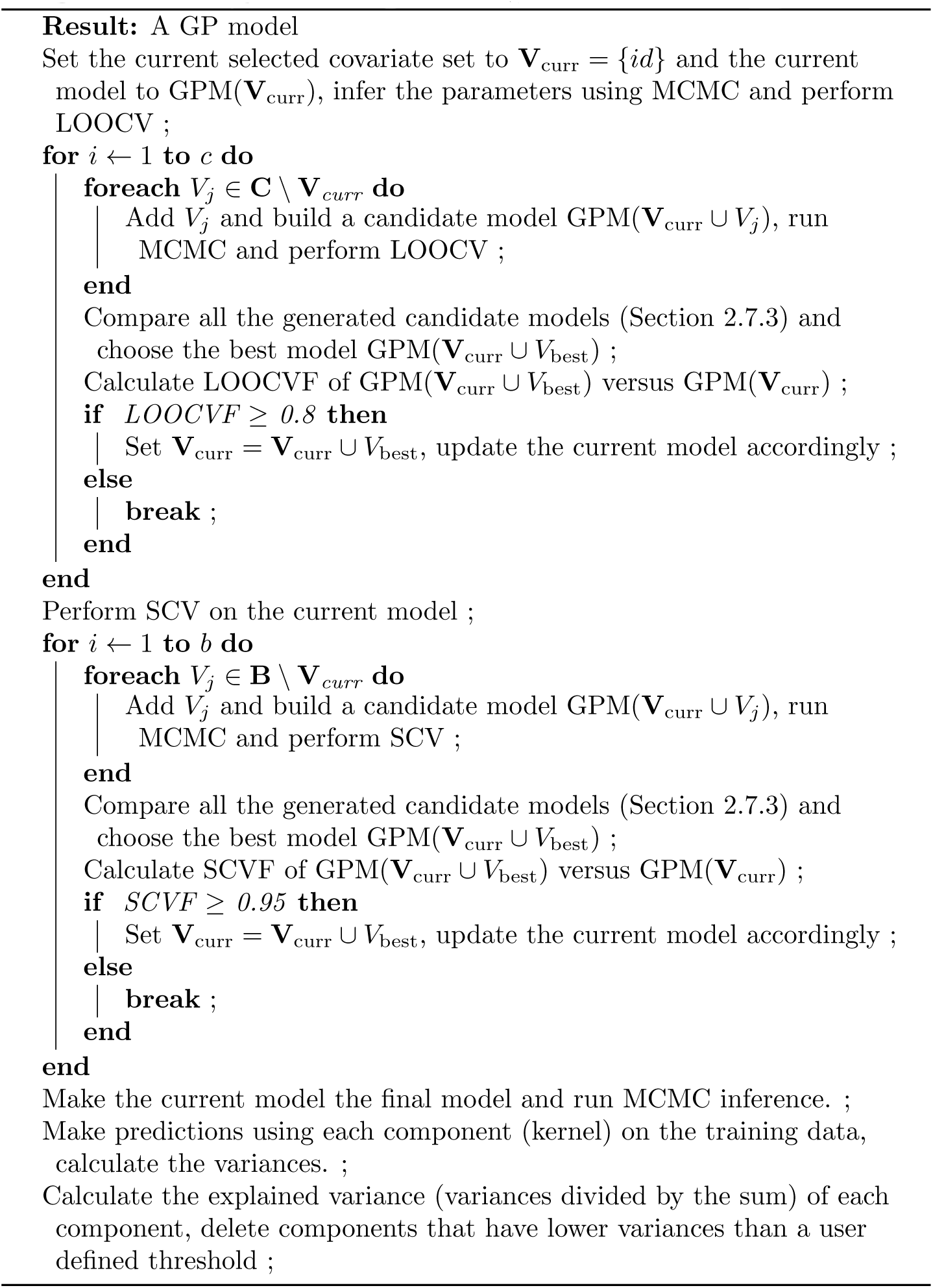
Stepwise GP regression algorithm

The algorithm tries to select covariates with reasonably large effects and the thresholds of the LOOCVF and SCVF are determined by the user (defaults are 0.8 and 0.95).

### 4 Software architecture

In many occasions more than one target variable is measured, such as in transcriptome studies using microarrays or RNA-sequencing, which means that we need to run LonGP for many target variables at the same time. Fortunately, several parts of our method can be efficiently parallelised. We designed the LonGP software so that it can be easily deployed and parallelised in a modern computing cluster with shared storage, as shown in Fig. 4. Briefly, there are three types of nodes in the physical layer. The task manager monitors the whole process and assigns different tasks to the main workers and slaves. The main workers focus on one target variable and ensure that the tasks are executed in the right order. It also informs the task manager about the parallel tasks that are available. The slaves run parallel tasks assigned by the task manager. When a main worker finishes its job, it will turn into a slave node.

### 5 Tables for simulation experiments

**Table 1.** Model selection accuracy as a function of noise variance. Table shows the number of times the correct model is identified among 100 Monte Carlo simulations.

| Generated Datasets | noise = 1 | noise = 3 | noise = 5 | noise = 8 |
| --- | --- | --- | --- | --- |
| AGPM1 | 100 | 98 | 93 | 95 |
| AGPM2 | 98 | 95 | 92 | 86 |
| AGPM3 | 96 | 95 | 93 | 91 |
| AGPM4 | 96 | 92 | 92 | 90 |
| AGPM5 | 94 | 88 | 82 | 80 |

**Table 2.** Model selection accuracy as a function of sample size. Table shows the number of times the correct model is identified among 100 Monte Carlo simulations.

| Generated Datasets | 10 cases and 10 controls | 20 cases and 20 controls | 30 cases and 30 controls | 40 cases and 40 controls |
| --- | --- | --- | --- | --- |
| AGPM1 | 90 | 98 | 94 | 98 |
| AGPM2 | 71 | 95 | 93 | 95 |
| AGPM3 | 85 | 95 | 95 | 97 |
| AGPM4 | 85 | 92 | 94 | 96 |
| AGPM5 | 71 | 88 | 96 | 97 |

**Table 3.** Model selection accuracy as a function of sampling time points. Table shows the number of times the correct model is identified among 100 Monte Carlo simulations.

| Generated Datasets | 2 months | 3 months | 4 months | 6 months |
| --- | --- | --- | --- | --- |
| AGPM1 | 97 | 98 | 94 | 96 |
| AGPM2 | 95 | 95 | 88 | 85 |
| AGPM3 | 97 | 95 | 91 | 93 |
| AGPM4 | 96 | 92 | 86 | 86 |
| AGPM5 | 94 | 88 | 87 | 86 |

**Table 4.** Inclusion of *diseaseAge* in the final model for simulated data with 20 cases and 20 controls, noise variance 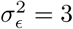 and 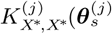 samples taken every 3 months. Table shows the number of times the *diseaseAge* covariate is included in the inferred model among 100 Monte Carlo simulations.

| Generated Datasets | *diseaseAge* detected | *diseaseAge* not detected |
| --- | --- | --- |
| AGPM1 | 0 | 100 |
| AGPM2 | 1 | 99 |
| AGPM3 | 0 | 100 |
| AGPM4 | 97 | 3 |
| AGPM5 | 97 | 3 |

**Table 5.**
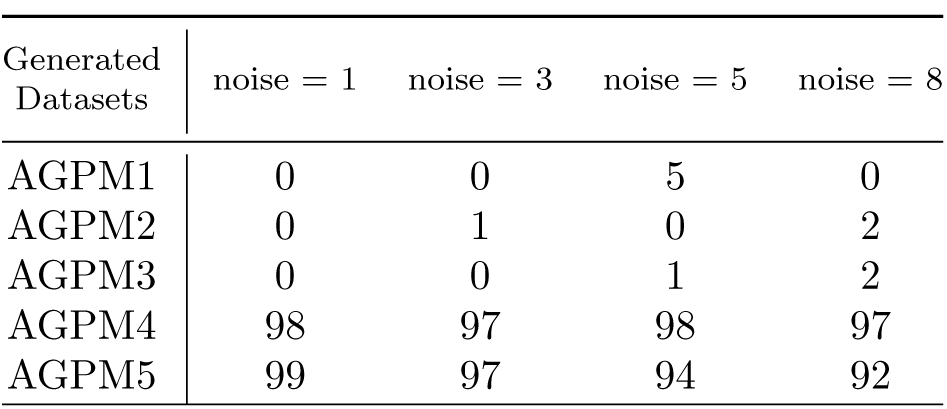
Inclusion of *diseaseAge* in the final model as a function of noise variance. Table shows the number of times the *diseaseAge* covariate is included in the inferred model among 100 Monte Carlo simulations.

| Generated Datasets | noise = 1 | noise = 3 | noise = 5 | noise = 8 |
| --- | --- | --- | --- | --- |
| AGPM1 | 0 | 0 | 5 | 0 |
| AGPM2 | 0 | 1 | 0 | 2 |
| AGPM3 | 0 | 0 | 1 | 2 |
| AGPM4 | 98 | 97 | 98 | 97 |
| AGPM5 | 99 | 97 | 94 | 92 |

**Table 6.**
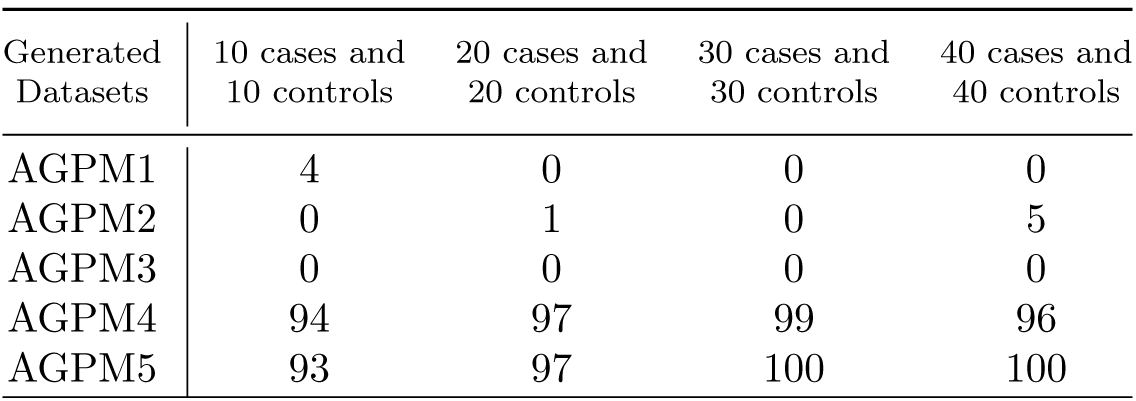
Inclusion of *diseaseAge* in the final model as a function of sample size. Table shows the number of times the *diseaseAge* covariate is included in the inferred model among 100 Monte Carlo simulations.

| Generated Datasets | 10 cases and 10 controls | 20 cases and 20 controls | 30 cases and 30 controls | 40 cases and 40 controls |
| --- | --- | --- | --- | --- |
| AGPM1 | 4 | 0 | 0 | 0 |
| AGPM2 | 0 | 1 | 0 | 5 |
| AGPM3 | 0 | 0 | 0 | 0 |
| AGPM4 | 94 | 97 | 99 | 96 |
| AGPM5 | 93 | 97 | 100 | 100 |

**Table 7.** Inclusion of *diseaseAge* in the final model as a function of sampling time points. Table shows the number of times the *diseaseAge* covariate is included in the inferred model among 100 Monte Carlo simulations.

| Generated Datasets | 2 months | 3 months | 4 months | 6 months |
| --- | --- | --- | --- | --- |
| AGPM1 | 0 | 0 | 0 | 0 |
| AGPM2 | 0 | 1 | 3 | 4 |
| AGPM3 | 0 | 0 | 1 | 1 |
| AGPM4 | 100 | 97 | 94 | 92 |
| AGPM5 | 98 | 97 | 94 | 92 |

## Appendices

- Supplementary File 1
- Supplementary File 2

Full result tables in xls format can be downloaded from: http://research.cs.aalto.fi/csb/software/longp/

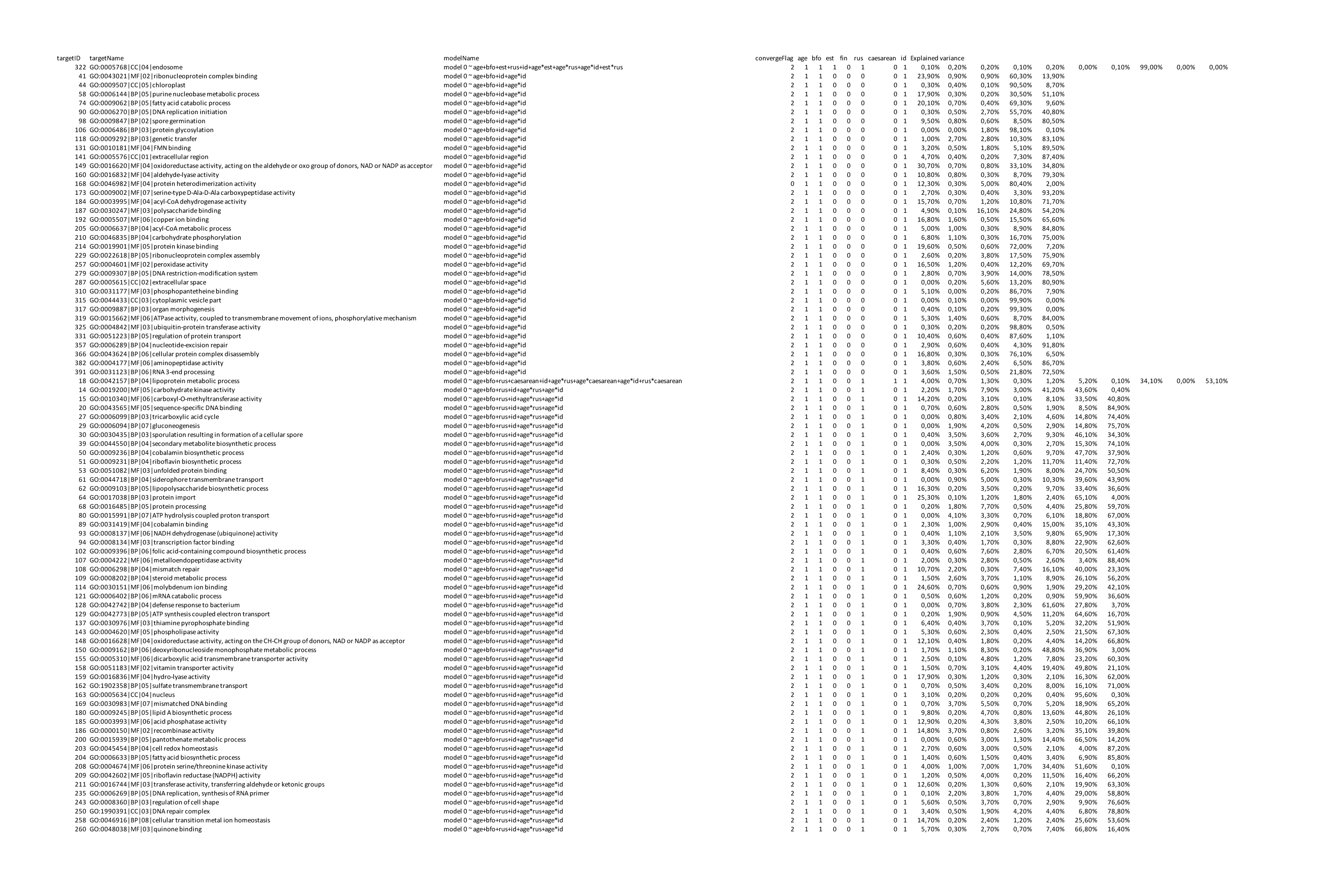
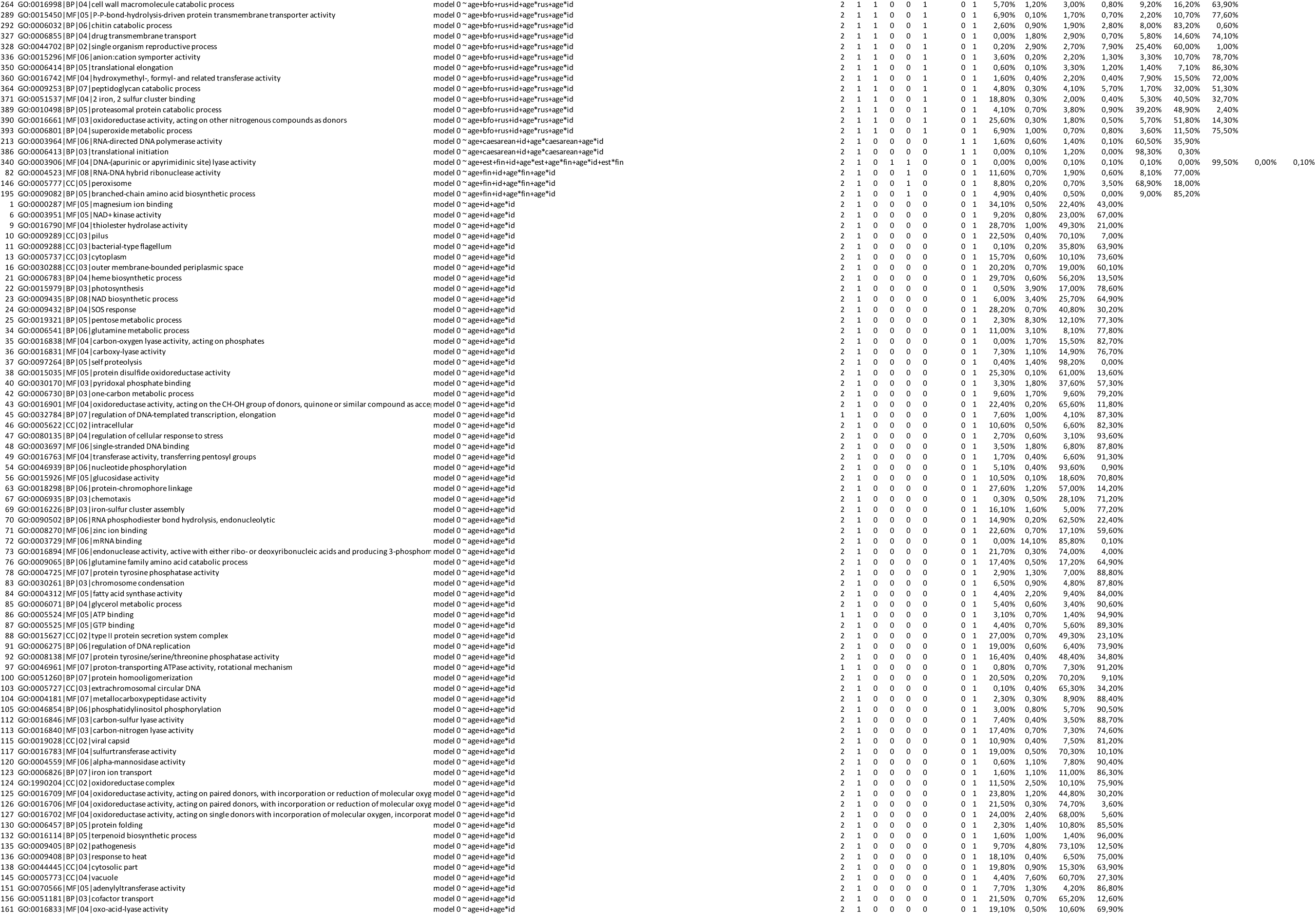
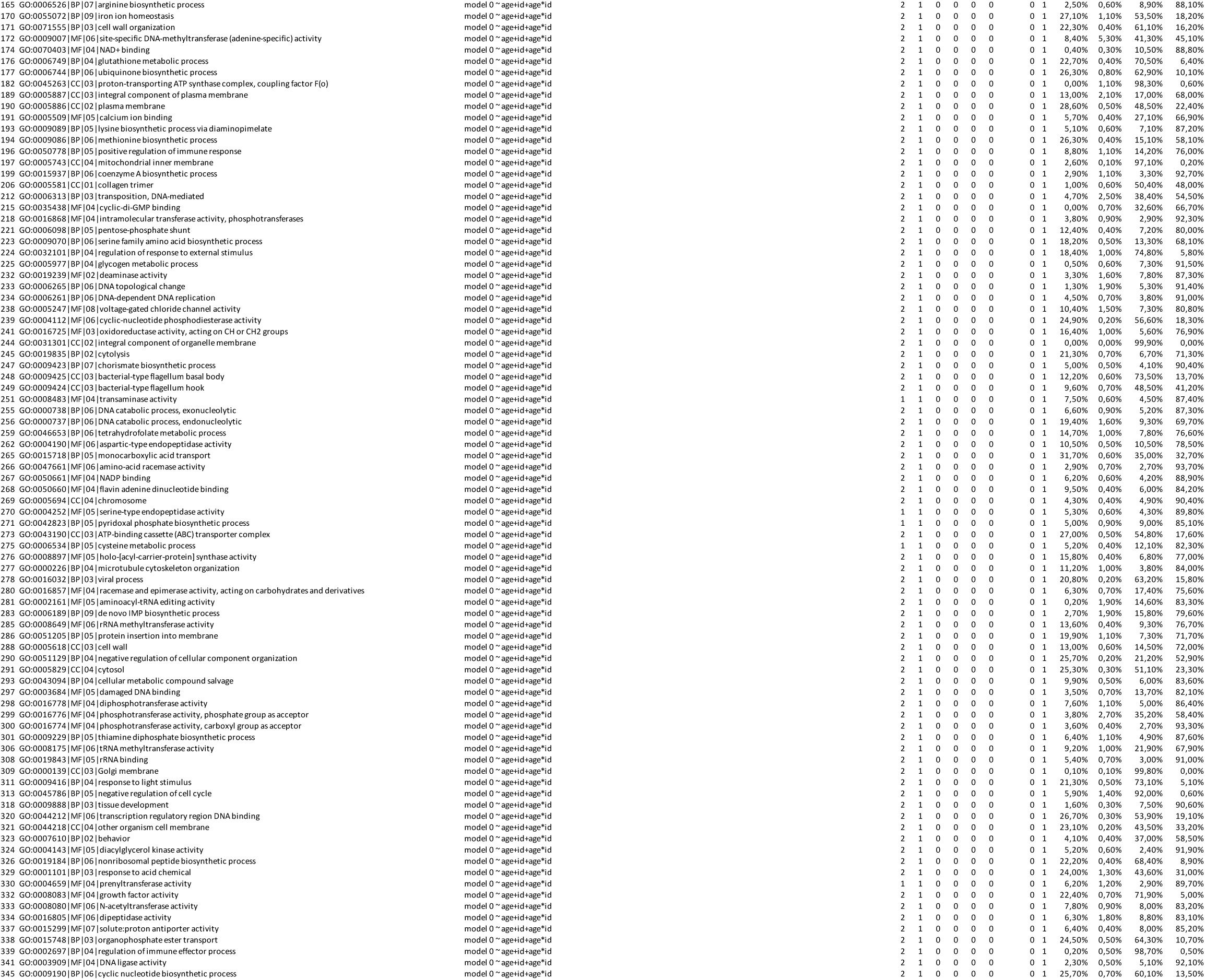
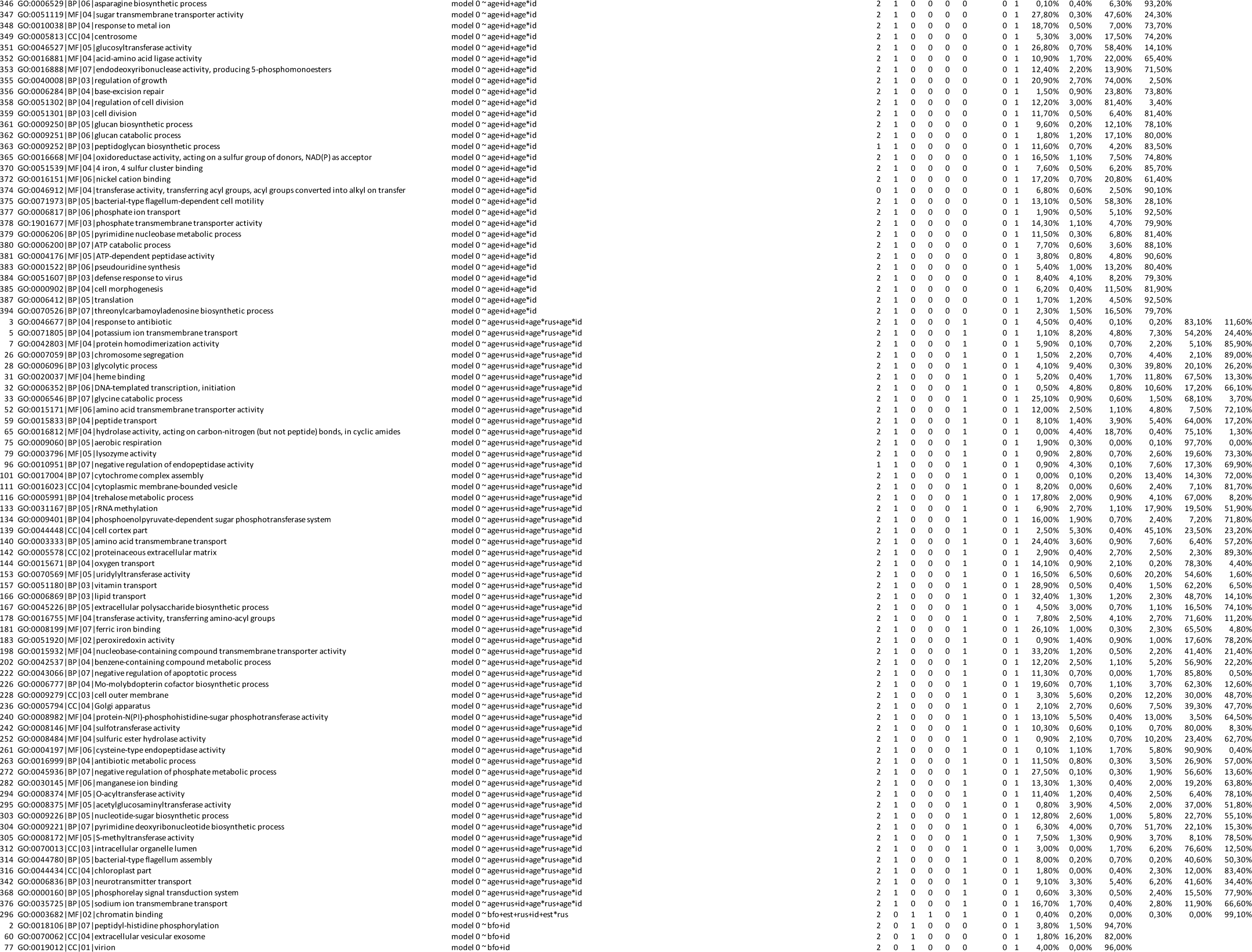
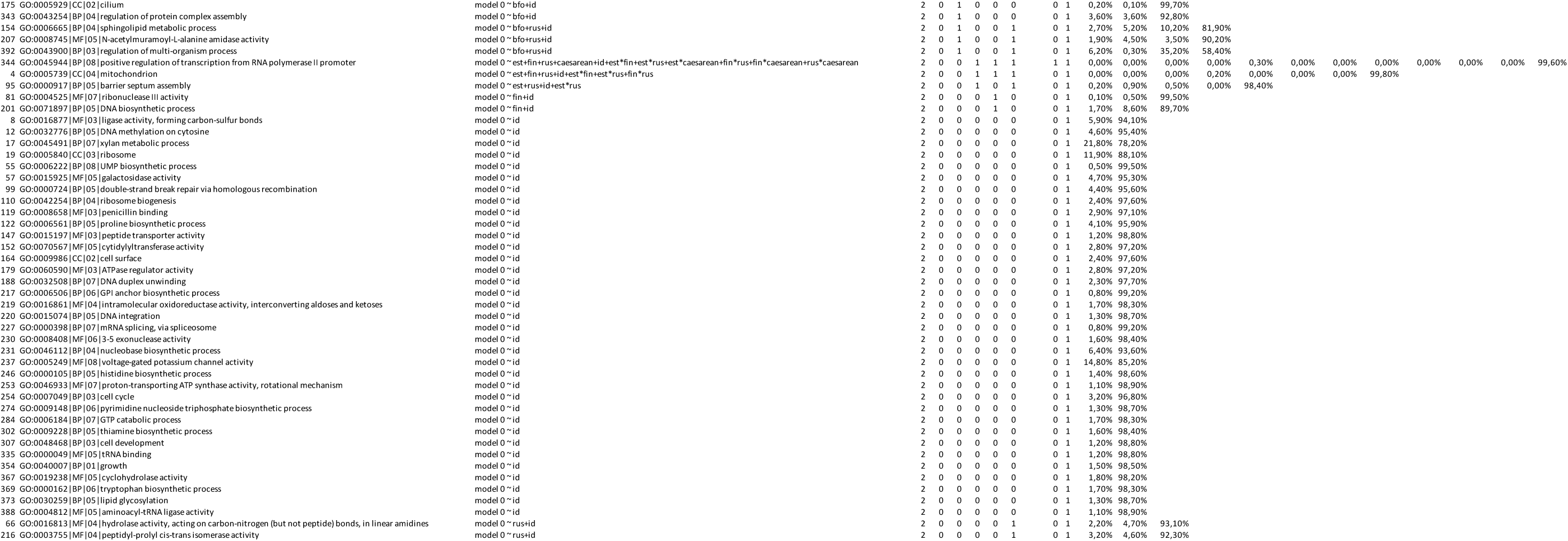
Supplementary File 1

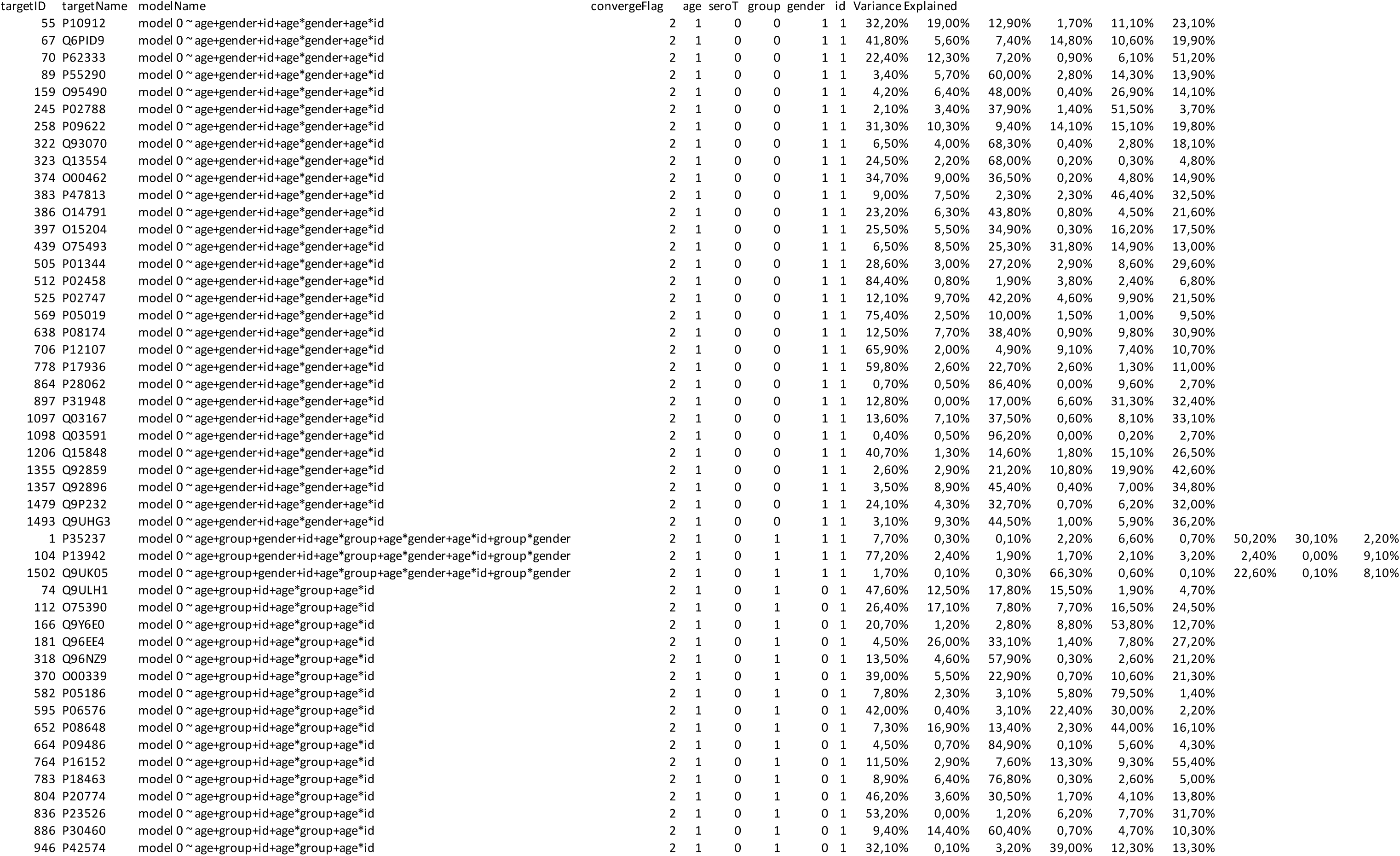
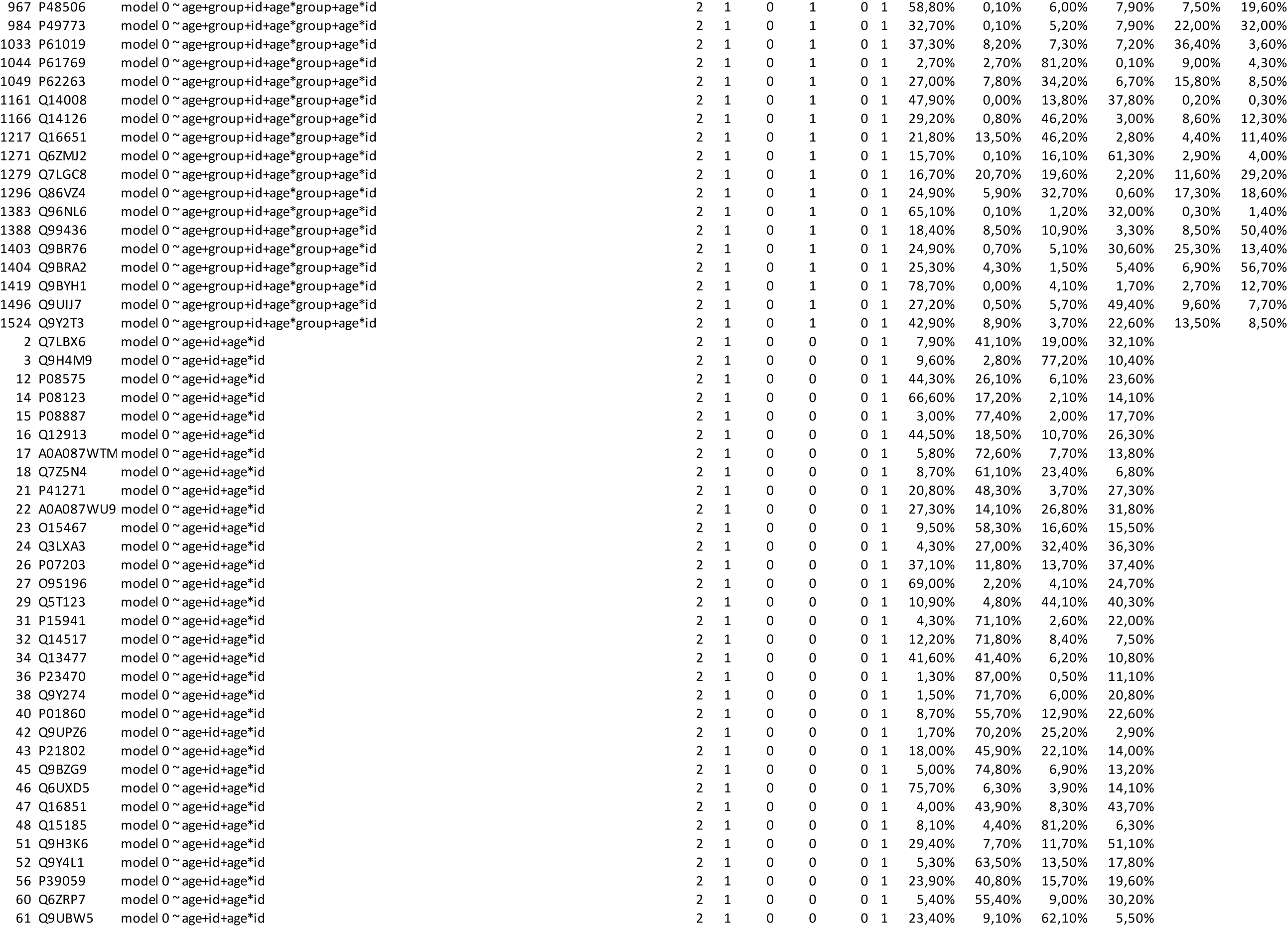
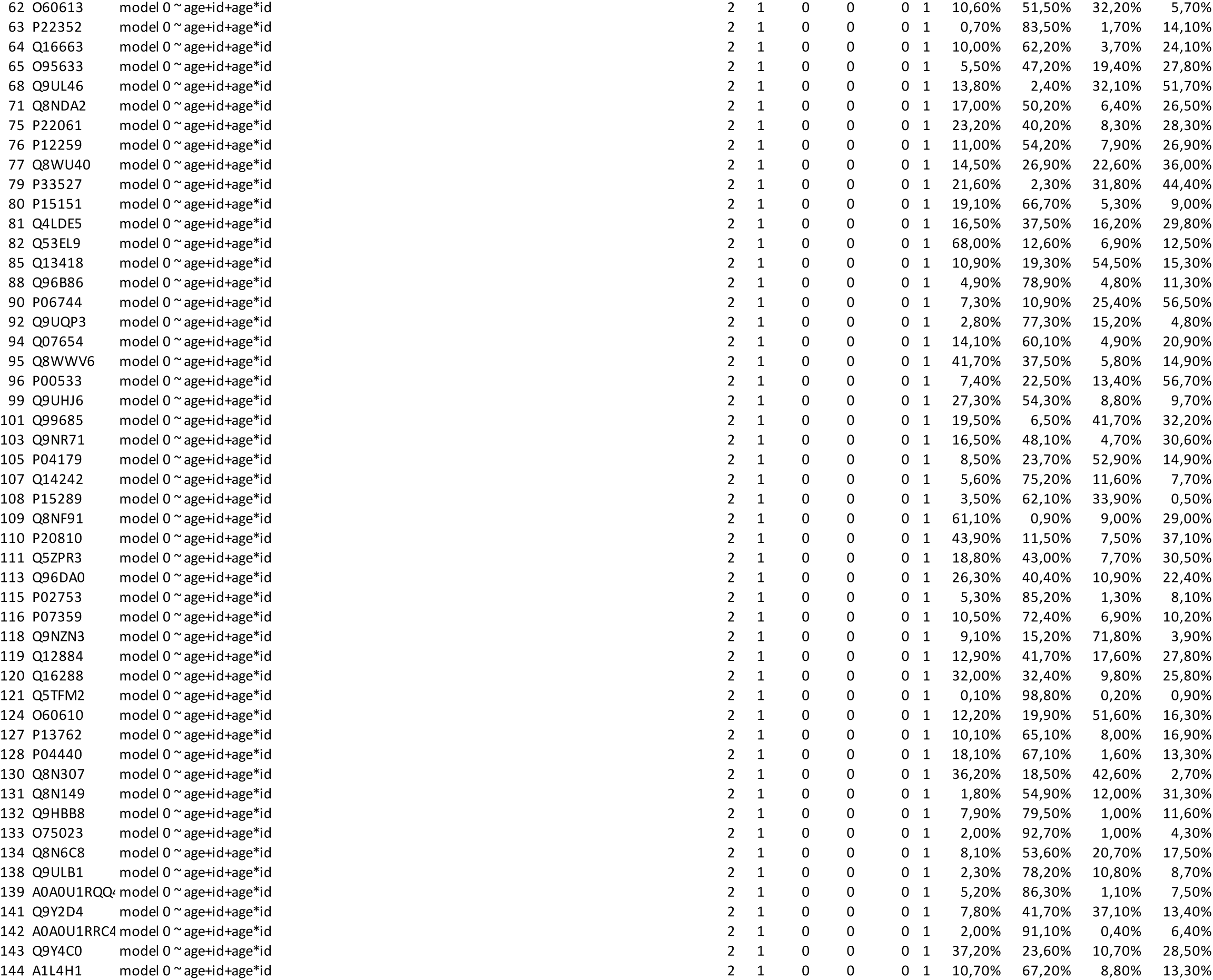
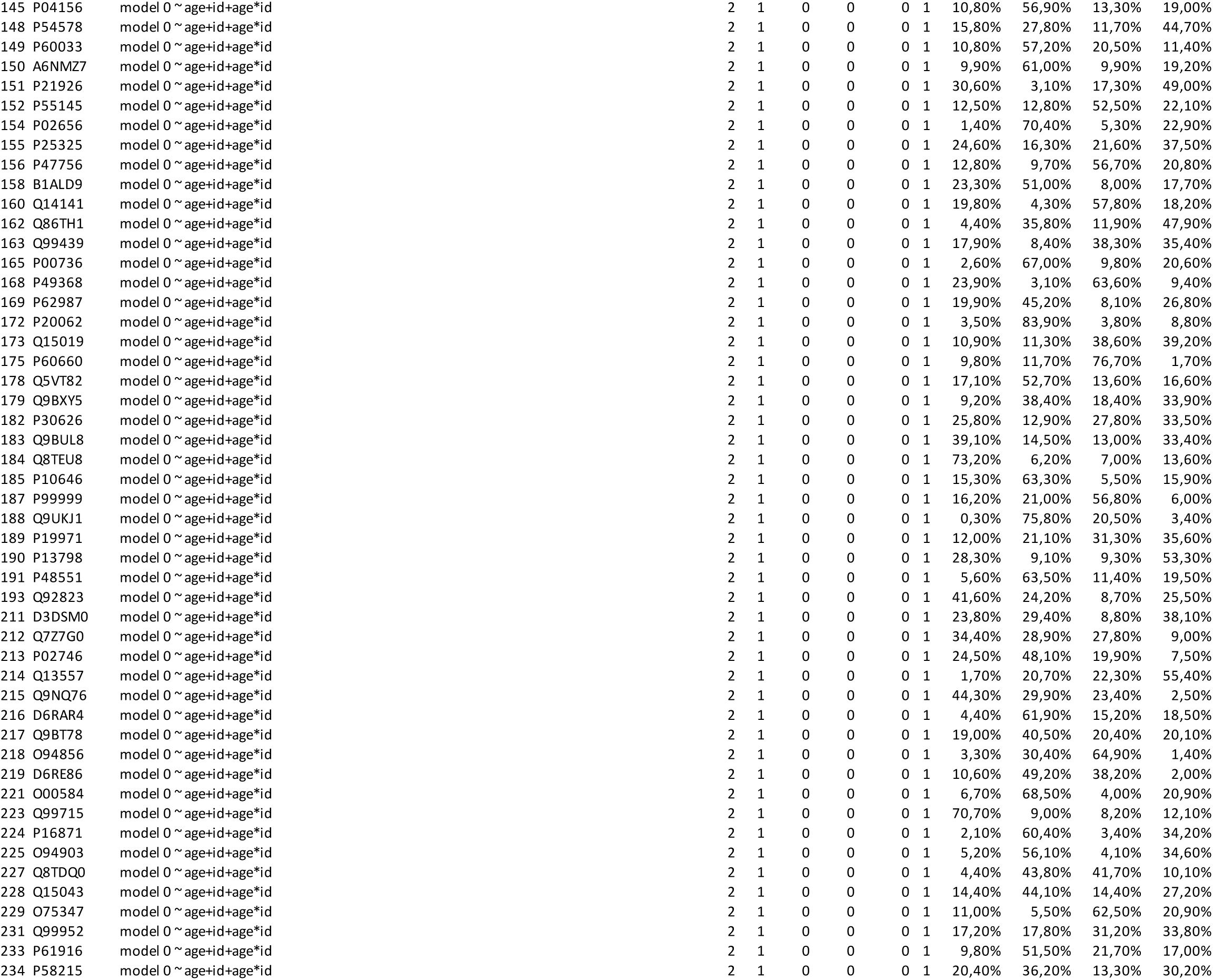
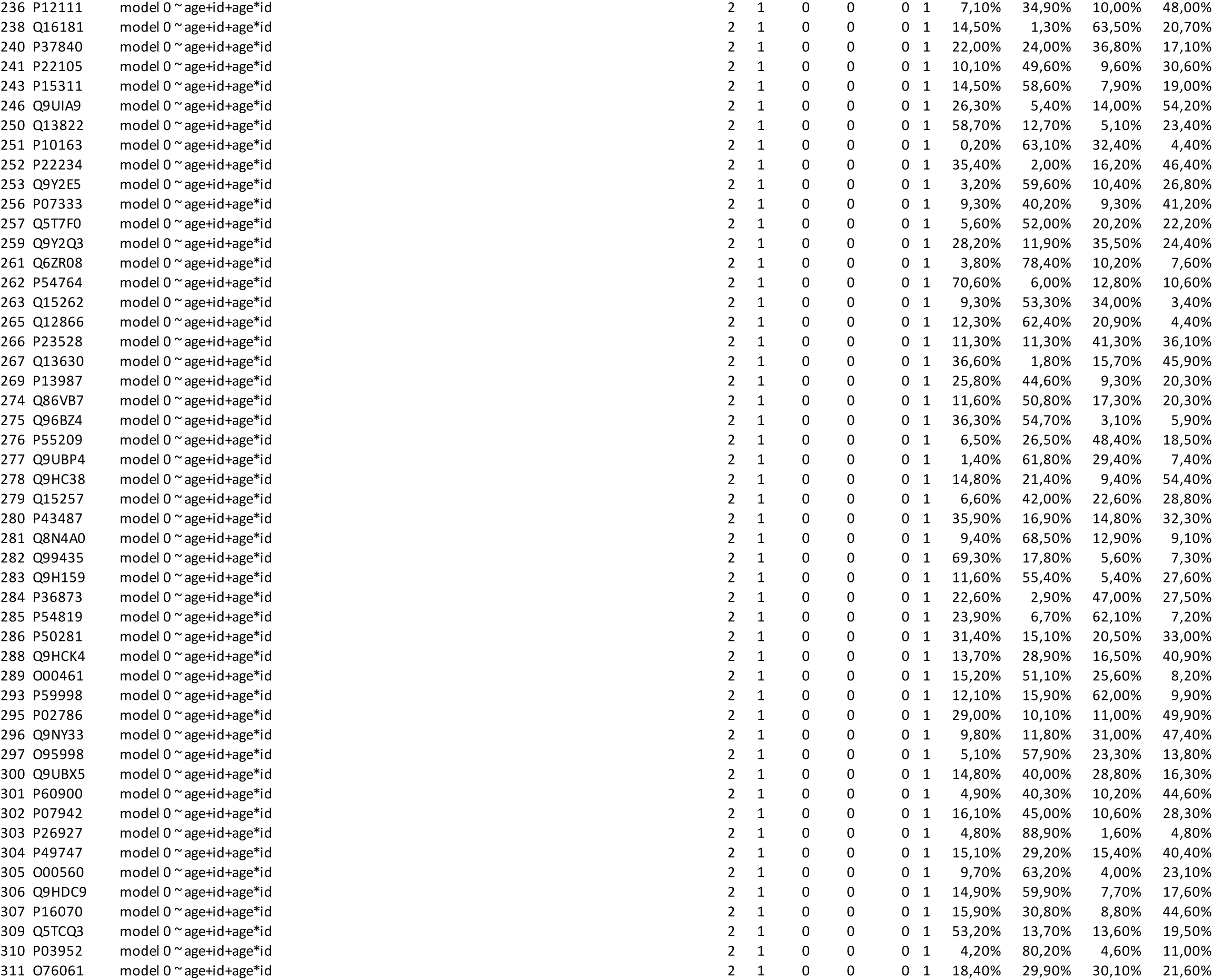
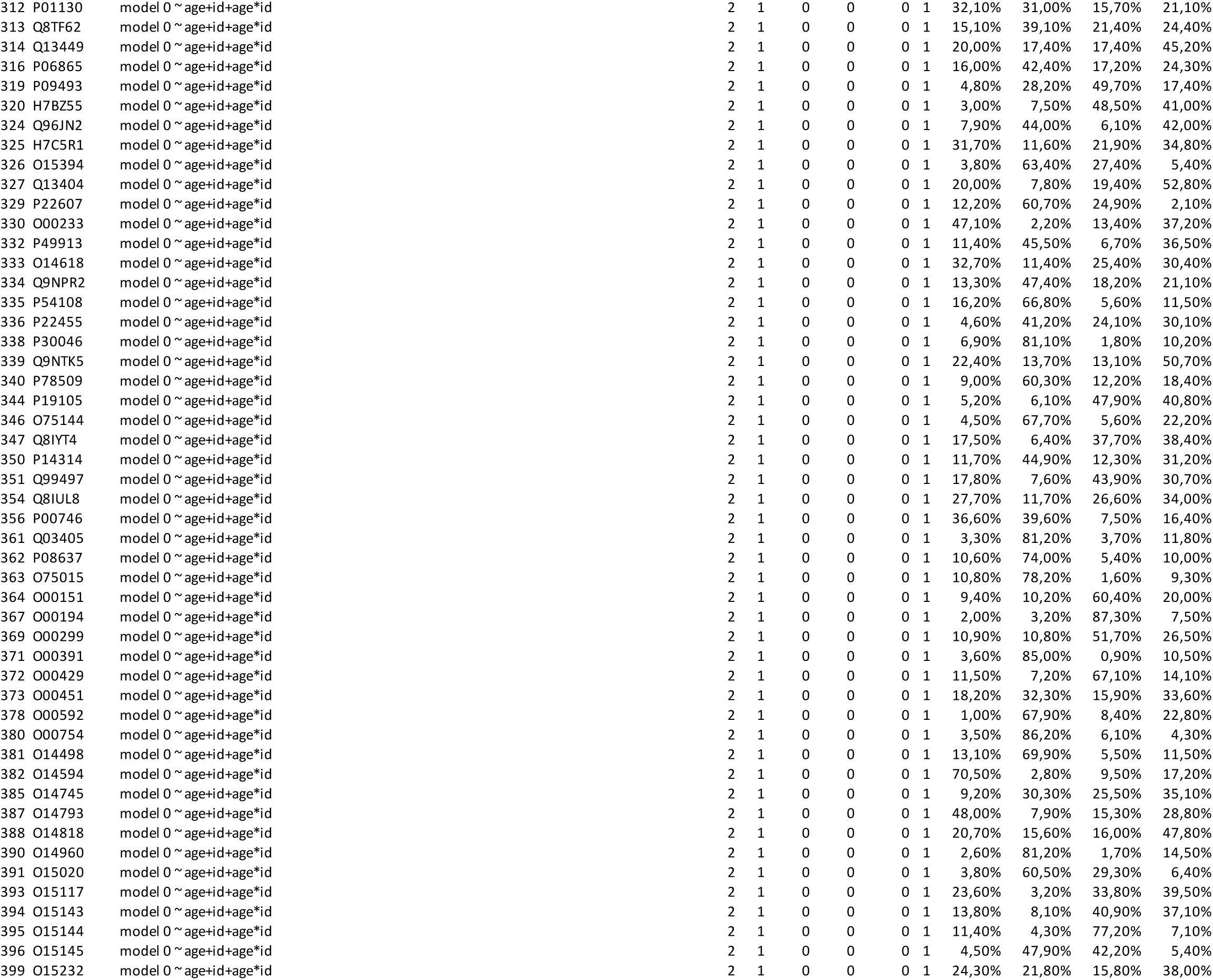
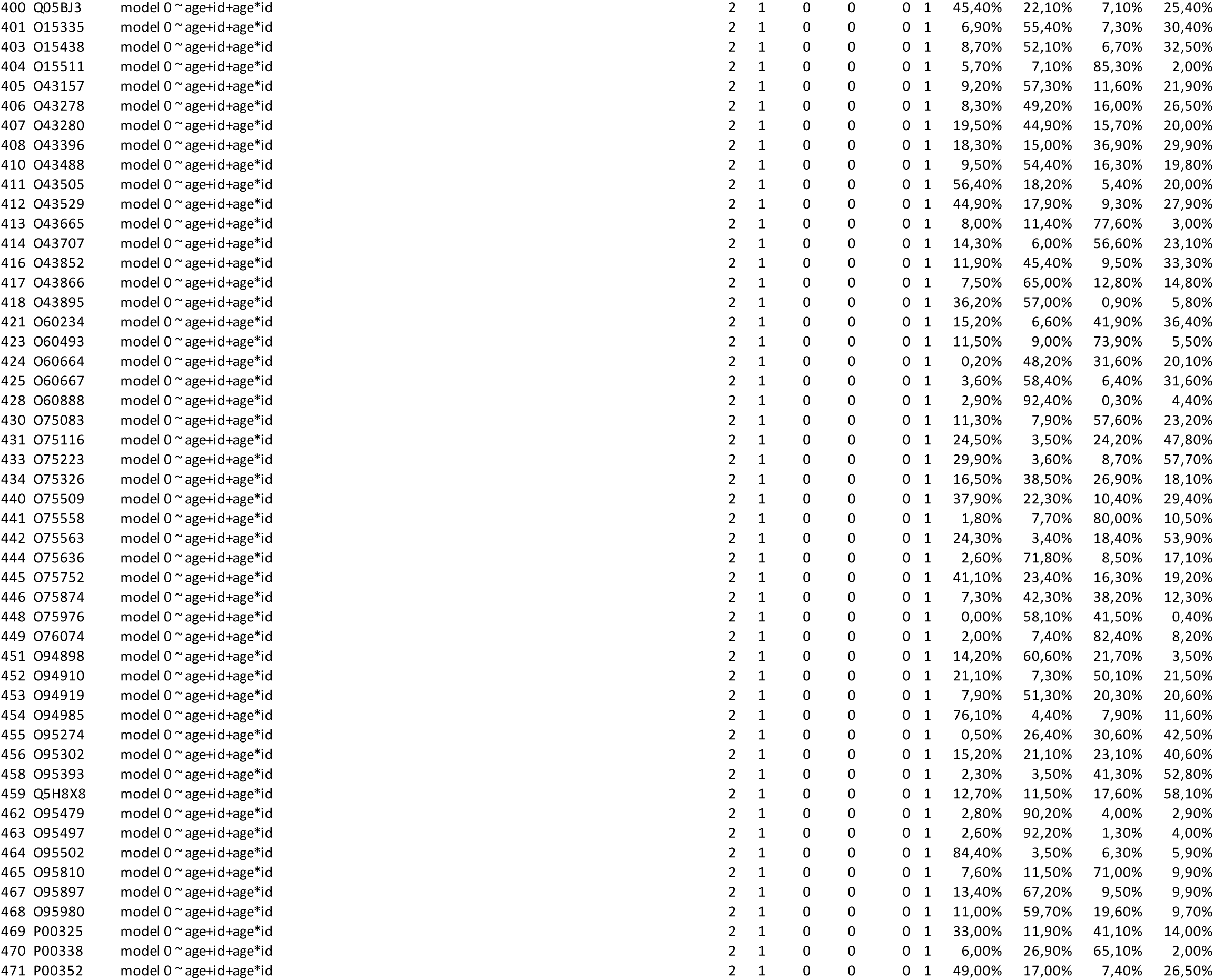
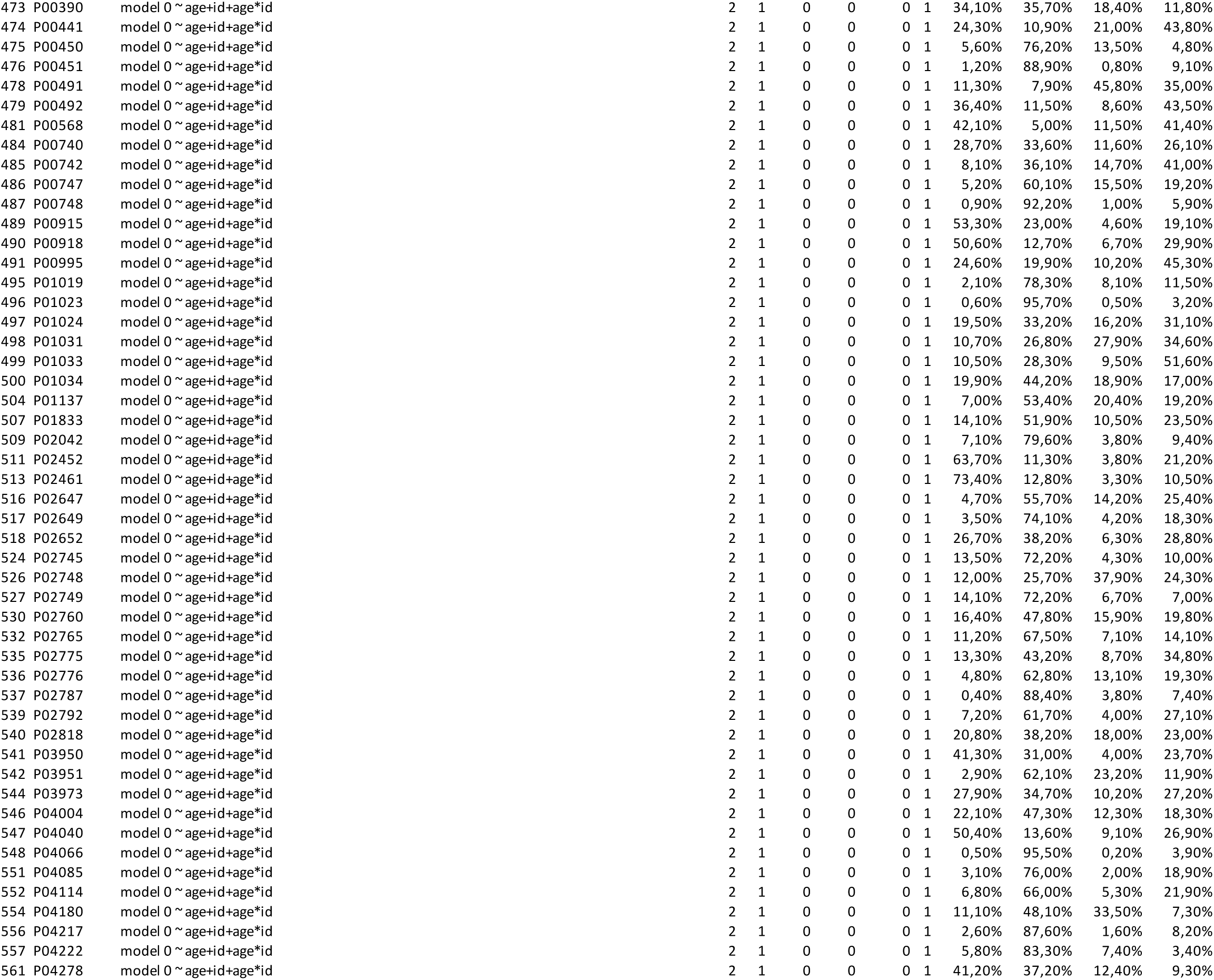
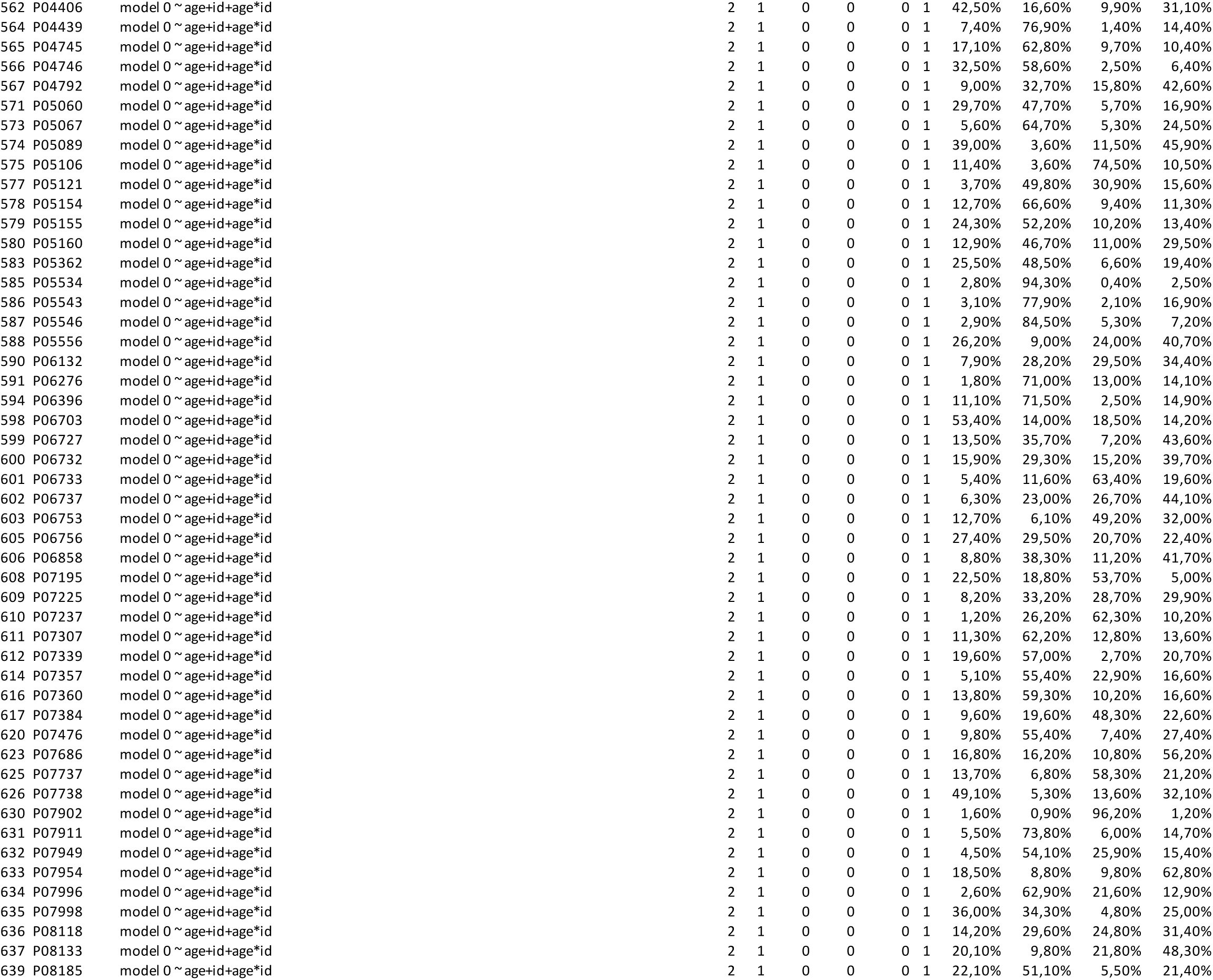
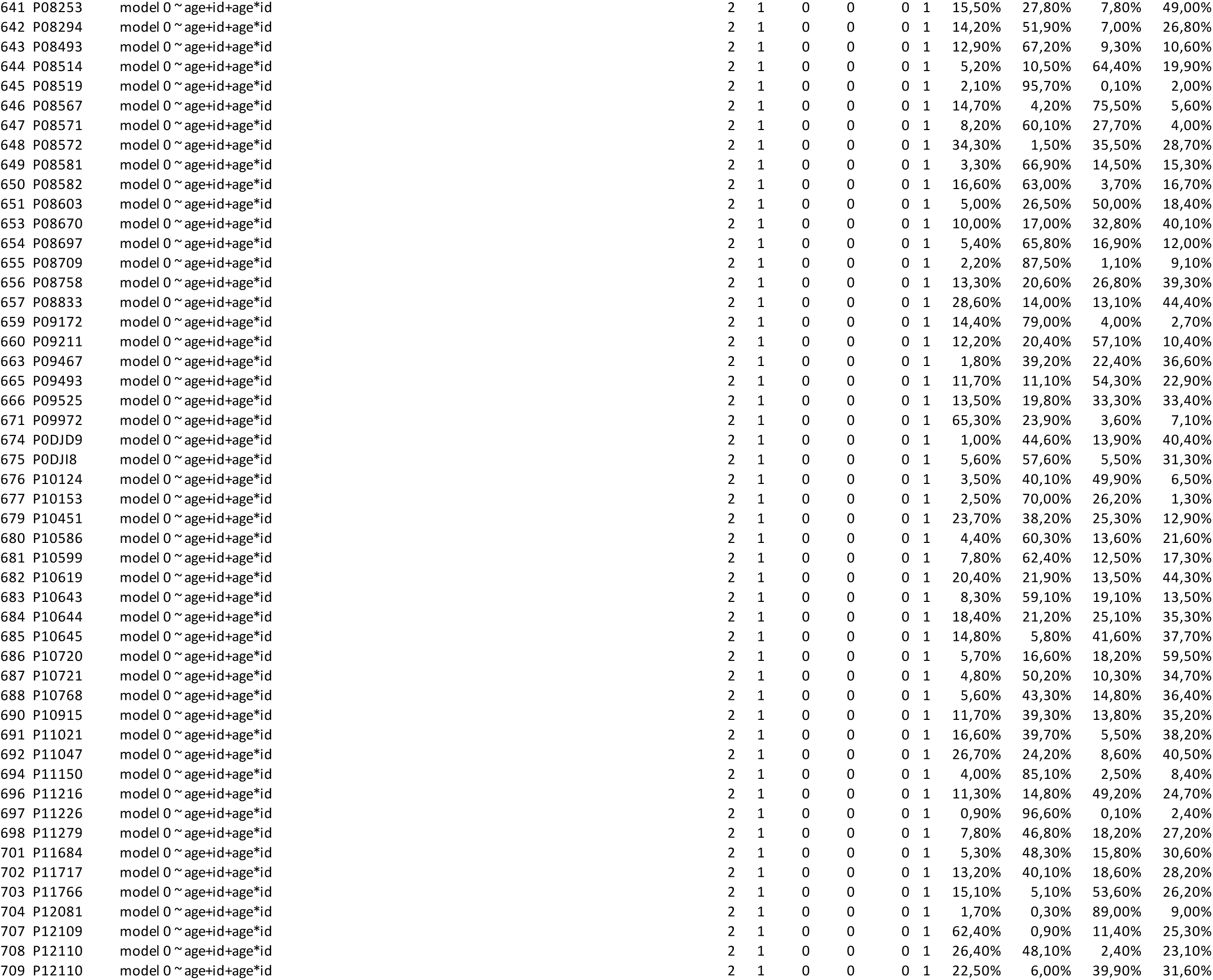
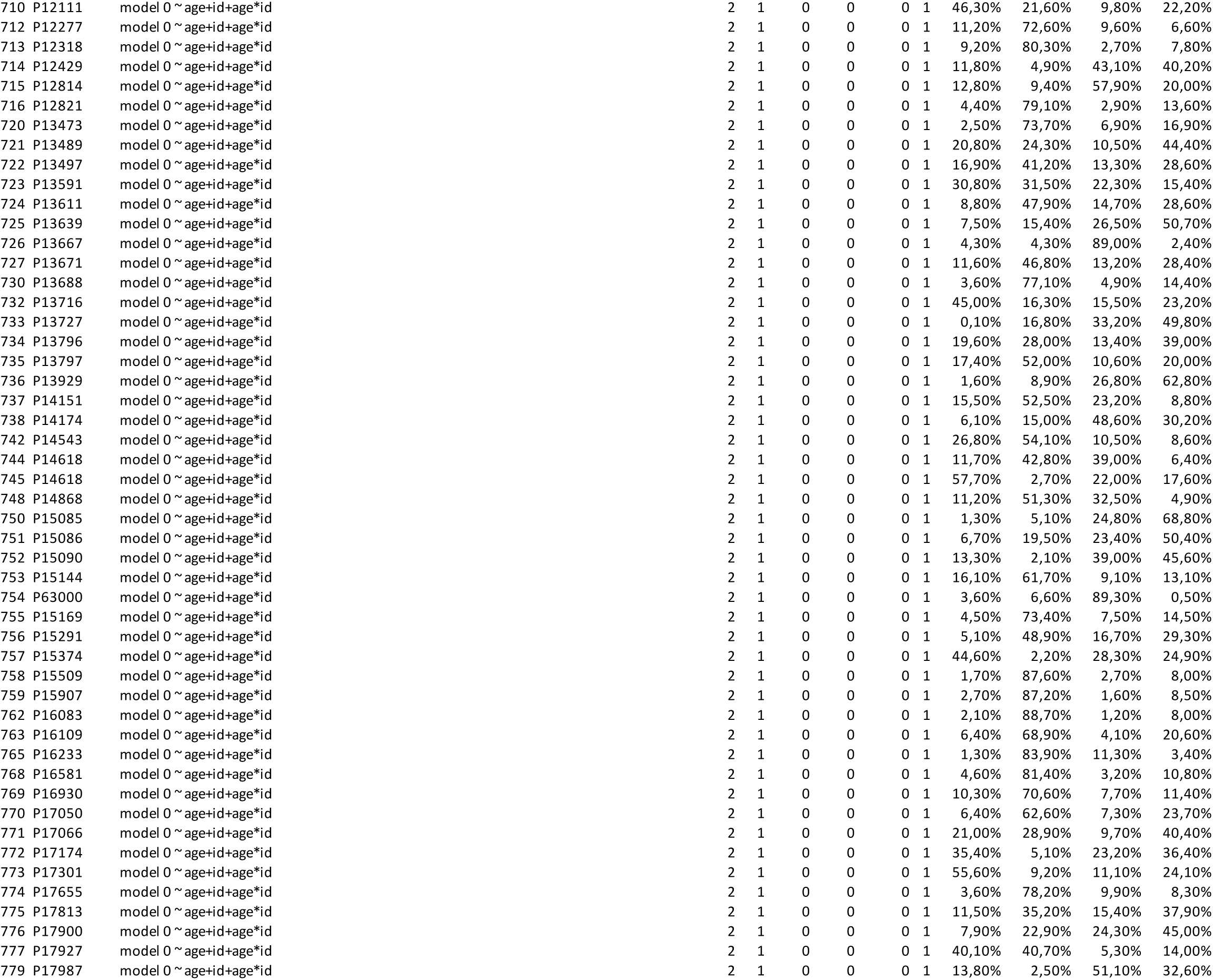
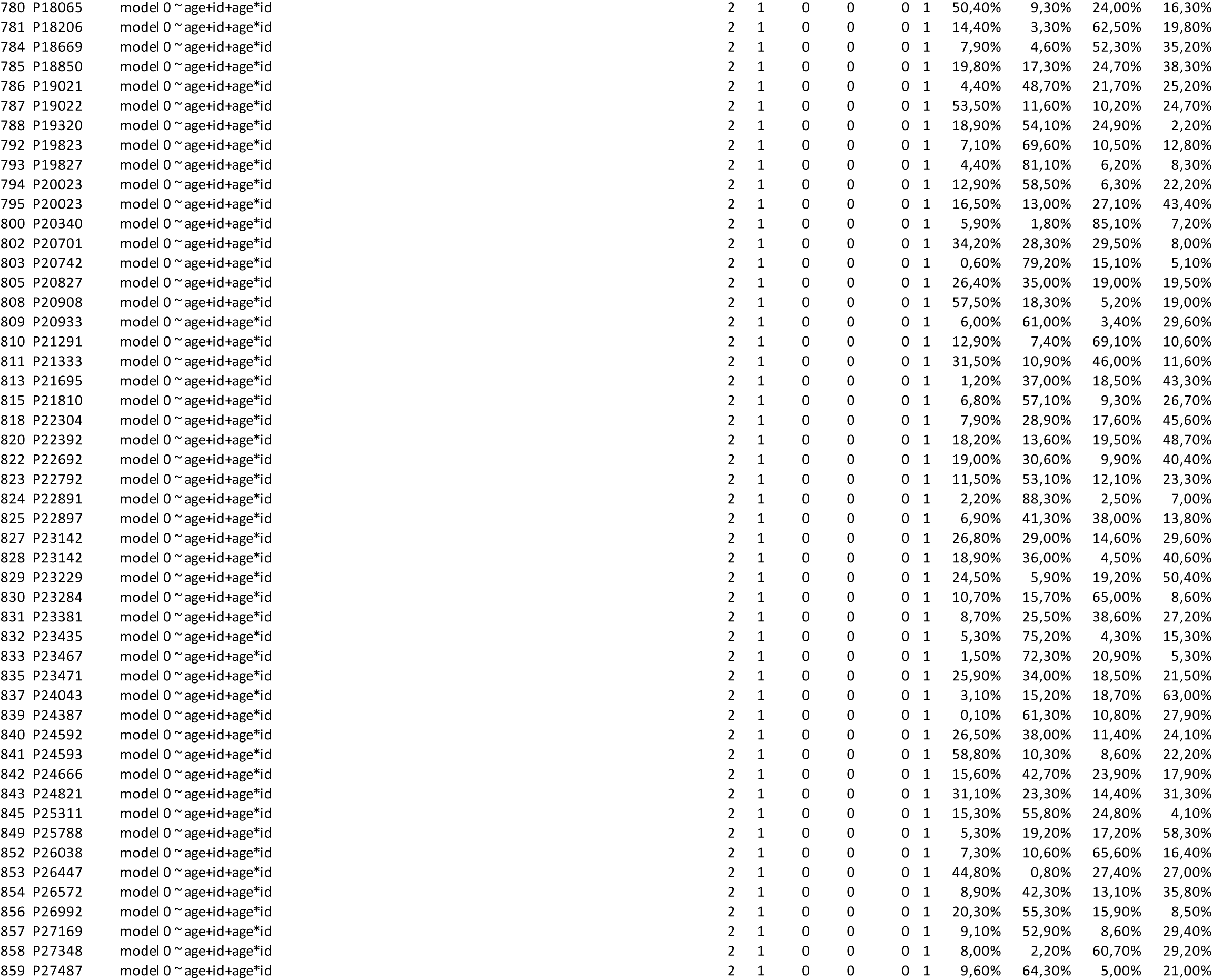
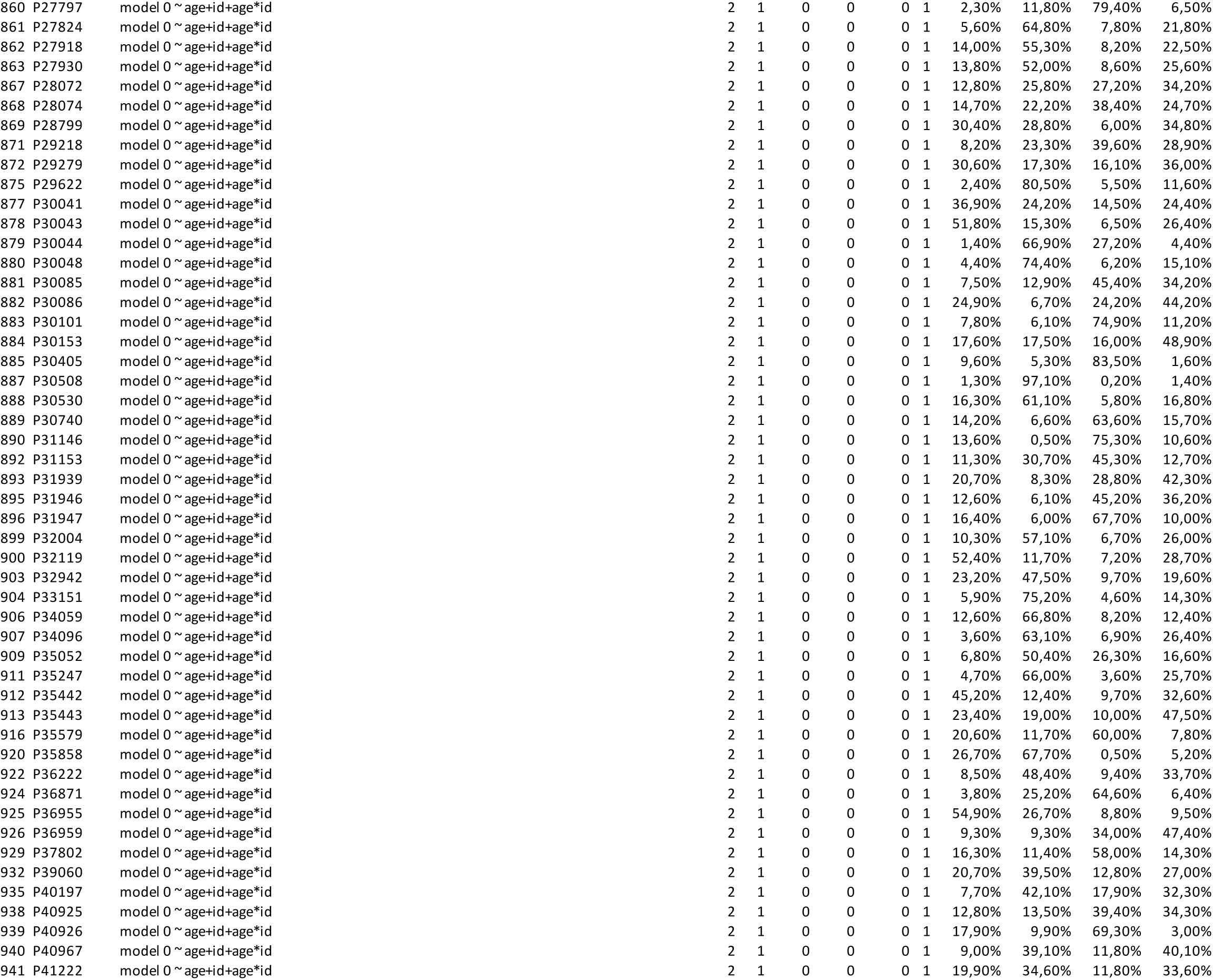
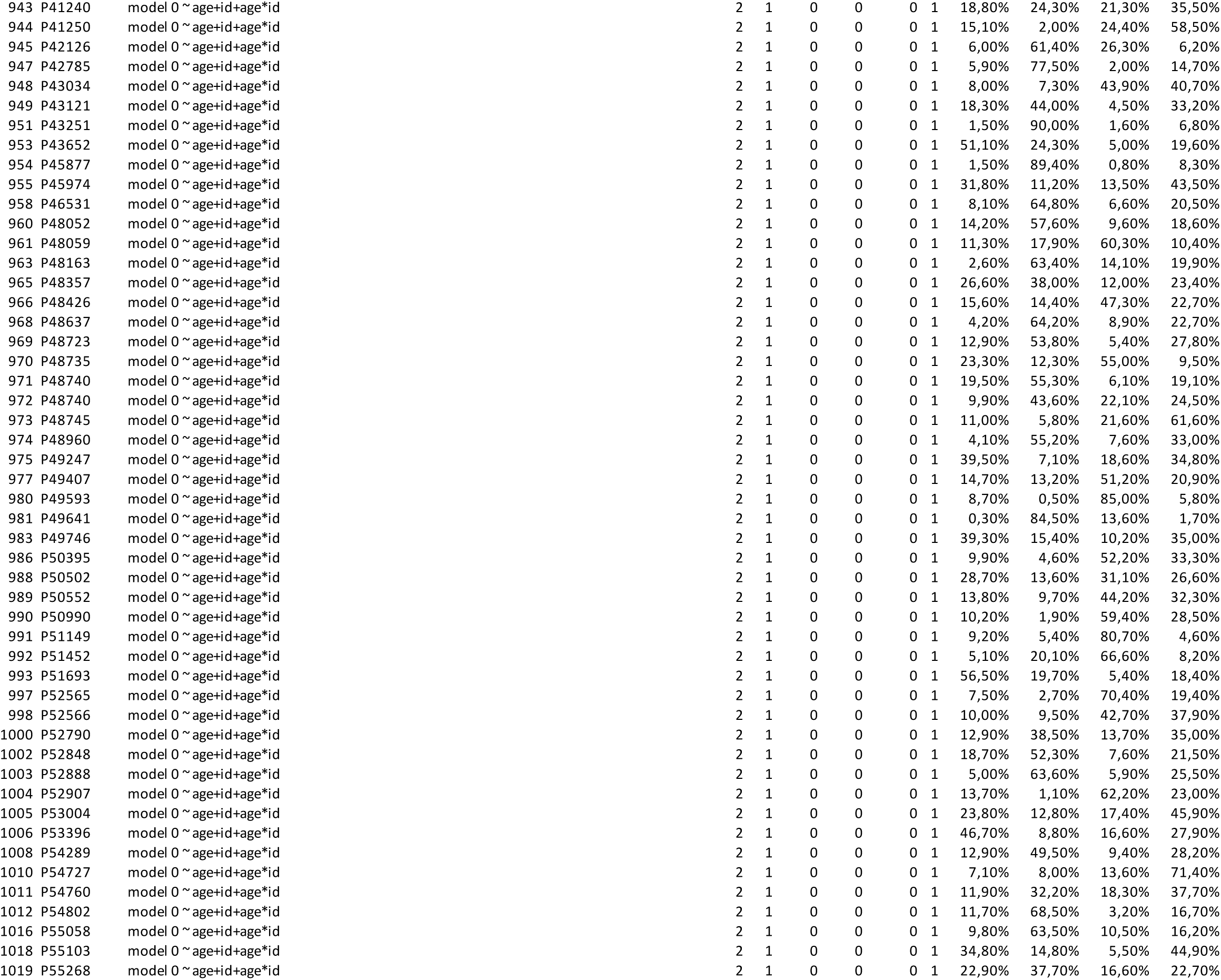
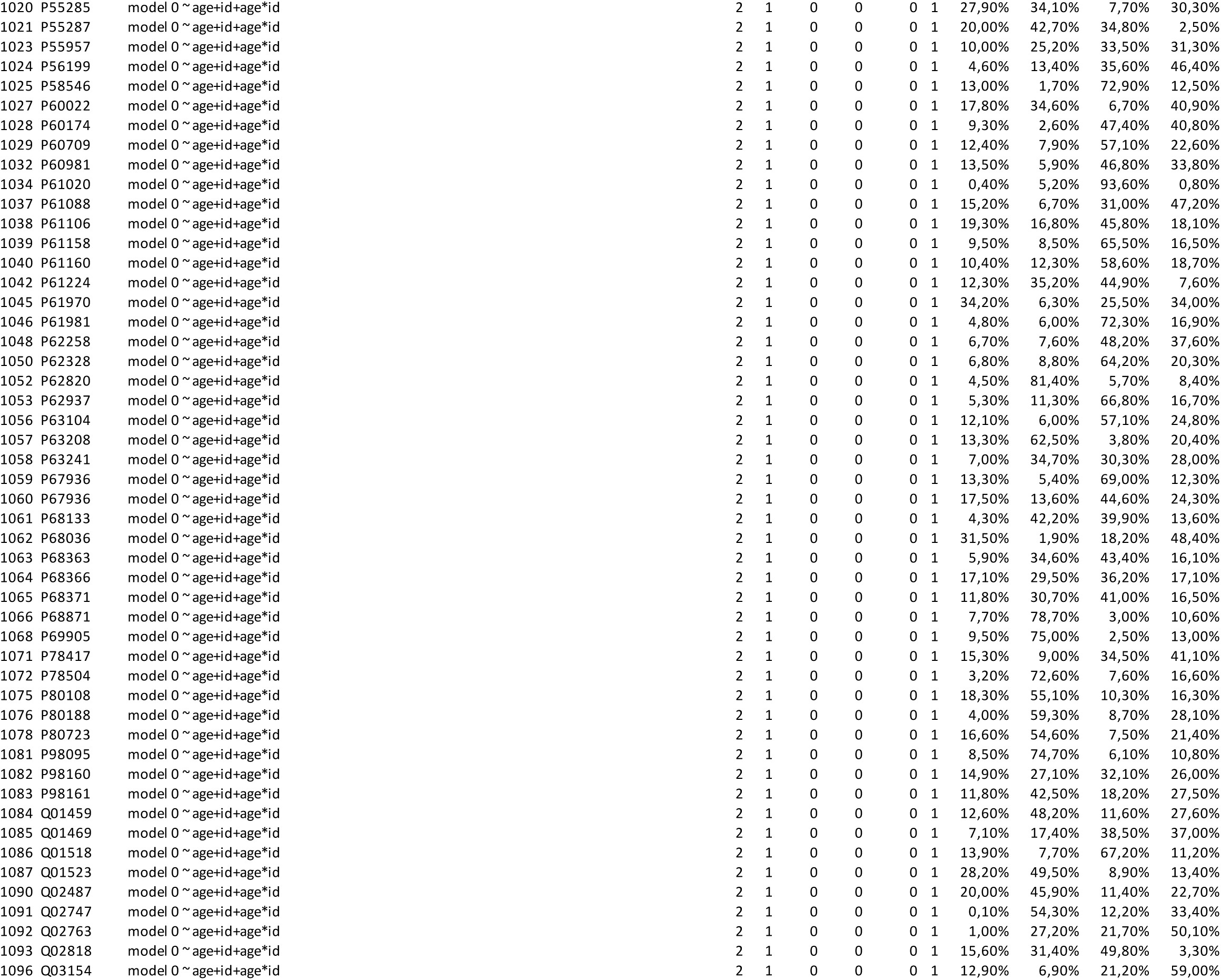
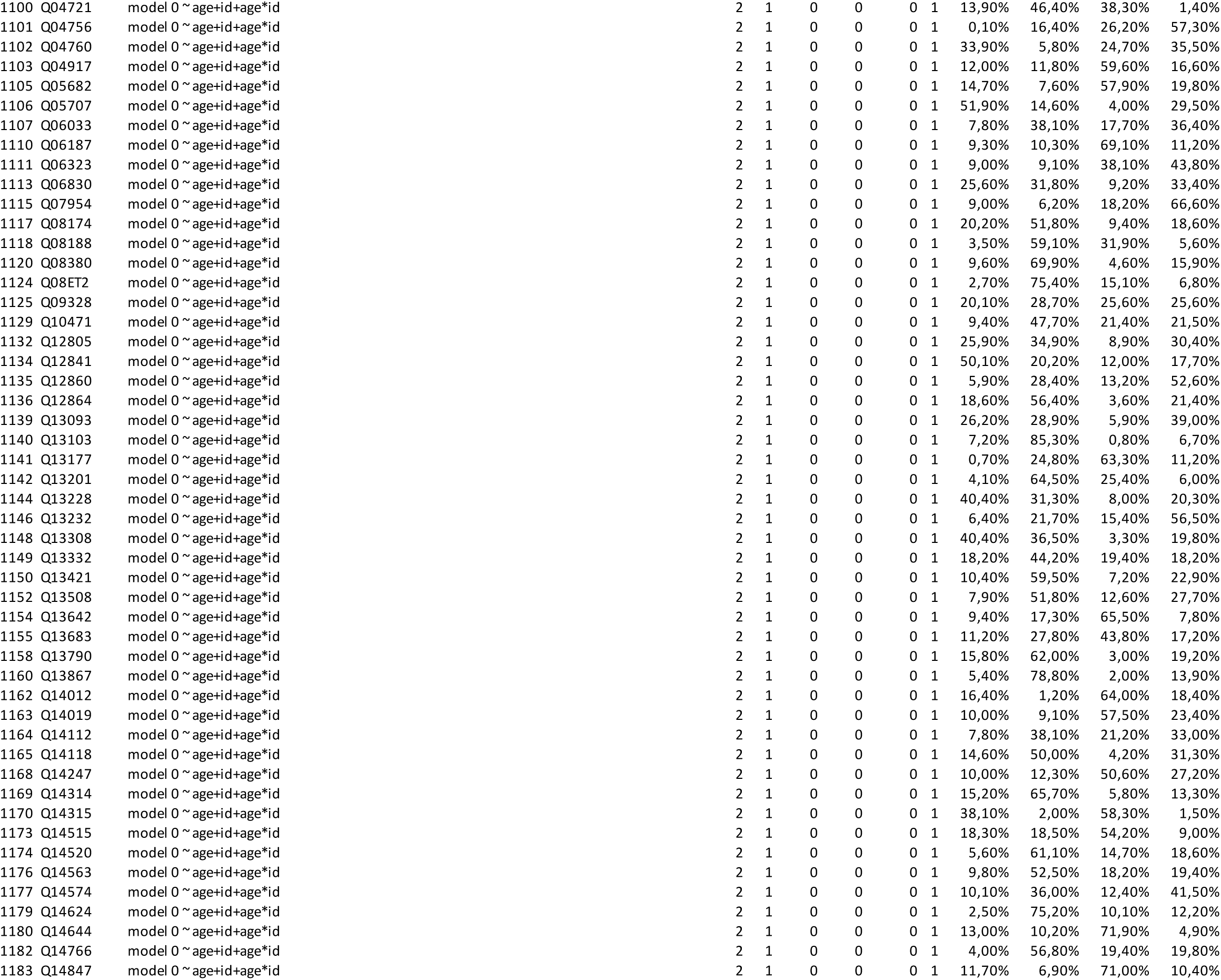
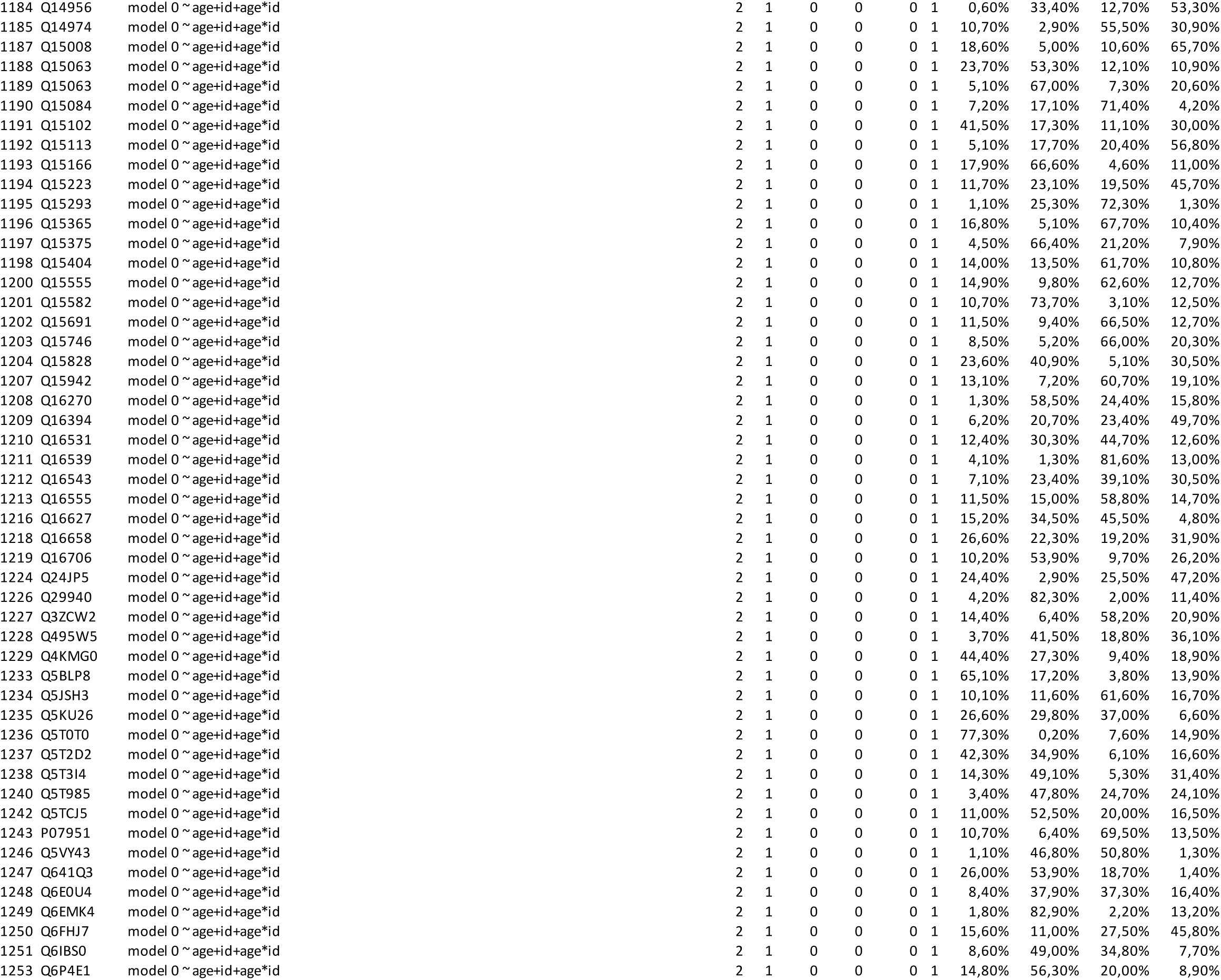
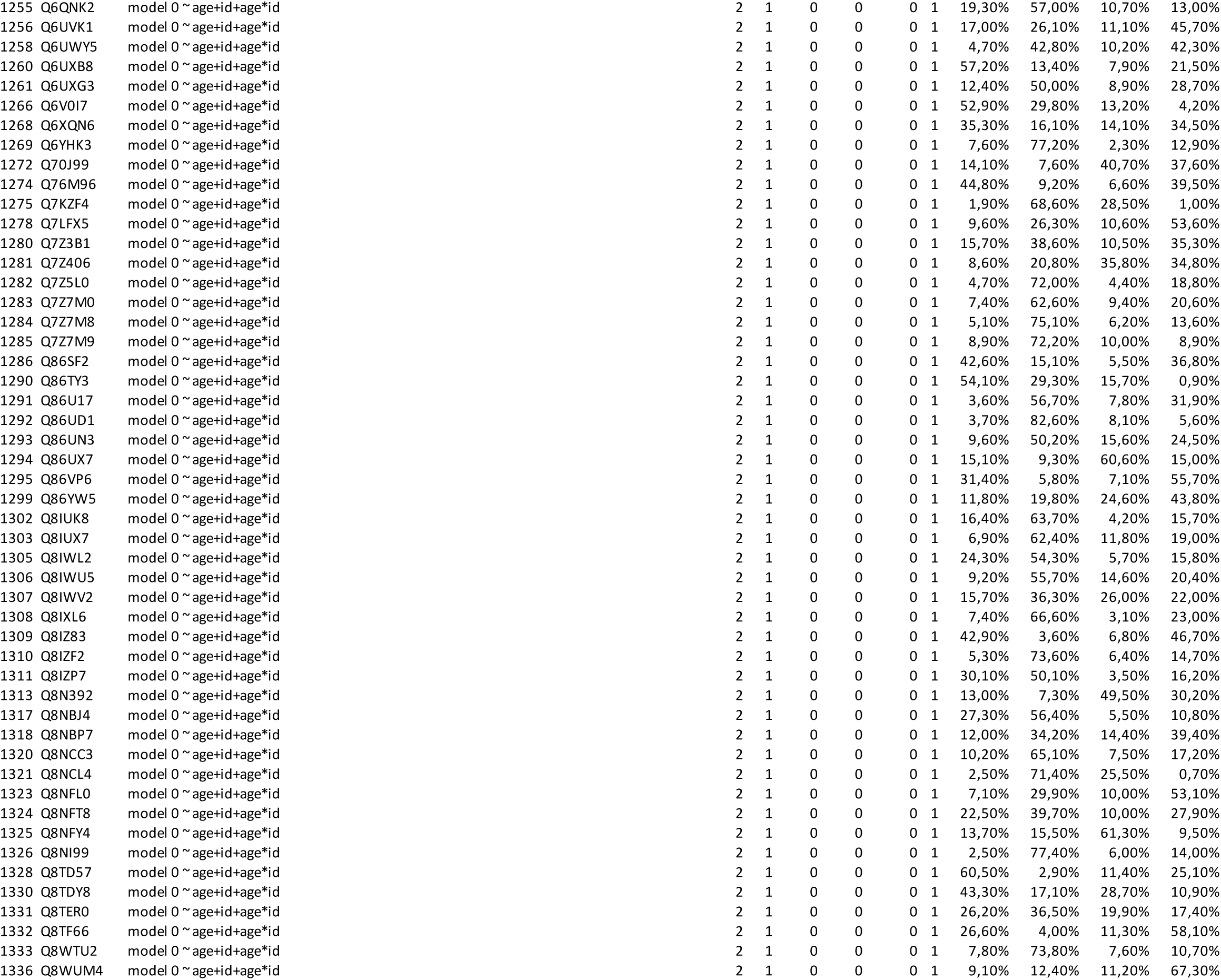
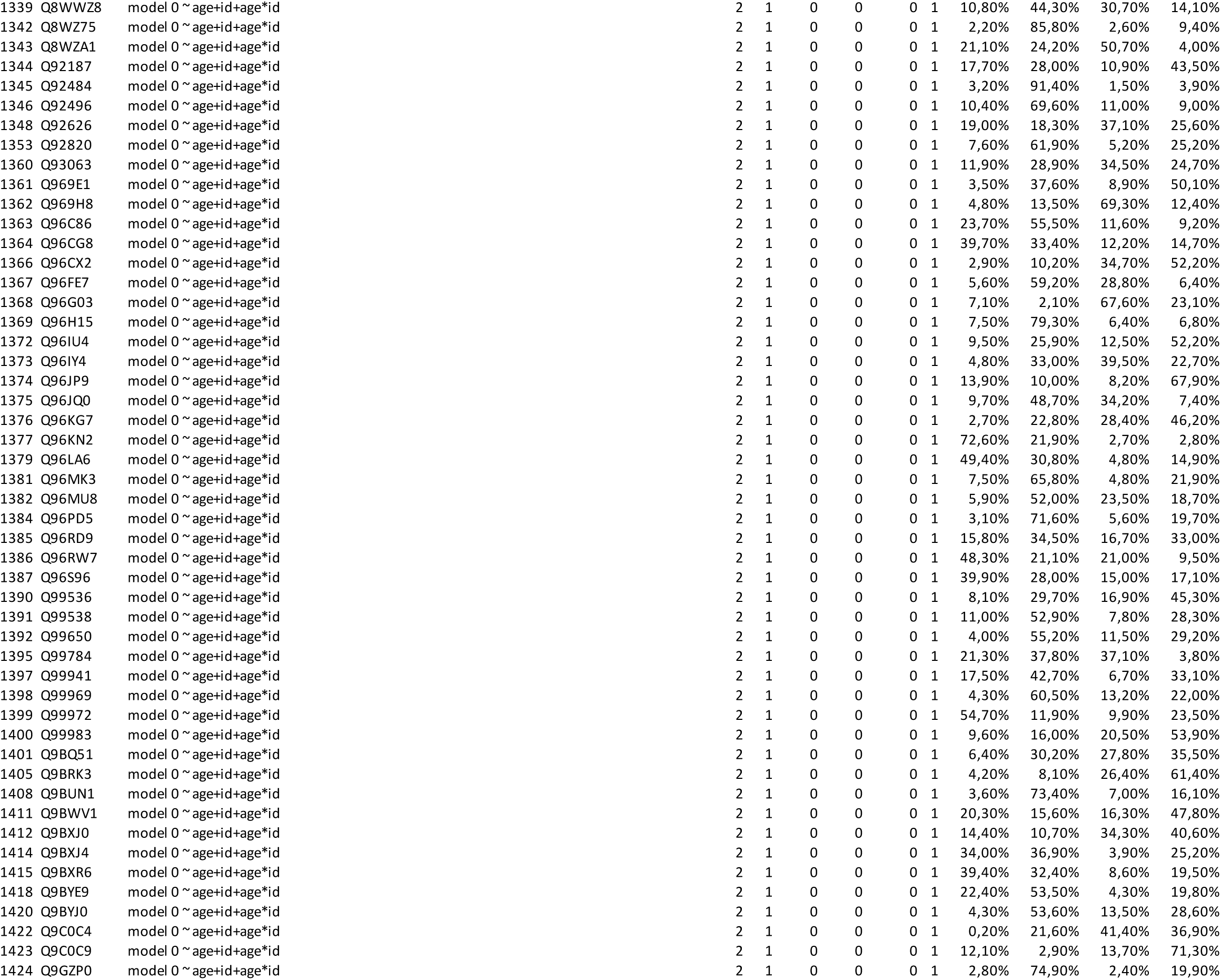
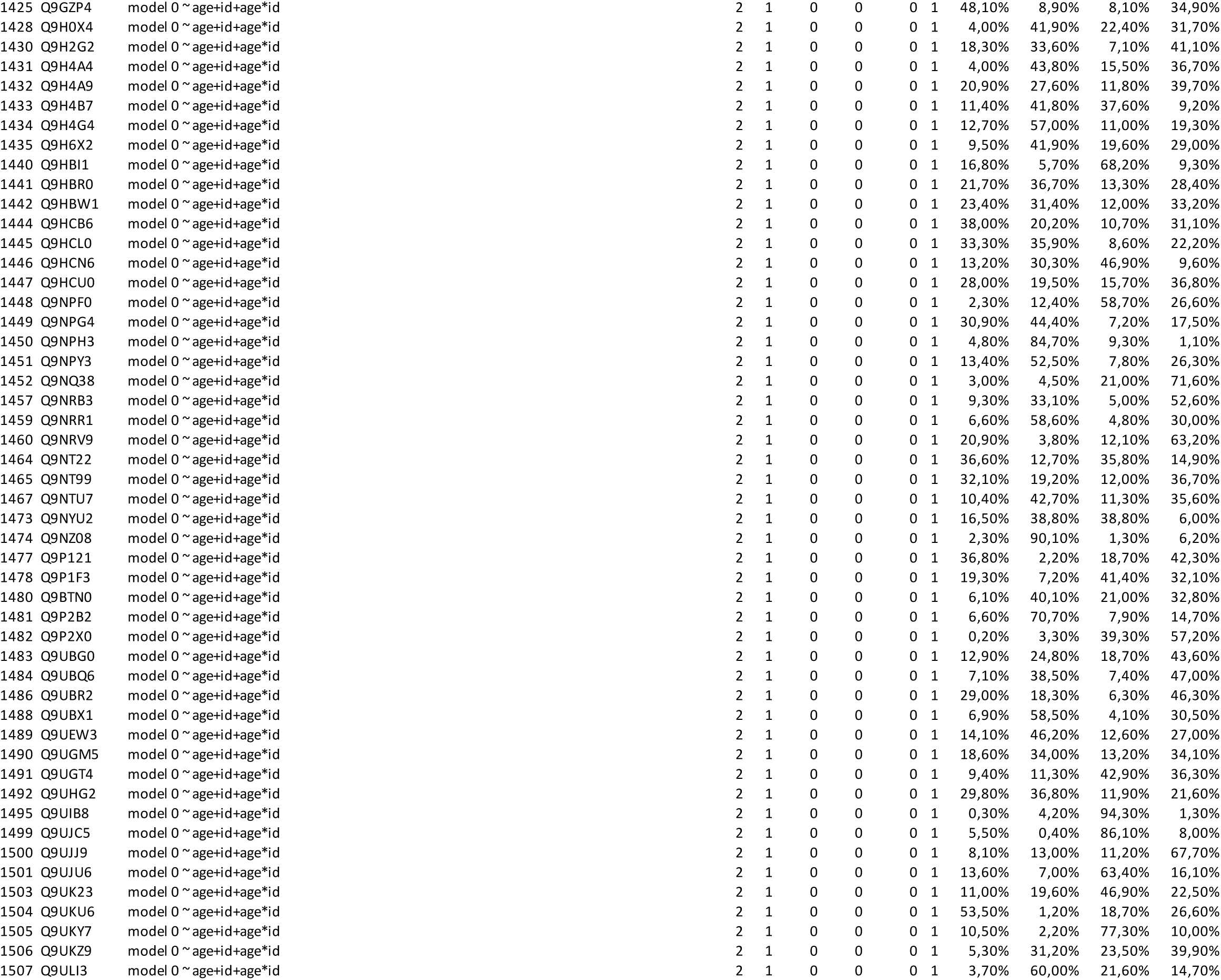
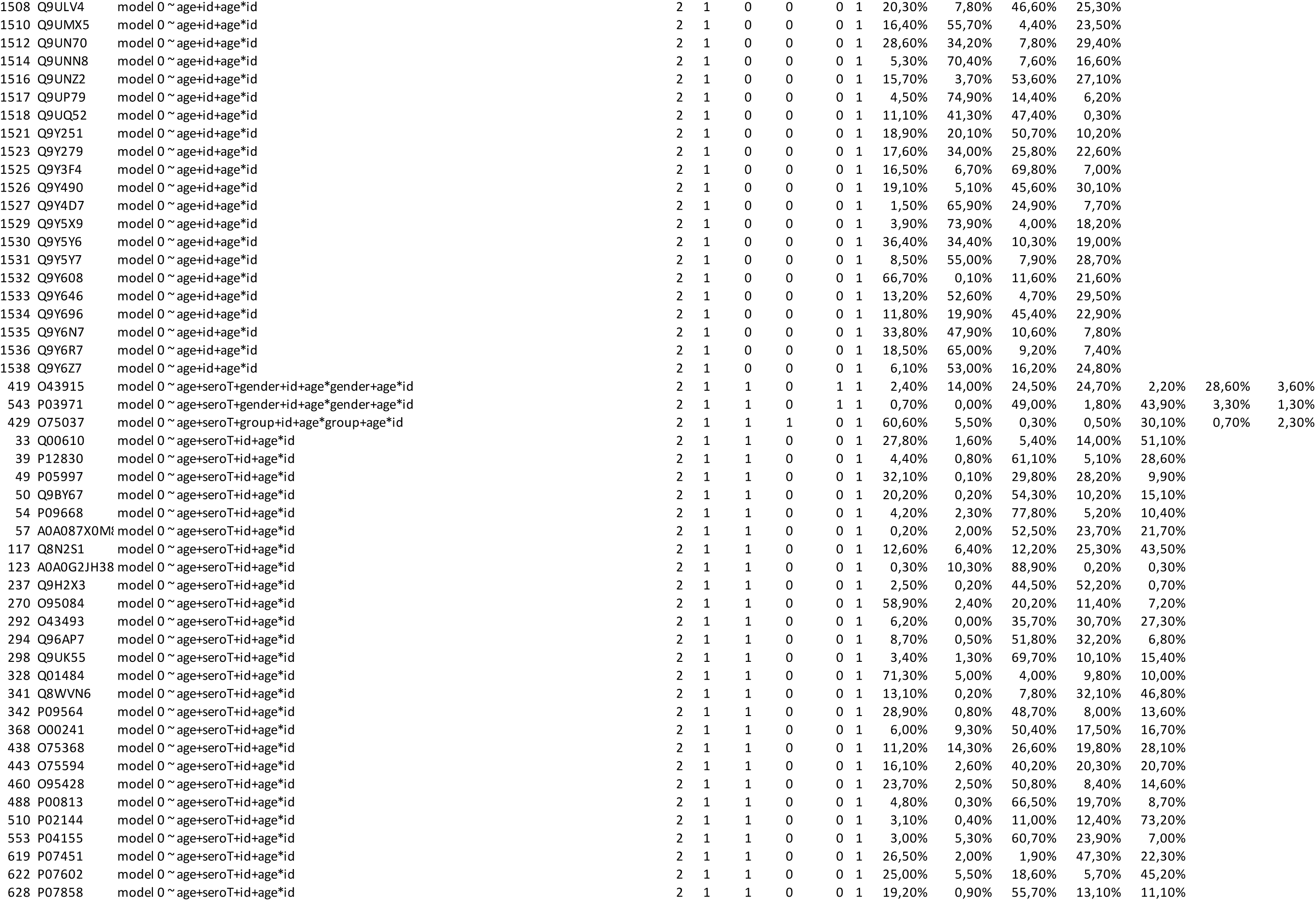
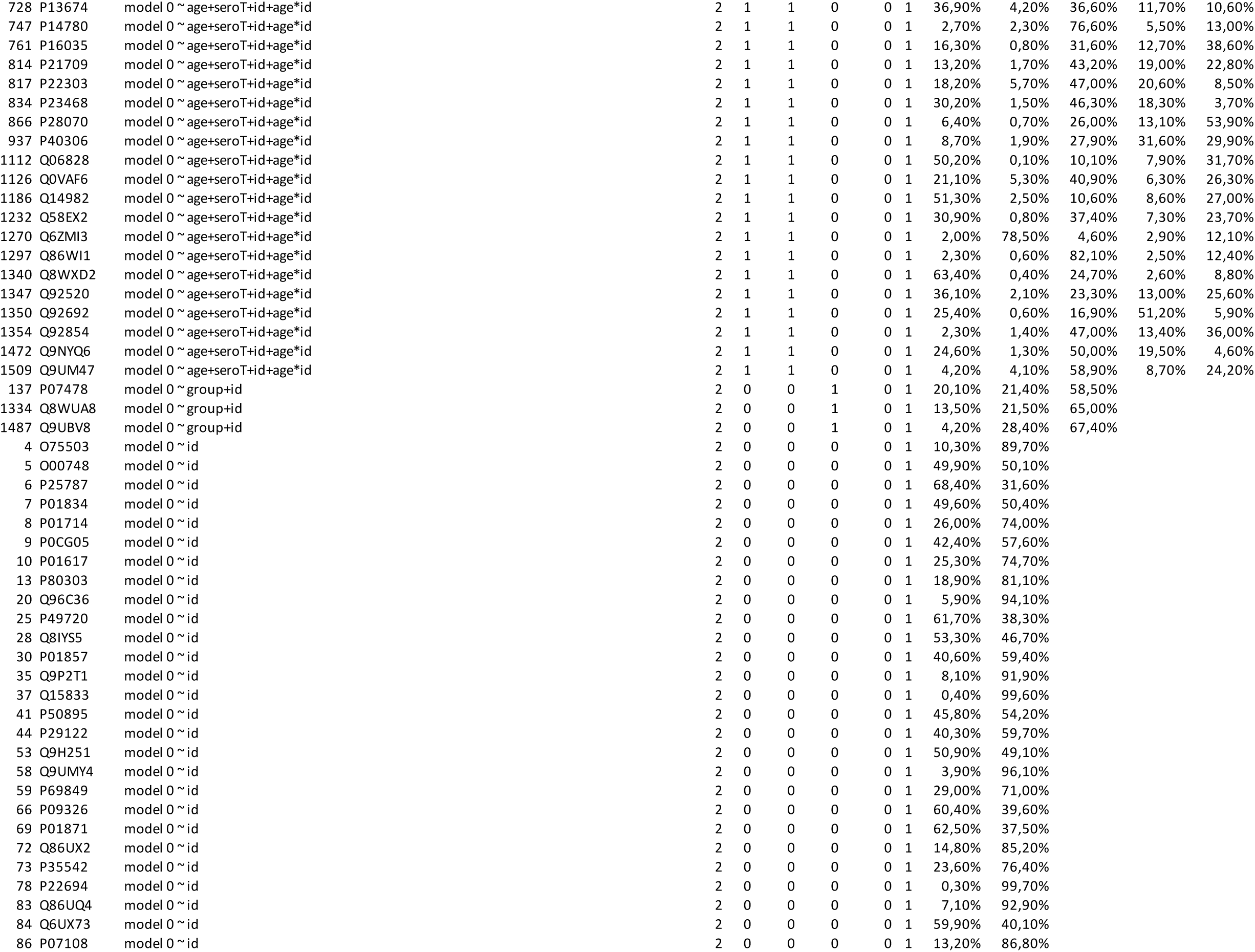
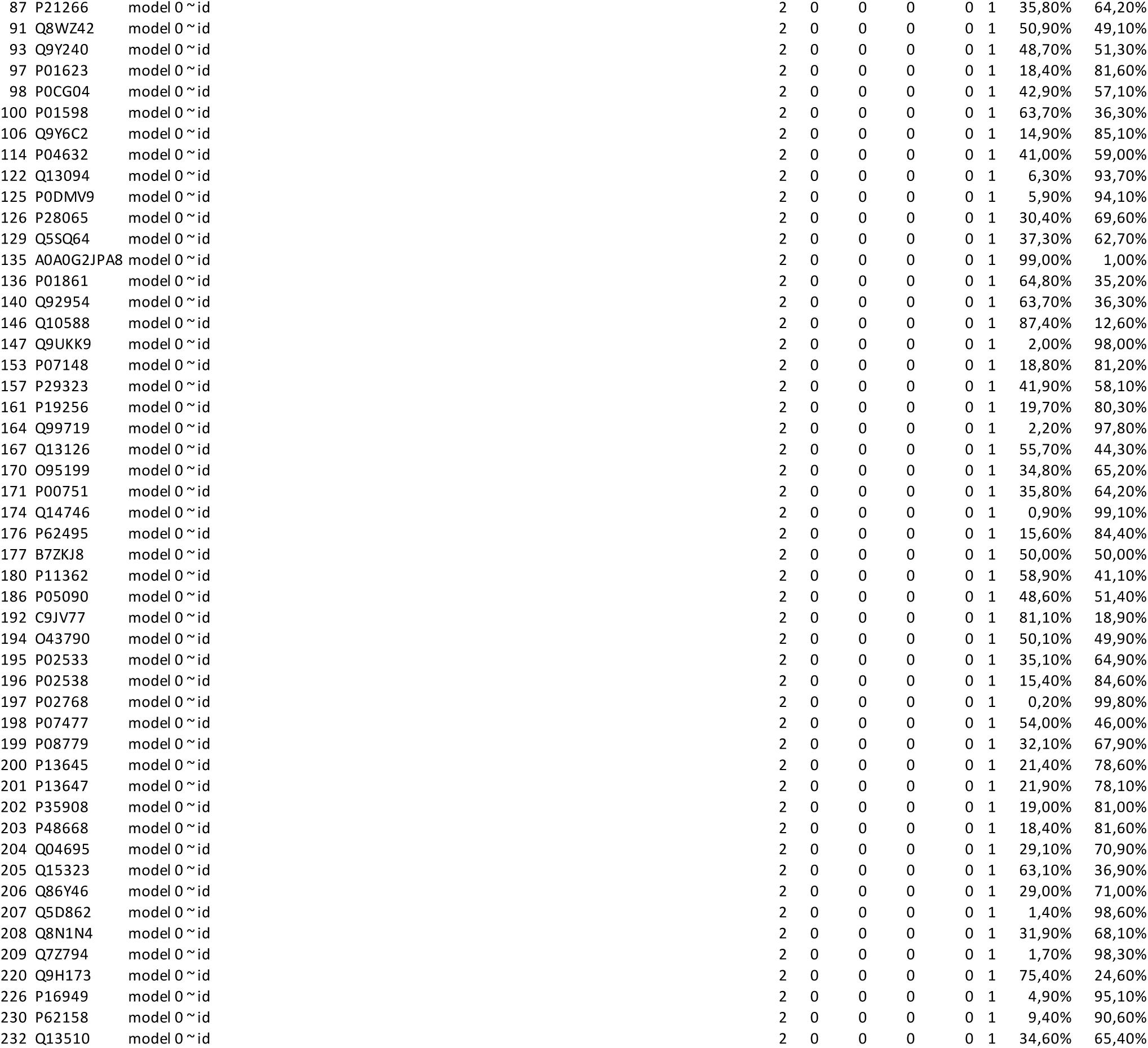
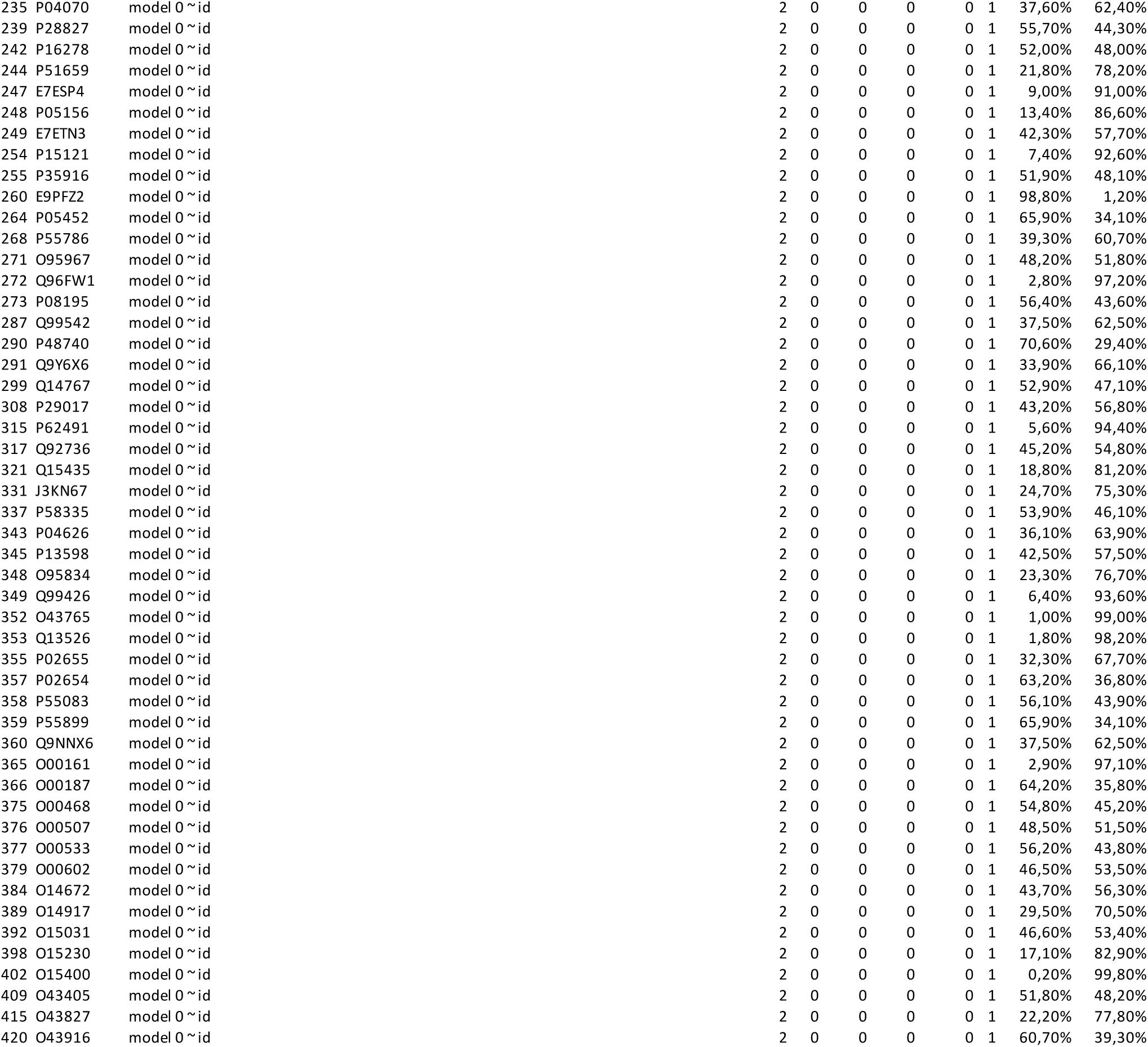
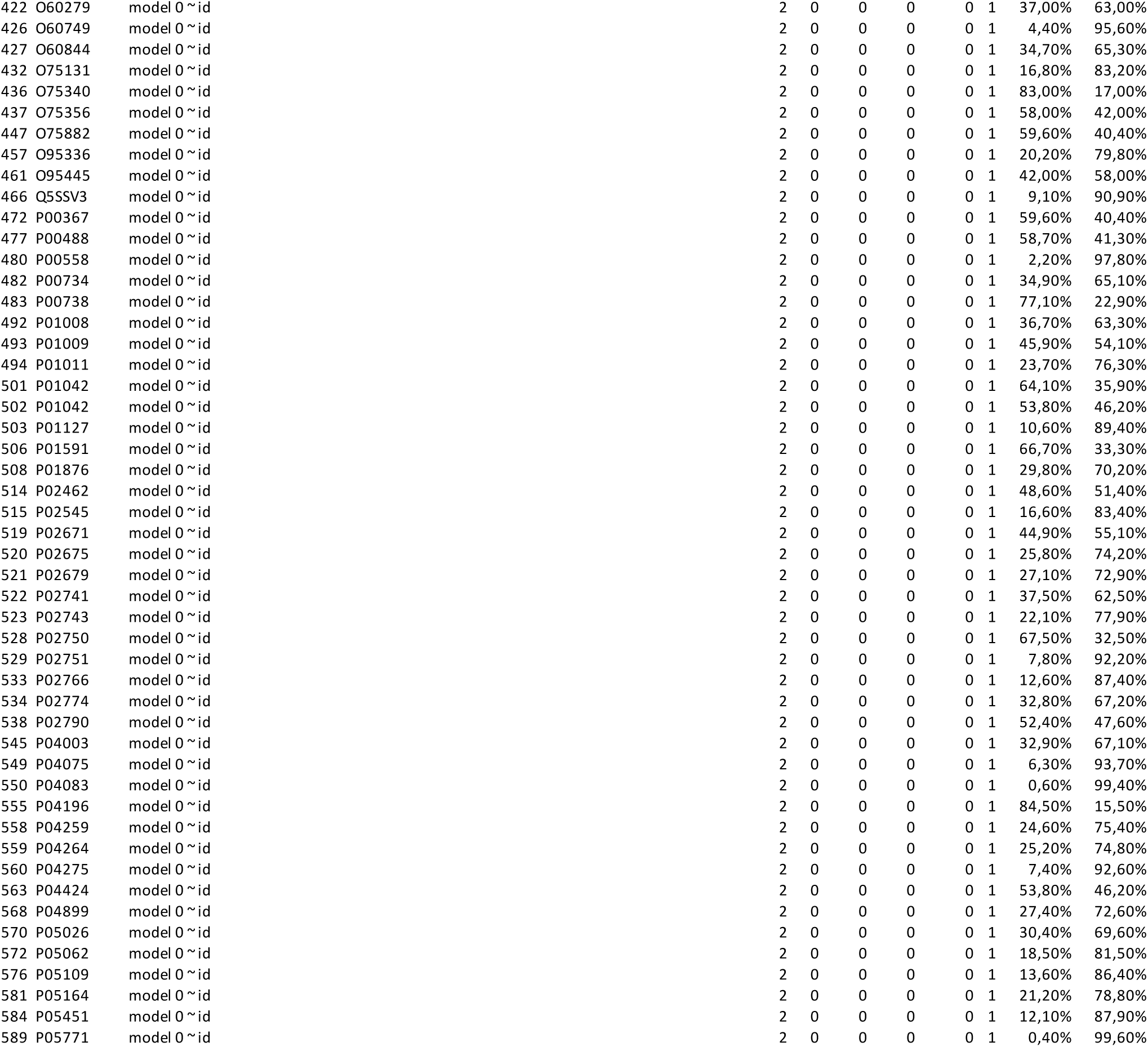
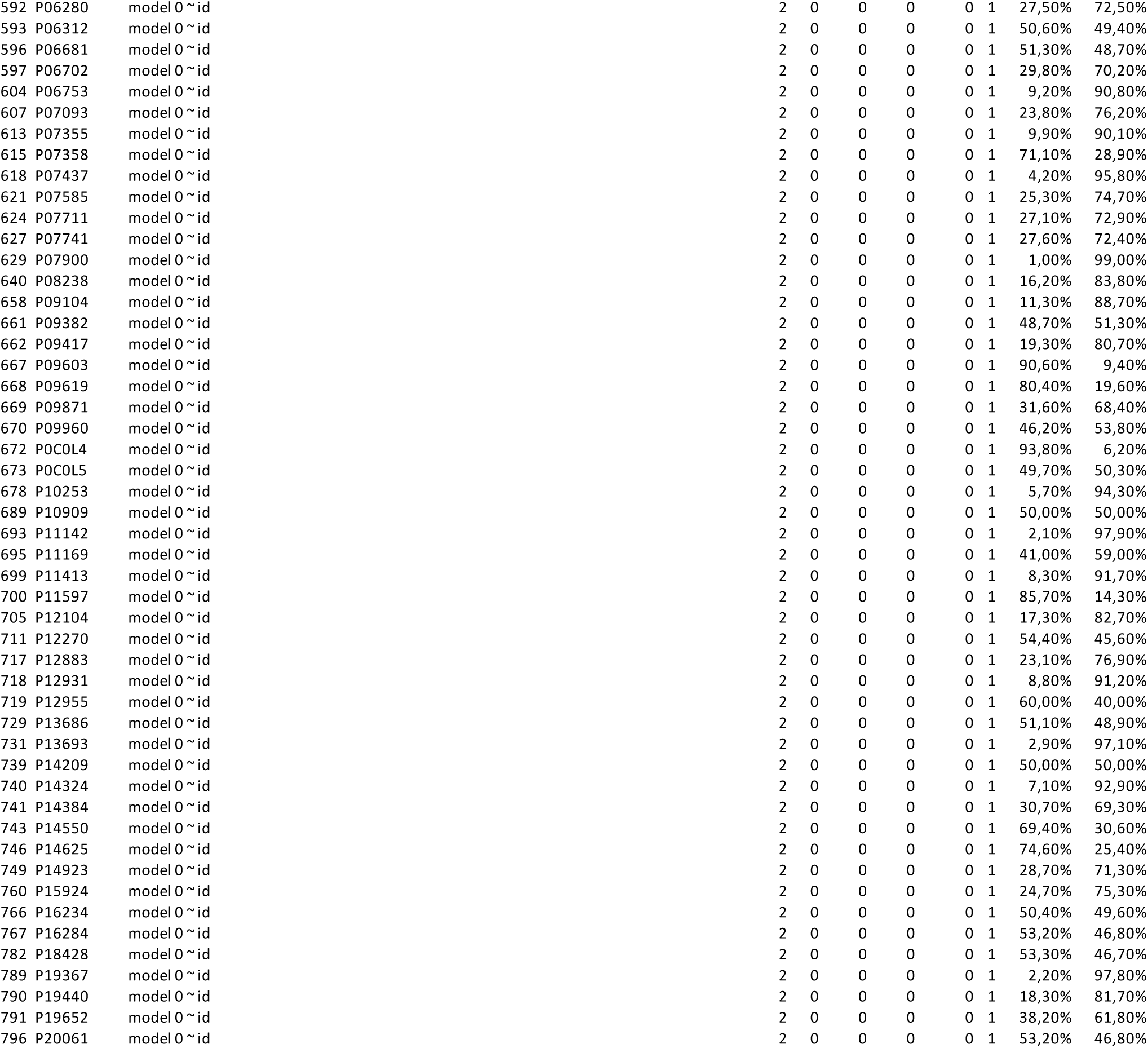
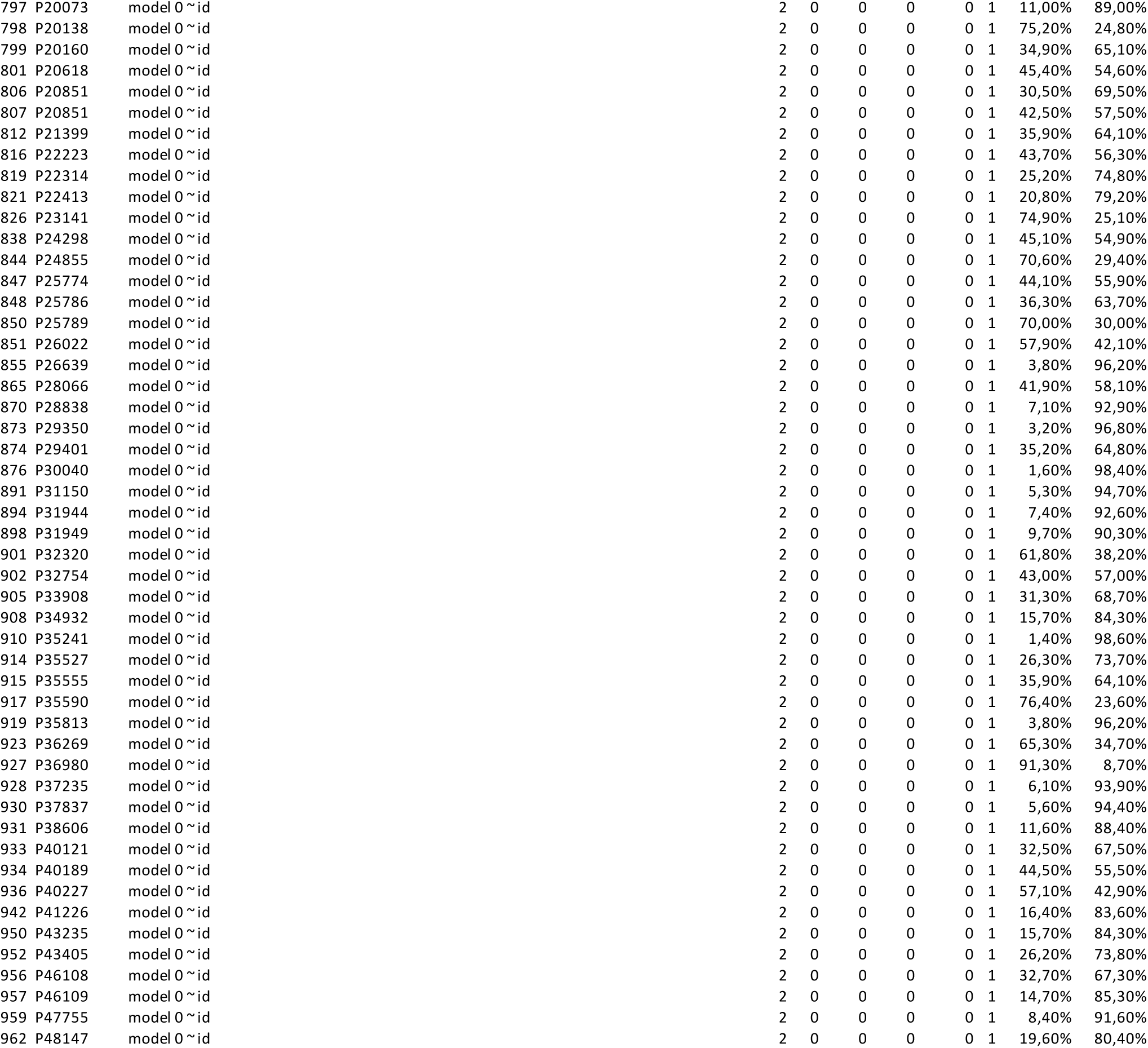
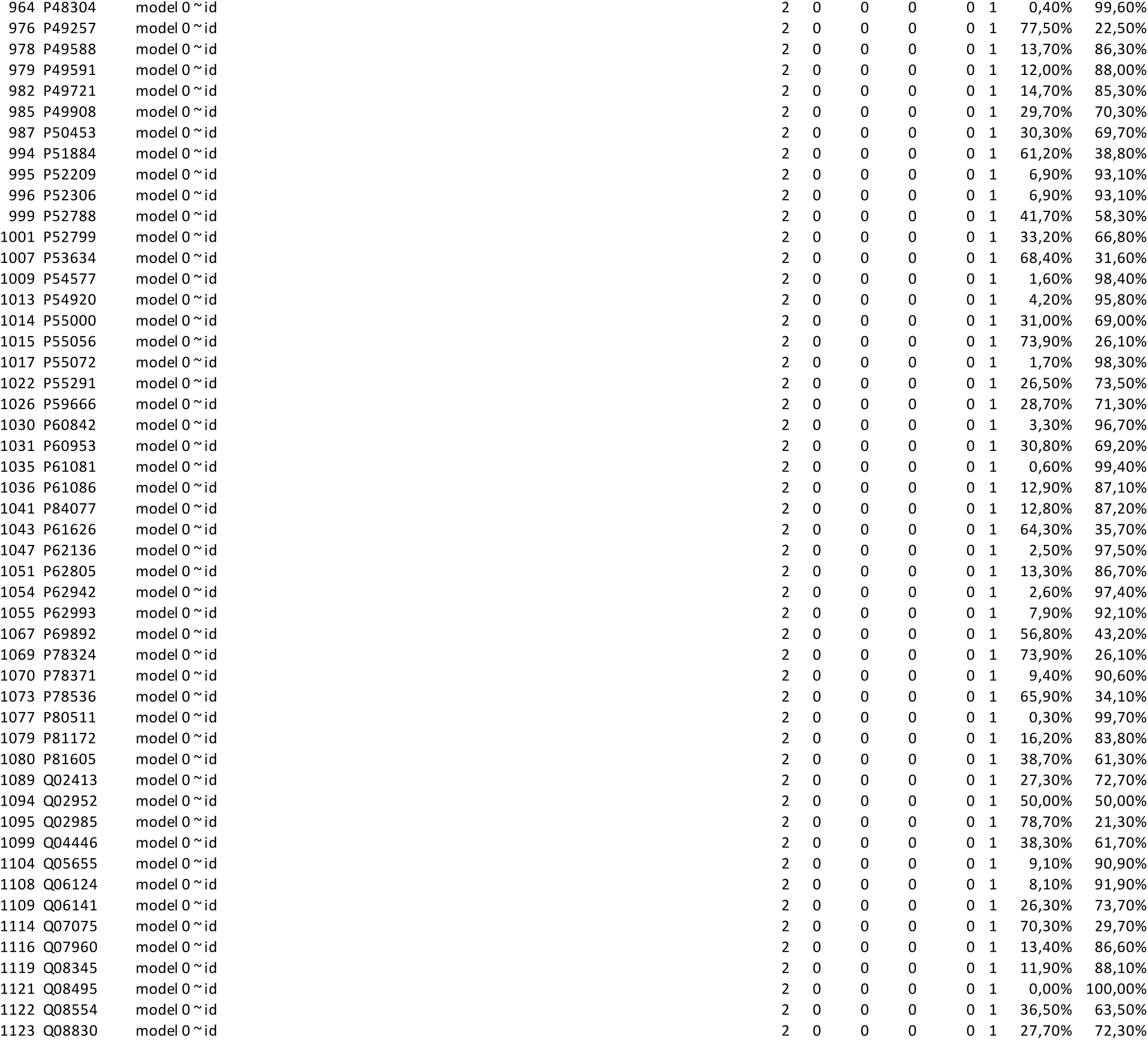
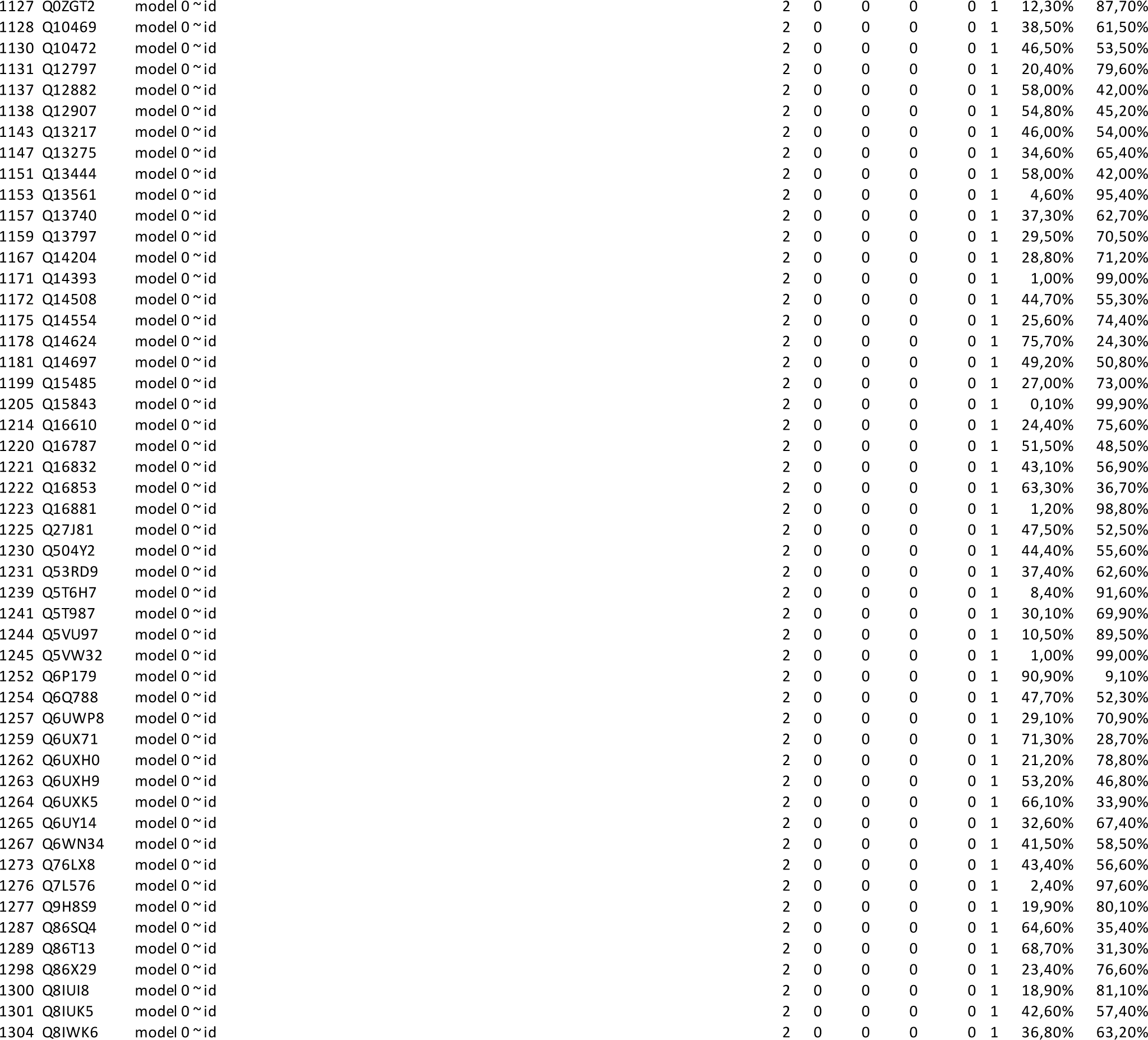
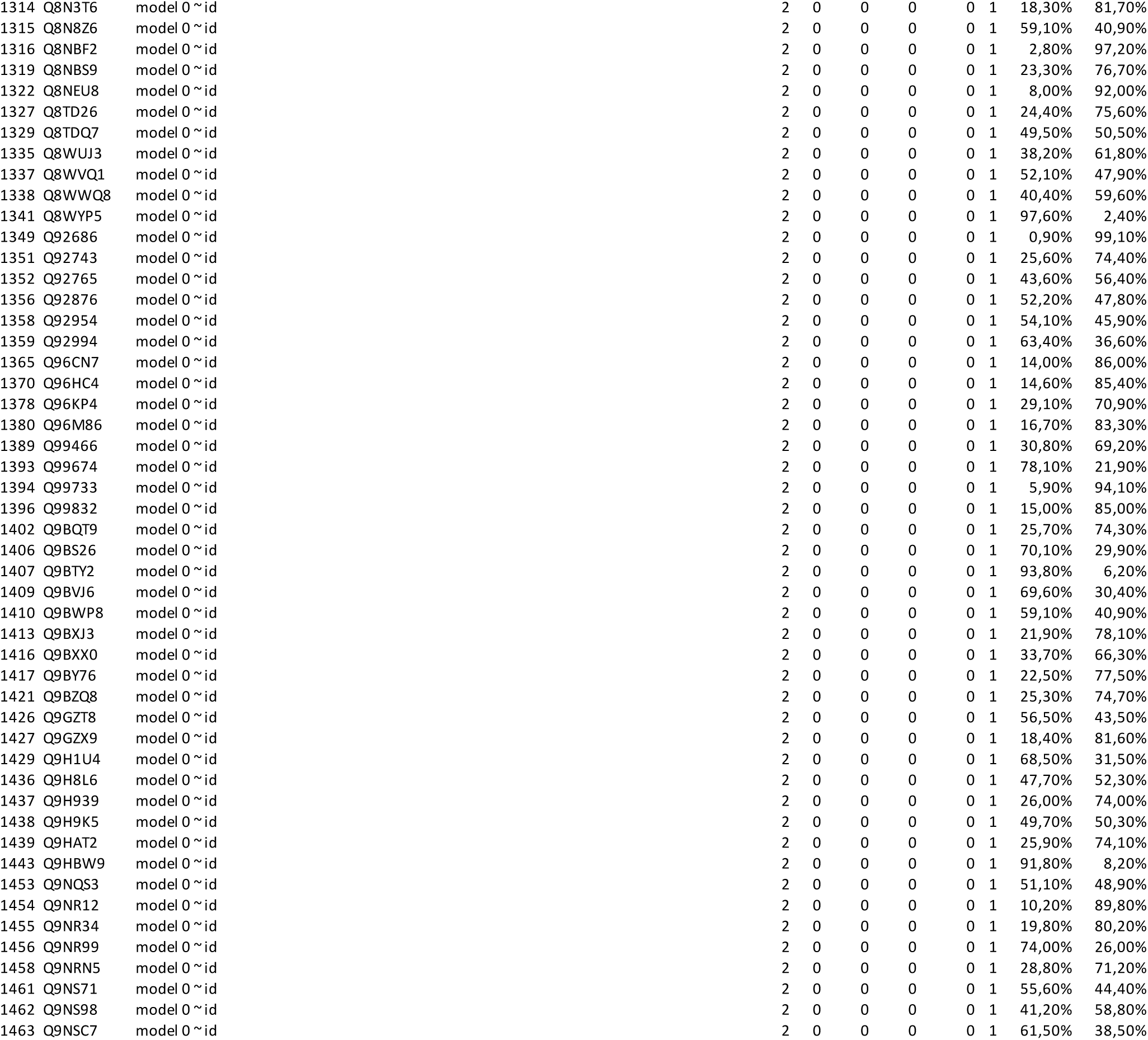
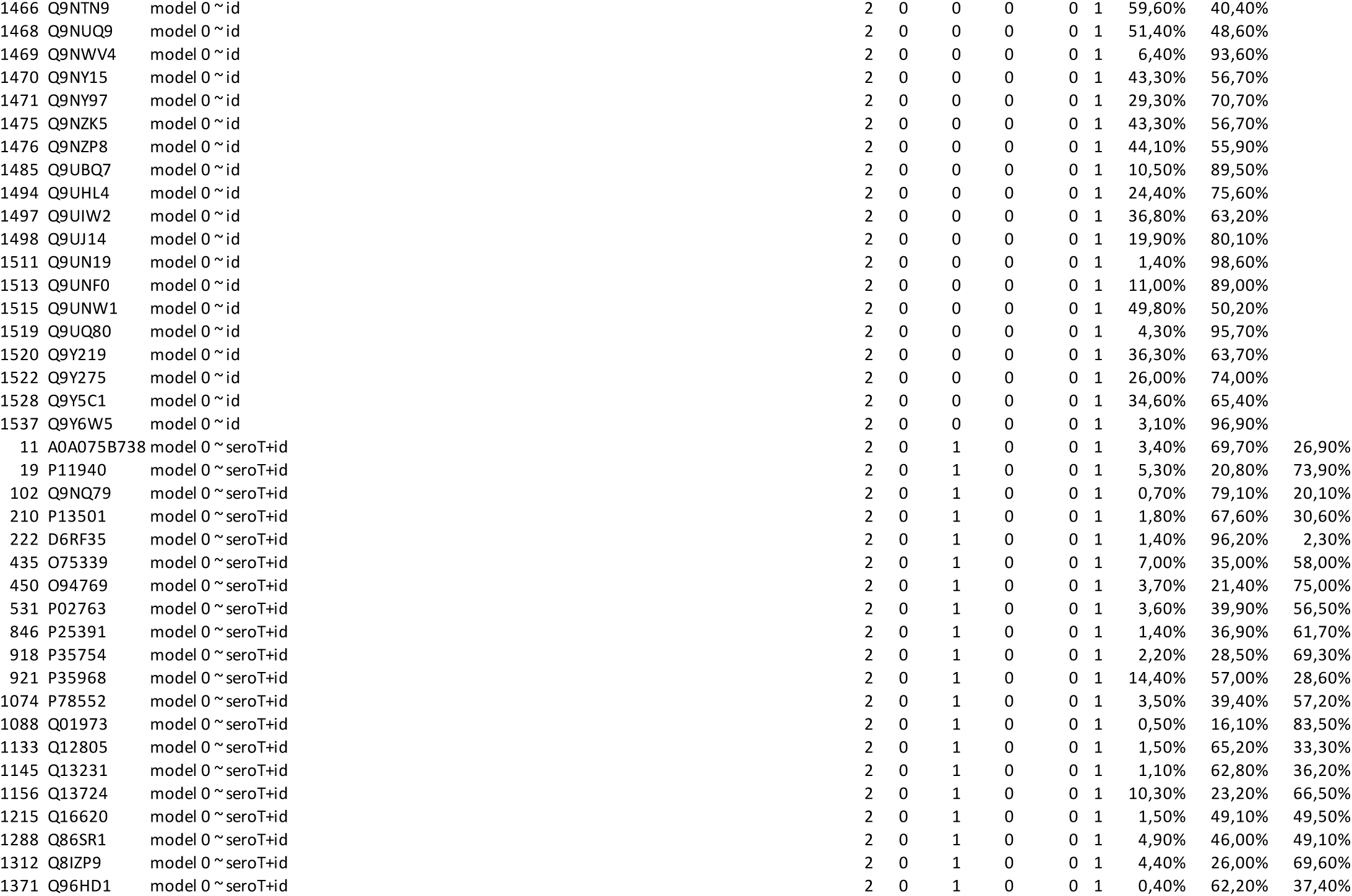
Supplementary File 2

